# Insights into human genetic variation and population history from 929 diverse genomes

**DOI:** 10.1101/674986

**Authors:** Anders Bergström, Shane A. McCarthy, Ruoyun Hui, Mohamed A. Almarri, Qasim Ayub, Petr Danecek, Yuan Chen, Sabine Felkel, Pille Hallast, Jack Kamm, Hélène Blanché, Jean-François Deleuze, Howard Cann, Swapan Mallick, David Reich, Manjinder S. Sandhu, Pontus Skoglund, Aylwyn Scally, Yali Xue, Richard Durbin, Chris Tyler-Smith

## Abstract

Genome sequences from diverse human groups are needed to understand the structure of genetic variation in our species and the history of, and relationships between, different populations. We present 929 high-coverage genome sequences from 54 diverse human populations, 26 of which are physically phased using linked-read sequencing. Analyses of these genomes reveal an excess of previously undocumented private genetic variation in southern and central Africa and in Oceania and the Americas, but an absence of fixed, private variants between major geographical regions. We also find deep and gradual population separations within Africa, contrasting population size histories between hunter-gatherer and agriculturalist groups in the last 10,000 years, a potentially major population growth episode after the peopling of the Americas, and a contrast between single Neanderthal but multiple Denisovan source populations contributing to present-day human populations. We also demonstrate benefits to the study of population relationships of genome sequences over ascertained array genotypes. These genome sequences are freely available as a resource with no access or analysis restrictions.

## Introduction

Genome sequences from diverse human groups can reveal the structure of genetic variation in our species and the history of, and relationships between, different populations, and provide a framework for the design and interpretation of medical-genetic studies. A consensus view of the history of our species includes divergence from the ancestors of the archaic Neanderthal and Denisovan groups 500,000-700,000 years ago, the appearance of anatomical modernity in Africa in the last few hundred thousand years, an expansion out of Africa and the Near East 50,000-70,000 years ago with a reduction in genetic diversity in the descendant populations, admixture with archaic groups in Eurasia shortly after this and large-scale population growth, migration and admixture following multiple independent transitions from hunter-gatherer to food producing lifestyles in the last 10,000 years (*1*). However, much still remains to be understood about the extent to which population histories differed between continents and regions, and how this has shaped the present-day distribution and structure of genetic variation across the species. Large-scale genome sequencing efforts have so far been restricted to large, metropolitan populations and employed low-coverage sequencing (*2*), while those sampling human groups more widely have mostly been limited to 1-3 genomes per population (*3*, *4*). Here, we present 929 high-coverage genome sequences from 54 geographically, linguistically and culturally diverse populations (Fig. 1A) from the Human Genome Diversity Project (HGDP)-CEPH panel (*5*), 142 previously sequenced (*3*, *6*, *7*) and 787 reported for the first time here. We also used linked-read technology (*8*) to physically resolve the haplotype phase of 26 of these genomes from 13 populations (table S1). Several iterations of genetic assays applied to the HGDP-CEPH panel have contributed greatly to the understanding of human genetic variation (*3*, *9*–*14*). We present here high-coverage HGDP-CEPH genome sequences and discuss additional insights that emerge from analysis of them.

**Figure 1:**
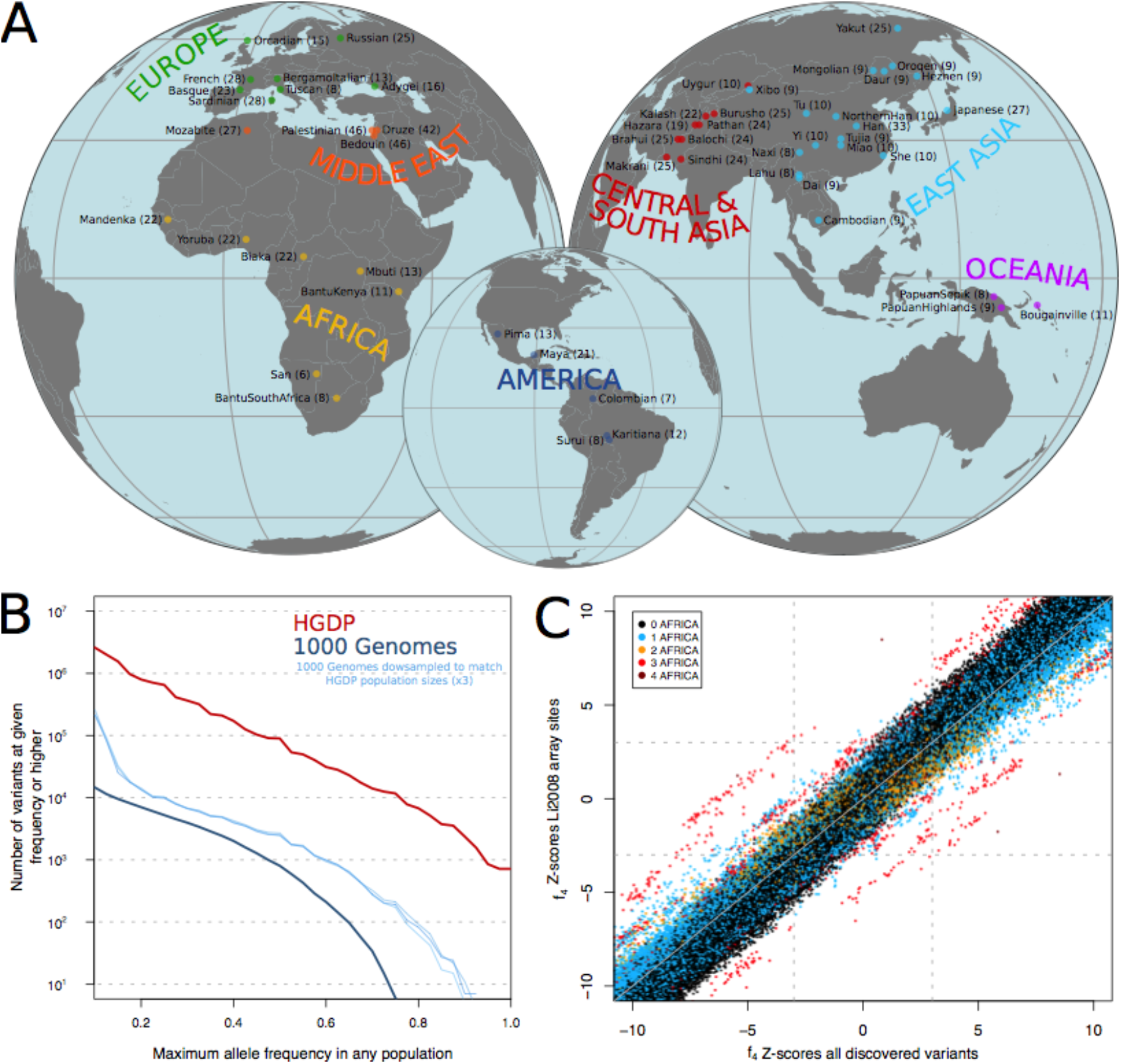
Genome sequencing and variant discovery in 54 diverse human populations. (A) Geographical origins of the 54 populations from the HGDP-CEPH panel, with the number of sequenced individuals from each in parentheses. (B) Maximum allele frequencies of variants discovered in the HGDP dataset but not in the 1000 Genomes phase 3 dataset, and vice versa. The vertical axis displays the number of variants that have a maximum allele frequency in any single population equal to or higher than the corresponding value at the horizontal axis. To account for higher sampling noise due to smaller population sample sizes in the HGDP dataset, results obtained on versions of the 1000 Genomes dataset down-sampled to match the HGDP sizes are also shown. To conservatively avoid counting variants that are actually present in both datasets but not called in one of them for technical reasons, any variant with a global frequency of >30% in a dataset is excluded. (C) Comparison of Z-scores from all possible f4-statistics involving the 54 populations using whole genome sequences and commonly used, ascertained genotyping array sites (*11*). Points are coloured according to the number of African populations included in the statistic.

### Genetic variant discovery across diverse human populations

We performed Illumina sequencing to an average coverage of 35x and mapped reads to the GRCh38 reference assembly. By analysing local sequencing coverage across the genome, we identified and excluded nine samples with large-scale alterations in chromosomal copy numbers that we presume arose during lymphoblastoid cell line culturing. The remaining individuals provided high-quality genotype calls (figs. S1, S2). In this set of 929 genomes we identified 67.3 million single-nucleotide polymorphisms (SNPs), 8.8 million small insertions or deletions (indels) and 39,997 copy number variants (CNVs). This is nearly as many as the 84.7 million SNPs discovered in 2504 individuals by the 1000 Genomes Project (*2*), reflecting increased sensitivity due to high-coverage sequencing as well as the greater diversity of human ancestries covered by the HGDP-CEPH panel. While the vast majority of the variants discovered by one of the studies but not the other are very low in frequency, the HGDP dataset contains substantial numbers of variants that were not identified by the 1000 Genomes Project but are common or even high-frequency in some populations: ~1 million variants at ≥20%, ~100,000 variants at ≥50% and even ~1000 variants fixed at 100% frequency in at least one sampled population (Fig. 1B). This highlights the importance of anthropologically-informed sampling for uncovering human genetic diversity.

The unbiased variant discovery enabled by whole-genome sequencing avoids potential ascertainment biases associated with the pre-defined variant sets used on genotyping arrays. We find that while analyses of the SNPs included on commonly-used arrays accurately recapitulate relationships between non-African populations, they sometimes dramatically distort relationships involving African populations (Fig. 1C). Some of the f_4_-statistics commonly used to study population history and admixture (*13*) even shift sign when using array SNPs compared to when using all discovered SNPs, thus incorrectly reversing the direction of the ancestry relationship one would infer from the same set of genomes (for example: f_4_(BantuKenya, San; Mandenka, Sardinian) is positive (Z=2.9) using all variants but negative (Z=-3.11) when using commonly employed array sites). We demonstrate that 1.3 million SNPs ascertained as polymorphic among three archaic human genomes, mainly reflecting shared ancestral variation (69% of them being polymorphic in Africa), provide more accurate f4-statistics than the variants on commonly used arrays, as well as more accurate F_ST_ values and cleaner estimates of individual ancestries in model-based clustering analyses (fig. S3), consistent with the theoretical properties of outgroup-ascertained variants (*13*).

Rare variants, largely absent from genotyping arrays, are more likely to derive from recent mutation and can therefore be particularly informative about recently shared ancestry between individuals. The patterns of rare variant sharing across the 929 genomes reveal abundant structure (Fig. 2A), as well as a general pattern of greater between-population rare allele sharing among Eurasian as opposed to Oceanian and American populations. We do not find a general increase in the power to detect population relationships in the form of non-zero f_4_ statistics when using all the discovered SNPs, most of which are rare, compared to using just the ~600,000 variants present on commonly used genotyping arrays (Fig. 1C). However, stratifying D-statistics by derived allele frequency can reveal more nuanced views of population relationships (*15*). In the presence of admixture, statistics of the form D(Chimp,X;A,B), quantifying the extent to which the allele frequencies of X are closer to those of A or B, can take different values for variants that have different derived allele frequencies in X. For example, we find that the west African Yoruba have a closer relationship to non-Africans than to the central African Mbuti at high allele frequencies but the opposite relationship at low frequencies (Fig. 2B), suggesting recent gene flow between Mbuti and Yoruba since the divergence of non-Africans. An excess sharing of San with Mandenka relative to Mbuti at low allele frequencies may similarly reflect low amounts of West African-related admixture into San (Fig. 2C) (*16*). The known Denisovan admixture in Oceanian populations manifests itself, without making use of any archaic genome sequences, in a greater affinity of African populations to Eurasians over Oceanians specifically at variants that are fixed in Africans (Fig. 2D). In a manner analogous to this, at fixed variants the central African Biaka have much greater affinity to Yoruba than to the Mandenka, another West African population (Fig. 2E), which would be consistent with Mandenka having some ancestry that is basal to other African ancestries (*17*).

**Figure 2:**
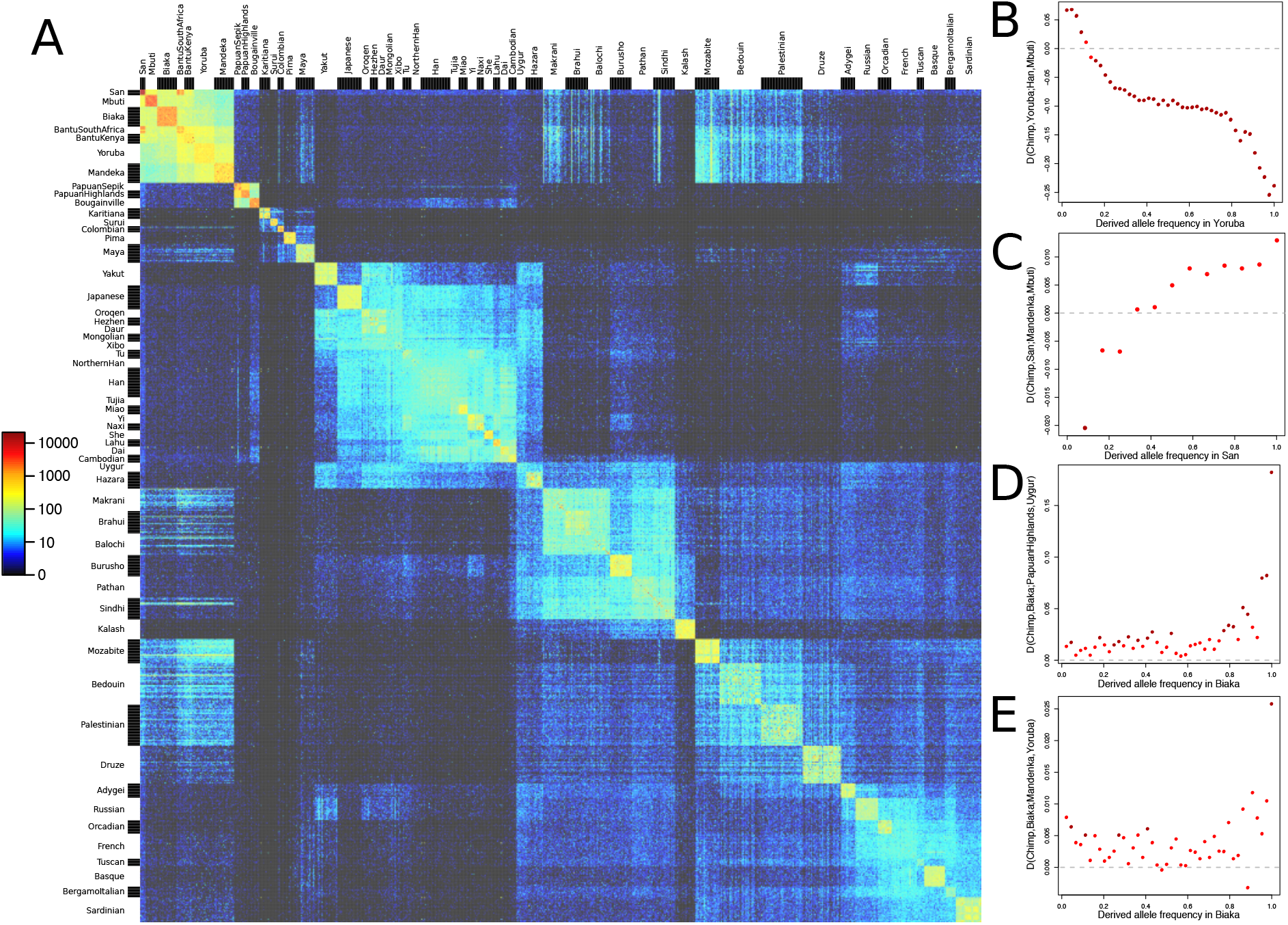
Insights into population relationships from low-frequency variants. (A) A heatmap of pairwise counts of doubleton alleles (alleles observed exactly twice across the dataset) between all 929 individuals, grouped by population. (B-D) D-statistics of the form D(Chimp,X;A,B), stratified by the derived allele frequency in X. Darker red points correspond to |Z| > 3.

The Y chromosome sequences in the dataset recapitulate the well-understood structure of the human Y chromosome phylogeny, but also contain a number of rare lineages of interest (fig. S9). An F* lineage representing the deepest known split in the FT branch that is carried by the vast majority of non-African men was found only once across the 1205 males of the 1000 Genomes Project (*18*). Here, we find it in five out of seven sampled males in the Lahu from Yunnan province in southern China (who also carry high levels of population-specific rare autosomal alleles (Fig. 2A)), pointing to the importance of East Asia for understanding the early dispersal of non-African Y chromosomes, and highlighting how sequencing of diverse human groups can recover genetic lineages that are globally rare.

### The extremes of human genetic differentiation

We next studied the extremes of human genetic variation by identifying variants that are private to geographic regions (excluding individuals with likely recent admixture from other regions). We find no such private variants that are fixed in a given continent or major region (Fig. 3A-C). The highest frequencies are reached by a few tens of variants present at >70% (and a few thousands at >50%) in each of Africa, the Americas and Oceania. In contrast, the highest frequency variants private to either Europe, East Asia, the Middle East or Central and South Asia reach just 10-30%. This likely reflects greater genetic connectivity within Eurasia owing to culturally driven migrations and admixture in the last 10,000 years, events which did not involve the more isolated populations of the Americas and Oceania (*1*), allowing variation accumulating in the latter to remain private. Even comparing Central and South America, we find variants private to one region but absent from the other reaching >40% frequency. Within Africa, ~1000 variants private to the rainforest hunter-gatherer groups Mbuti and Biaka reach >30%, and the highly diverged San of southern Africa harbour ~100,000 private variants at >30% frequency, ~1000 at >60% and even about 20 that are fixed in our small sample of six individuals.

**Figure 3:**
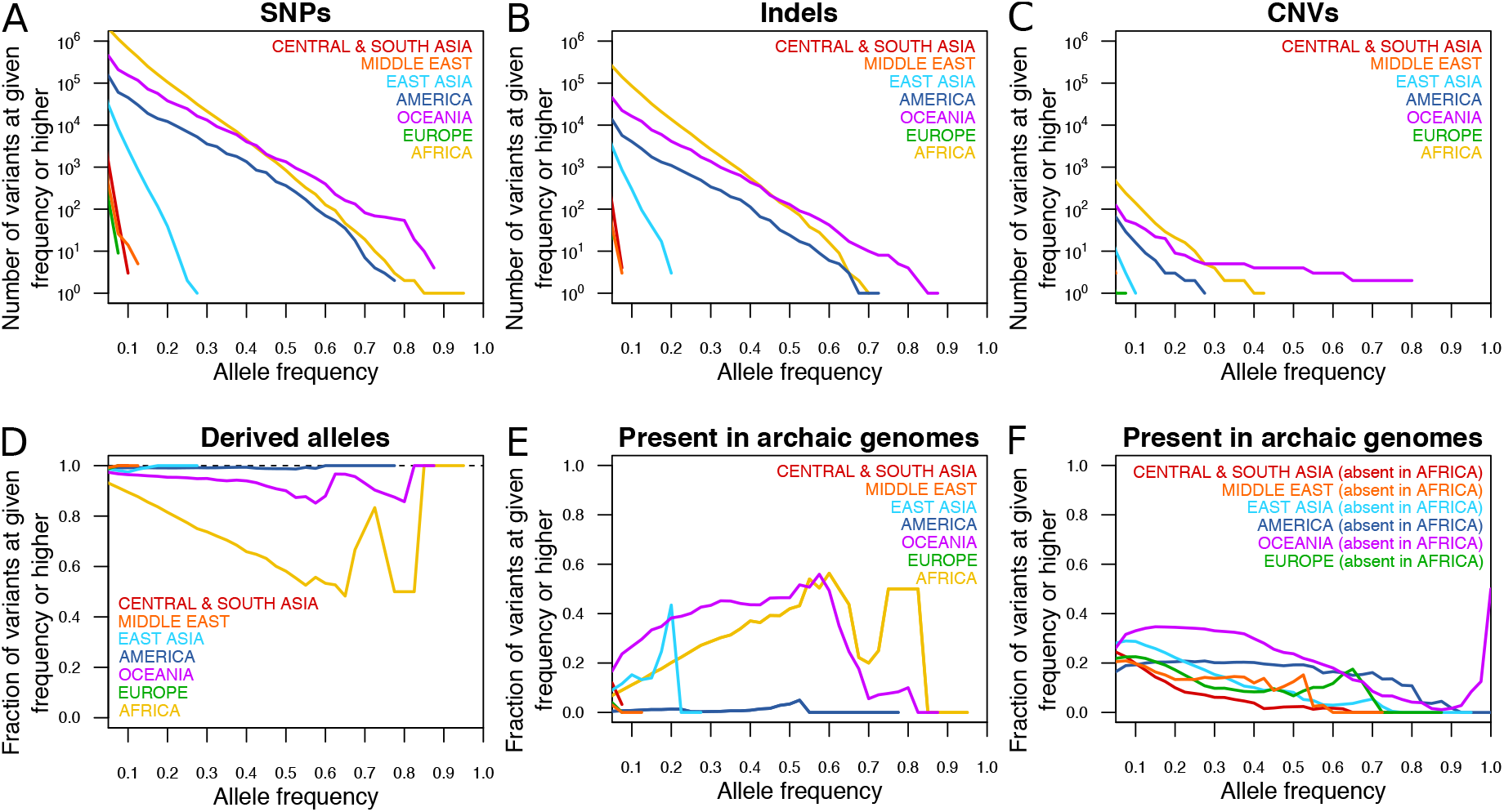
Counts and properties of geographically private variants. (A-C) Counts of region-specific variants. The vertical axis displays the number of variants private to a given geographical region that have an allele frequency in that region equal to or higher than the corresponding value at the horizontal axis. (A) SNPs. (B) Indels. (C) CNVs. (D) The fraction of SNPs private to a given region and at a frequency equal to or higher than the corresponding value on the horizontal axis for which the private allele is the derived as opposed to ancestral state. (E) The fraction of SNPs private to a given region and at a frequency equal to or higher than the corresponding value on the horizontal axis for which the private allele is observed in any of three high-coverage archaic genomes. (F) As E, but counting not variants absent from every other region, but only absent from Africa.

The vast majority of these geographically restricted variants reflect novel mutations that occurred after, or shortly before, the diversification of present-day groups, with >99% of alleles private to most non-African regions being the derived rather than the ancestral allele (Fig. 3D). Alleles private to Africa, however, include a higher proportion of ancestral alleles, and this proportion increases with allele frequency, reflecting old variants that have been lost outside of Africa. For the same reason, many high frequency private African variants are also found in available Neanderthal or Denisovan genomes (*6*, *15*, *19*) (Fig. 3E). The fraction of variants private to any given region outside of Africa that are shared with archaic genomes is very low, consistent with most or all gene flow from these archaic groups having occurred before the diversification of present-day non-African ancestries. The exception to this is Oceania, in which at least ~35% of private variants present at ≥20% frequency are shared with the Denisovan genome. Generally, at least ~20% of common (>10% allele frequency) variants that are present outside of Africa but absent inside Africa are shared with and thus likely derive from admixture with Neanderthals and Denisovans (Fig. 3F). The remaining up to ~80% are more likely to have derived from novel mutations, which thus have been a stronger force than archaic admixture in introducing novel variants into present-day human populations.

Indel variants private to geographic regions display frequency distributions similar to those of SNPs, although reduced in overall numbers by approximately 10-fold (Fig. 3B). The same is mostly true of CNVs, with an even greater reduction in overall numbers, except for a slight excess of high-frequency private CNVs in Oceanians over what would be expected based on the number of private Oceanian SNPs (Fig. 3C, fig. S4). Several of these variants are shared with the available Denisovan genome, suggesting that, relative to other variant classes and geographical regions, positive selection has acted with a disproportionate strength on copy number variants of archaic origin in the history of Oceanian populations.

### Effective population size histories

We next examined what present-day patterns of genetic variation can tell us about the past demographic histories of different human populations. The distribution of coalescence times between chromosomes sampled from the same population can be used to infer changes in effective population size over time (*20*, *21*), but resolution in recent times is limited when analysing single human genomes, and haplotype phasing errors can cause artefacts when using multiple genomes (*22*, *23*). We therefore applied a related method (SMC++) (*23*) which extends this approach to also incorporate information from the site frequency spectrum as estimated from a larger number of unphased genomes, thereby enabling inference of effective population sizes into more recent time periods (Fig. 4A). In Europe and East Asia, most populations are inferred to have experienced major growth in the last 10,000 years, but some geographically or culturally more isolated groups less so, including the European Sardinians, Basques, Orkney islanders, the southern Chinese Lahu and the Siberian Yakut. In Africa, while the sizes of agriculturalist populations increased over the last 10,000 years, those of the hunter-gatherer groups, Biaka, Mbuti and San, saw no growth or even declined. These findings may reflect a more general pattern of human prehistory, in which hunter-gatherer groups which previously might have been more numerous and widespread retracted as agriculturalist groups expanded.

**Figure 4:**
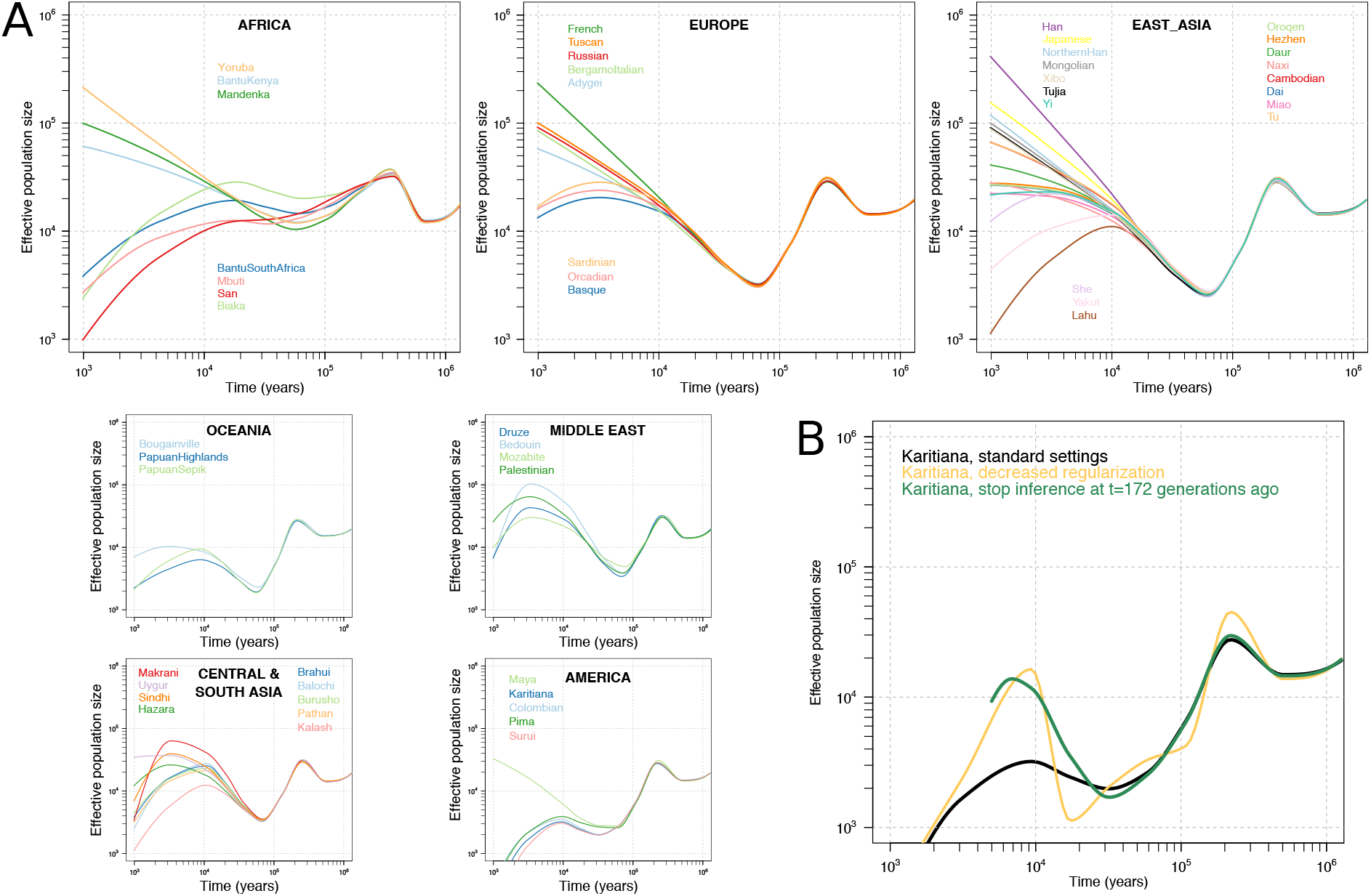
Effective population size histories of 54 diverse populations. (A) Effective population sizes for all populations inferred using SMC++, computed using composite likelihoods across six different distinguished individuals per population. Our ability to infer recent size histories in some South Asian and Middle Eastern populations might be confounded by the effects of recent endogamy. (B) Results for the Native American Karitiana population with varying SMC+ + parameter settings. Decreasing the regularization or excluding the last few thousand years from the time period of inference leads to curves displaying massive growth approximately in the period 10 to 20 kya.

We also find evidence for substantial population growth in the ancestors of Native Americans coinciding with entry into the American continents ~15 kya (Fig. 4B), mirroring observations of rapid diversification of mitochondrial and Y-chromosome lineages at this time (*24*, *25*) but not previously observed using autosomal data. While this inference is sensitive to SMC++ parameter settings and likely counteracted by very recent bottlenecks in the Native American groups, other populations do not display similar histories under these parameter settings. The inferred growth rate exceeds even those of large European and East Asian populations in the last 10,000 years, suggesting this could be one of the most dramatic growth episodes in modern human population history.

While informative, these analyses still appear to have limited resolution to infer more fine-scale population size histories during the transitions to agriculture, metal ages and other cultural processes that have occurred during the last 10,000 years. This might require yet larger sample sizes, novel analytical methods that exploit other features of genetic variation (*26*), or both.

### The time depth and mode of human population separations

We used the 26 genomes physically phased by linked-read technology to study the time course of population separations using the MSMC2 method (*21*, *27*). As a heuristic approximation to the split time between two populations we take the point at which the estimated rate of coalescence between them is half of the rate of coalescence within them, but we also assess how gradual or extended over time the splits were by comparing the shape of the curves to those obtained by running the method on simulated instant split scenarios without subsequent gene flow. Assuming a mutation rate of 1.25 × 10^−8^ per base-pair per generation and a generation time of 29 years, our midpoint estimates suggest (Fig. 5A) splits between the two central African rain forest hunter-gatherer groups Mbuti and Biaka ~62 kya, Mbuti and the west African Yoruba ~69 kya, Yoruba and the southern African San ~126 kya and between San and both of Biaka and Mbuti ~110 kya. Non-Africans have separation midpoints from Yoruba ~76 kya, Biaka ~96 kya, Mbuti ~123 kya and, representing the deepest split in the dataset, from San ~162 kya. However, all of these curves are clearly inconsistent with clean splits, suggesting a picture where genetic separations within Africa were gradual and shaped by ongoing gene flow over tens of thousands of years. For example, there is evidence of gene flow between San and Biaka until at least 50 kya, and between each of Mbuti, Biaka and Yoruba until the present day or as recently as the method can infer.

**Figure 5:**
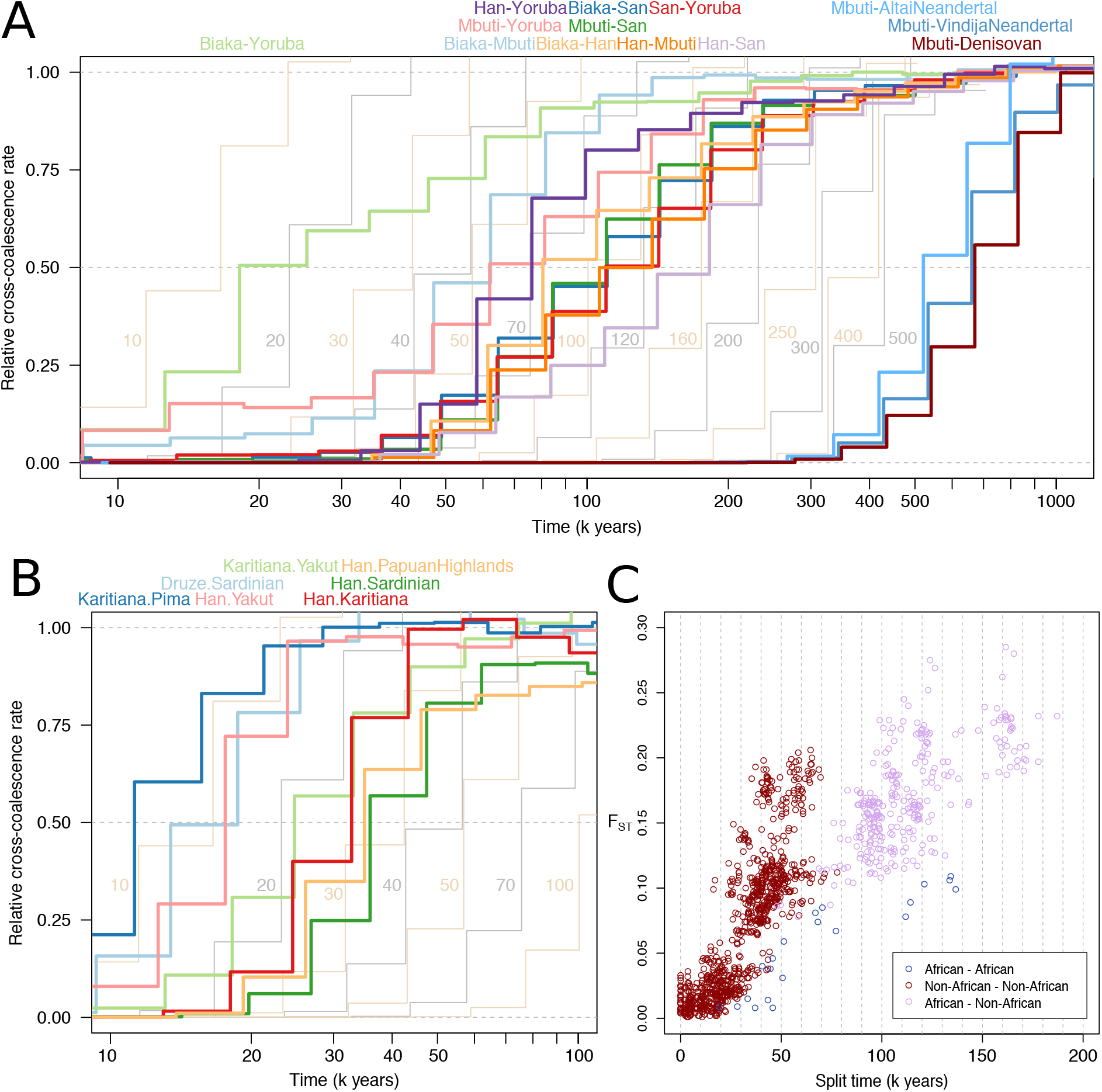
The time depth and mode of population separations. (A) MSMC2 cross-population results for pairs of African populations, including Han Chinese as a representative of non-Africans, as well as between archaic populations and Mbuti as a representative of modern humans. Curves between modern human groups were computed using 4 physically phased haplotypes per population, while curves between modern and archaic groups were computed using 2 haplotypes per population and unphased archaic genomes. The results of simulated histories with instantaneous separations at different time points are displayed in the background in alternating yellow and grey curves. (B) MSMC2 cross-population results, as in A, for pairs of non-African populations. (C) Split times estimated under simple, sudden pairwise split models using momi2 for all possible pairs among the 54 populations against F_ST_, a measure of allele frequency differentiation. The plot does not include Native American populations, as we could not obtain reliable momi2 fits for these.

For the deepest splits, there is some evidence of genetic separation dating back to before 300 or even 500 kya, in the sense that even by that time the rate of coalescence between populations still differs from that within populations. The implication of this would be that there lived populations already at this time which have contributed more to some present-day human ancestries than to others. We find that a small degree of such deep structure in MSMC2 curves might be spuriously caused by batch effects associated with sequencing and genotyping pairs of chromosomes from diploid human samples together, but that such effects are not large enough to fully explain the differences in coalescence rates at these time scales (fig. S6). However, even if this signal reflects actual ancient population structure, its magnitude is such that it would only apply to small fractions of present-day ancestries. An analogy to this is how Neanderthal and Denisovan admixture results in a few percent of non-African ancestries separating from some African ancestries approximately half a million years ago, while most of the ancestry was connected until much more recently. We argue, in the light of such composite ancestries in present-day human populations and the clear deviation of our MSMC2 results from instant split behaviours, that single point estimates are inadequate for describing the timing of early modern human population separations. A more meaningful summary of our results might be that the structure we observe among human populations today formed predominantly during the last 250 kya, but that some small fraction of present-day ancestries retains traces of structure that is older than this, potentially by hundreds of thousands of years.

We also applied MSMC2 to the history of separation between archaic and modern human populations. While the method relies on phased haplotypes, the high degree of homozygosity of Neanderthals and Denisovans means that it might still perform well despite the absence of phase information for heterozygous sites in these genomes. The midpoint estimates suggest that modern and archaic populations separated 550-700 kya (Fig. 5A), in line with, but potentially slightly earlier than, estimates obtained with other methods (*15*, *19*). These results also provide relative constraints on the overall time depth of modern human structure that are independent of the mutation rate we use to scale the results, in the sense that the deepest modern human midpoints are less than one-third of the age of the midpoints of the archaic curves. However, the deep tails of some modern human curves partly overlap a time period when genetic separation from the archaics might still not have been complete. The separation between archaic and modern humans appears more sudden than those between different modern human populations, and only slightly less sudden than expected under an instant split scenario, suggesting a qualitatively different mode of separation between modern and archaic groups than between modern human groups within Africa. While the divergence time between modern human and Neanderthal mitochondrial genomes shows that there is at least some ancestry shared more recently than 500 kya (*28*), these MSMC2 results suggest that post-split gene flow to and from the archaic groups, likely geographically restricted to Eurasia, overall would have been limited.

Outside of Africa, the time depths of population splits are in line with previous estimates (*3*, *4*, *21*), with all populations sharing most of their ancestry within the last 70 kya (Fig. 5B). Our analyses of these physically phased genomes do not replicate a previously observed earlier divergence of West Africans from Oceanians than from Eurasians in MSMC analyses (*4*, *27*), suggesting those results were caused by some artefact of statistical phasing. Instead, all non-African populations display very similar histories of separation from African populations (fig. S5). Like those within Africa, many curves between non-African populations are more gradual than instant split simulations. However, some curves, including those between the Central American Pima and the South American Karitiana, between Han Chinese and the Siberian Yakut, or between the European Sardinians and the Near Eastern Druze, do not deviate appreciably from those expected under instant splits. This suggests that once modern humans had expanded into the geographically diverse and fragmented continents outside of Africa, populations would sometimes separate suddenly and without much subsequent gene flow.

We also fit simple pairwise split models for the complete set of 1431 population pairs to the site-frequency spectrum using momi2 (*29*), obtaining estimates with high concordance to the MSMC2 midpoints (r = 0.93). This much larger set of split time estimates is consistent with present-day populations sharing the majority of their ancestry within the last 200 kya. Using these estimates, we also find that the strength of allele frequency differentiation between populations (FST) relative to split times is about three times greater outside than inside of Africa (Fig. 5C). This could partly reflect increased rates of drift in some non-African populations, but is likely largely explained by the amplifying effects on F_ST_ of the reduced diversity of these groups following their shared bottleneck event (*30*).

### The genetic contribution of archaic hominins to present-day human populations

We estimate an average of 2.4% and 2.1% Neanderthal ancestry in eastern non-Africans and western non-Africans, respectively. We estimate 2.8% (95% confidence interval: 2.1-3.6%) Denisovan ancestry in Papuan highlanders, substantially lower than the first estimate of 4-6% (*31*) based on less comprehensive modern and archaic data, but only slightly lower than more recent estimates (*6*, *32*, *33*). The proportion of ancestry that remains in present-day Oceanian populations after the Denisovan admixture is thus likely not much higher than the amount of Neanderthal ancestry that remains in non-Africans generally.

We identified Neanderthal and Denisovan segments in non-African genomes using a hidden Markov model, and studied the diversity of these haplotypes to learn about the structure of these admixture events and whether they involved one or more source populations. For Neanderthals, several lines of evidence are consistent with there having been a single source with no apparent contribution from any additional population which was detectably different in terms of ancestry, geographical distribution or admixture time. Neanderthal segments recovered from modern genomes across the world show very similar distributions along the genome (fig. S14 and table S6) and profiles of divergence to available archaic genomes (fig. S15), and different Neanderthal haplotypes detected at the same location in modern genomes rarely form geographically structured clusters (fig. S20, table S8). The structure of absolute divergence (D_XY_) in Neanderthal segments between pairs of non-African populations mirrors that in unadmixed segments (Fig. 6A), suggesting a shared admixture event before these populations diverged from each other. A substantial later episode of admixture from Neanderthals into one or more modern populations would have resulted in greater structure (more divergence between some populations) in the Neanderthal segments relative to that in unadmixed segments. Instead, the diversity in unadmixed segments relative to that in Neanderthal segments is higher in western than in eastern non-Africans, perhaps due to gene flow from a source with little or no Neanderthal ancestry into the former (*34*). Although phylogenetic reconstructions indicate that some regions in the genome contain more than 10 different introgressing Neanderthal haplotypes (Fig. 6B, table S7), thus clearly ruling out the scenario of a single contributing Neanderthal individual, the average genetic diversity of admixed Neanderthal sequences is limited (Fig. 6B,C). Coalescent simulations suggest that, genome-wide, as few as 2-4 founding haplotypes are sufficient to produce the observed distribution of haplotype network sizes.

**Figure 6:**
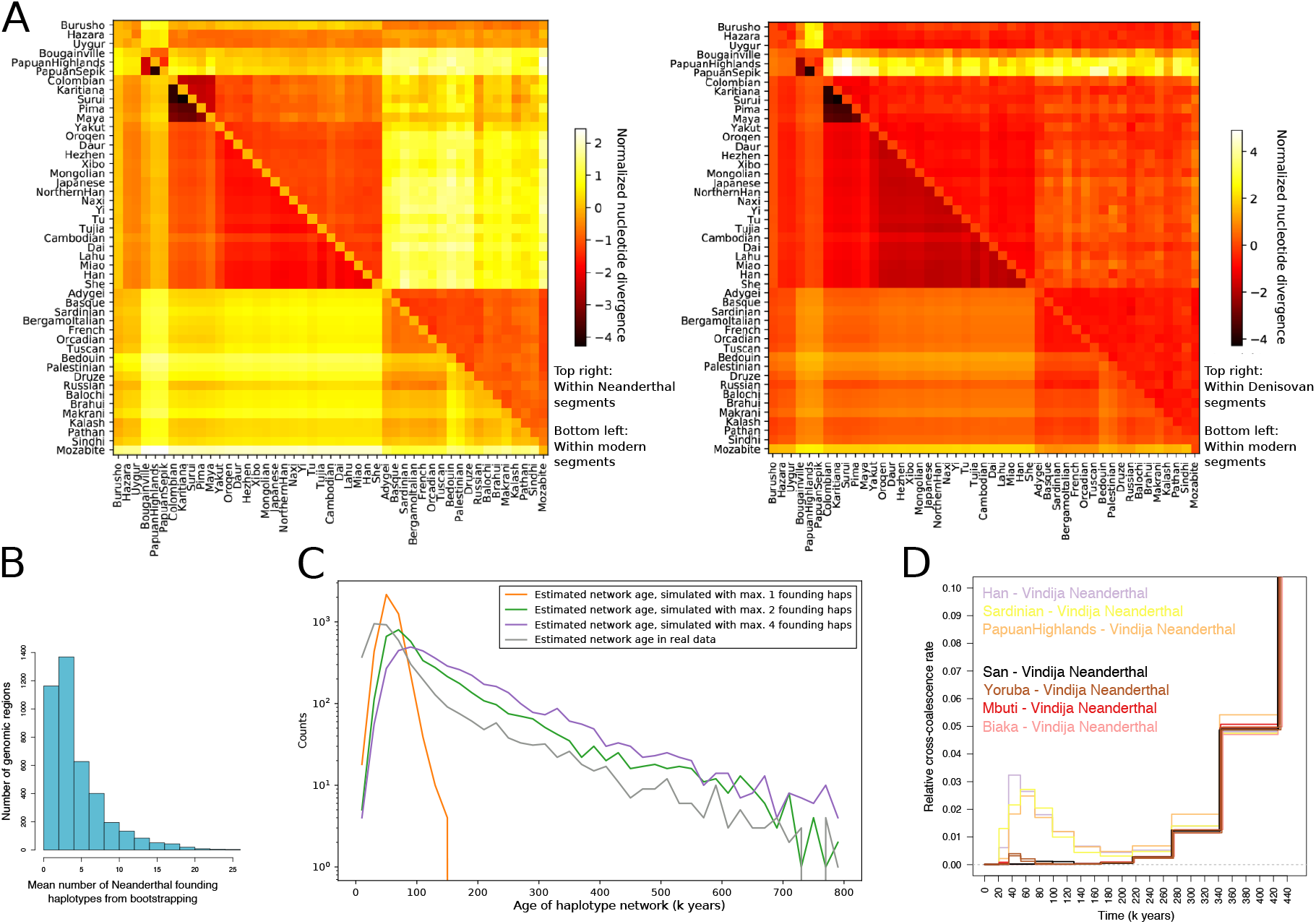
Archaic haplotypes in modern human populations. (A) Nucleotide divergence D_XY_ within segments deriving from archaic admixture and within other segments in non-African populations. (B) The mean number of archaic founding haplotypes estimated by constructing maximum likelihood trees for each archaic segment identified in present-day non-Africans, and then determining the number of ancestral branches in the tree at the approximate time of admixture (2000 generations ago). (C) The distribution of estimated ages of archaic haplotype networks in the present-day human population. The distribution is compared to results obtained in simulations performed with different numbers of archaic founding haplotypes. (D) MSMC2 cross-population results for African (two individual curves per population) and selected non-African (one individual curve per population) against the Vindija Neanderthal, zooming in on the signal of Neanderthal genome flow in modern human genomes (note the highly reduced range of the vertical axis).

In contrast, Denisovan segments show evidence of a more complex admixture history. Segments in Oceania are distinct from those in East Asia, the Americas and South Asia, as shown by their different distribution along the genome (fig. S14 and table S8), high D_XY_ values (Fig. 6A) and a clear separation in most haplotype networks between these two geographical groups (fig. S21, table S8), corresponding to a deep divergence between the Denisovan source populations. East Asian populations also harbour some Denisovan segments that are very similar to the Altai Denisovan genome but which are absent from Oceania (fig. S15). This is consistent with the Denisovan ancestry in Oceania having originated from a separate gene flow event not experienced in other parts of the world (*35*). We do not, however, find clear evidence of more than one source in Oceanians (*36*). The more complicated structure of the Denisovan segments in East Asia (and likely also in America and South Asia) is difficult to explain by one or even two admixture events, and may possibly reflect encounters with multiple Denisovan populations by the ancestors of modern humans in Asia. Some Denisovan haplotypes found in Cambodians are somewhat distinct from those in the rest of East Asia with tentative connections to those in Oceania. Overall, these results paint a picture of an admixture history from Denisovan-related populations into modern humans that is substantially more complex than the history of admixture from Neanderthals.

In MSMC2 analyses, we find that non-Africans display clear modes of non-zero cross-coalescence rates with the Vindija Neanderthal in recent time periods (<100 kya), providing an additional line of evidence for the known admixture episode without requiring assumptions about African populations lacking admixture (Fig. 6D, fig. S7). The Denisovan gene flow into Oceanians is also visible in these analyses but is less pronounced and substantially shifted backwards in time (fig. S7), consistent with the introgressing population being highly diverged from the sequenced individual from the Altai mountains. The West African Yoruba also display a Neanderthal admixture signal, similar in shape but much less pronounced than the signal in non-Africans (Fig. 6D). Other African populations do not clearly display the same behaviour. These results provide evidence for low amounts of Neanderthal ancestry in West Africa, consistent with previous results based on other approaches (*15*, *19*), and we estimate this at 0.18%±0.06% in Yoruba using an f_4_-ratio (assuming Mbuti has none). The most likely source for this is West Eurasian admixture (*37*), and assuming a simple linear relationship to Neanderthal ancestry, our estimate implies 8.6%±3% Eurasian ancestry in Yoruba.

While there is an excess of haplotypes deriving from archaic admixture in non-Africans, many single variants present in archaic populations are also present in Africans due to their having segregated in the population ancestral to archaic and modern humans, and some of these variants were subsequently lost in non-Africans due to increased genetic drift. Counting how many of the variants carried in heterozygote state in archaic individuals are segregating in balanced sets of African and non-African genomes, we find that more Vindija Neanderthal variants survive in non-Africans than in Africans (31.0% vs 26.4%). However, more Denisovan variants survive in Africans (18.9% vs 20.3%). These numbers might change if larger numbers of Oceanian populations were surveyed, but they highlight how the high levels of genetic diversity in African populations mean that, despite having received much less or no Neanderthal and Denisovan admixture, they still retain a substantial, and only partly overlapping (Fig. 3E), subset of the variants which were segregating in late archaic populations.

## Discussion

While the number of human genomes sequenced as part of medically-motivated genetic studies is rapidly growing into the hundreds of thousands, the number resulting from anthropologically-informed sampling to characterize human diversity still remains in the hundreds to low thousands. With the set of 929 genomes from 54 diverse human populations presented here, we greatly extend the number of high-coverage genomes freely available to the research community as part of human global diversity datasets, and substantially expand the catalogue of genetic variation to many underrepresented ancestries. Our analyses of these genomes highlight several aspects of human genetic diversity and history, including the extent and source of geographically restricted variants in different parts of the world, the time depth of separation and extensive gene flow between populations in Africa, a potentially dramatic population expansion following entry into the Americas and a simple pattern of Neanderthal admixture contrasting with a more complex pattern of Denisovan admixture.

One aim of the 1000 Genomes Project (*2*) was to capture most common human genetic variation, which it achieved in the populations included in the study. However, the more diverse HGDP dataset reveals that there are several human ancestries for which this aim was not achieved, and which harbour substantial amounts of genetic variation, some of it common, that so far has been documented poorly or not at all. This is particularly true of Africa and the ancestries represented by the southern African San, and central African Mbuti and Biaka groups. Outside of Africa, Oceanian populations represent one of the major lineages of non-African ancestries and have substantial amounts of private variation, some of it deriving from Denisovan admixture. Any biomedical implications of variants common in these populations but rare or absent elsewhere are unknown, and will remain unknown until genetic association studies are extended to include these and other currently underrepresented ancestries.

Our analyses demonstrate the value of generating multiple high-coverage whole-genome sequences to characterise variation in a population, compared to genotyping using arrays, sequencing to low-coverage or sequencing just small numbers of genomes. In particular, such an approach enables unbiased variant discovery, including of large numbers of low-frequency variants, and higher resolution assessments of allele frequencies. The experimental phasing of haplotypes using linked-read technology aids analyses of deep human population history and structural variation, and is now becoming a feasible alternative to statistical phasing, especially useful in diverse populations. However, short read sequencing still imposes limitations on the ability to identify more complex structural variation. We expect the application of long-read or linked-read sequencing technologies to large sets of diverse human genomes, combined with de-novo assembly or variation graph (*38*) approaches that are less reliant on the human reference assembly, to unveil these additional layers of human genetic diversity.

While the HGDP genome dataset substantially expands our genomic record of human diversity, it too contains considerable gaps in its geographical, linguistic and cultural coverage. We therefore argue for the importance of continued sequencing of diverse human genomes. Given the scale of ongoing medical and national genome projects, producing high-coverage genome sequences for at least ten individuals from each of the approximately 7000 (*39*) human linguistic groups would now arguably not be an overly ambitious goal for the human genomics community. Such an achievement would represent a scientifically and culturally important step towards diversity and inclusion in human genomics research.

## Data availability

Raw read alignments are available from the European Nucleotide Archive under study accession PRJEB6463. Processed per-sample read alignment files are made available by the International Genome Sample Resource at the European Bioinformatics Institute (EMBL-EBI) (http://www.internationalgenome.org/). The 10x Genomics sequencing data generated for 26 samples are available at the European Nucleotide Archive under study accession PRJEB14173. Genotype calls and other downstream analysis files are available from the Wellcome Sanger Institute (ftp://ngs.sanger.ac.uk/production/hgdp). DNA extracts from the samples in the HGDP-CEPH collection can be obtained from the CEPH Biobank at Fondation Jean Dausset-CEPH in Paris, France (http://www.cephb.fr/en/hgdp_panel.php).

## Acknowledgments

We thank the sample donors who made this research possible, as well as the CEPH Biobank, Paris, France (BIORESOURCES) at Fondation Jean Dausset-CEPH, for maintaining the cell line resource and distributing DNA. We thank the Wellcome Sanger Institute sequencing facility for generating data, and Susan Fairley and colleagues at the International Genome Sample Resource for incorporating and hosting data. We thank Jonathan Terhorst, Stephan Schiffels, Robert Handsaker, Deepti Gurdasani and members of the Tyler-Smith and Durbin groups for useful advice and discussions. A.B., S.A.M., M.A.A, Q.A., P.D., Y.C., S.F., P.H., J.K, M.S.S., Y.X., R.D. and C.T.-S. were supported by Wellcome grants 098051 and 206194, and S.A.M. and R.D. also by Wellcome grant 207492. A.B. and P.S. were supported by the Francis Crick Institute (FC001595) which receives its core funding from Cancer Research UK, the UK Medical Research Council and the Wellcome Trust. R.H. was supported by a Gates Cambridge scholarship. P.H. was supported by Estonian Research Council Grant PUT1036. D.R. is an Investigator of the Howard Hughes Medical Institute.

