## Supplementary material for "Insights into human genetic variation and population history from 929 diverse genomes": Materials and Methods

**This PDF file includes:**
Materials and Methods
Figs. S1 to S23
Tables S1 to S9

### Materials and Methods

#### Sequencing and data processing

*Sequencing and read processing*

DNA extracted from lymphoblastoid cell lines was shipped from the CEPH-Biobank at Fondation Jean Dausset-CEPH laboratory in Paris, and sequenced at the Wellcome Sanger Institute in Hinxton, United Kingdom. PCR libraries were constructed for an initial batch of samples (PCR-free library construction was not available for large-scale production at the institute at this time), while PCR-free libraries were constructed for the rest. Libraries were sequenced on Illumina HiSeq X machines, on single lanes for the PCR libraries and multiplexed across 12 lanes for the PCR-free libraries, producing paired-end reads of length 2×151 base-pairs (bp), a mean insert size of 447 bp and a mean coverage of 35.0X. 14 of these libraries were described and used for population genetic analyses in a previous publication (*7*).

Reads were processed through the automated pipeline of the Wellcome Sanger Institute sequencing facility, mapping to the GRCh38 reference assembly (GRCh38_full_analysis_set_plus_decoy_hla.fa) using bwa mem version 0.7.12 (*40*) with the –T 0 parameter. Tools from the biobambam package (*41*) were used to trim adaptor sequences from reads prior to mapping (bamadapterclip) and to mark duplicate reads after mapping (bamstreamingmarkduplicates). We performed no further post-processing of the alignments, e.g. base quality recalibration or local indel realignment.

Data previously generated on a subset of the samples from the HGDP-CEPH panel as part of the Simons Genome Diversity Project (*3*) (hereafter referred to as "SGDP") was also incorporated into the project (paired-end reads of length 2×100 bp, mean insert size of 310 bp, mean coverage of 42.4x). Data from another earlier publication (*6*) (hereafter referred to as "Meyer") was obtained from ENA experiment accession number SRX103808, de-multiplexed allowing for one nucleotide mismatch in the barcode sequence and then also incorporated (paired-end read lengths of 94+100 bp or 95+101 bp, mean insert size of 264 bp, mean coverage of 28.7).

Reads from the SGDP and Meyer libraries were processed as the Sanger libraries above, except the trimadap tool (https://github.com/lh3/trimadap) was used for adapter trimming, the –T 0 parameter to bwa mem was not applied, and the bwa-postalt.js post-processing script was run after mapping.

Sample quality control, some of which is described in more detail below, included assessments of overall sequencing coverage, read error rates, genotype concordance to published genotype array data and cell-line chromosomal artefacts. For a few samples in the panel we had more than one library from different sources (i.e. Sanger, SGDP or Meyer, and PCR or PCR-free), and in each such case choose to include the library with the highest array genotype concordance, but excepting that rule in a few cases to avoid any of the 54 populations in the panel having an atypical composition of libraries from these different sources. After quality control and sample exclusions, the final set of 929 libraries contained 649 Sanger PCR-free, 152 Sanger PCR, 111 SGDP PCR-free, 9 SGDP PCR and 8 Meyer PCR libraries.

Previously published array genotypes available on the panel were lifted over from hg18 (*11*) and GRCh37 (*13*) to the GRCh38 assembly using the NCBI Remap tool and through looking up rs numbers in dbSNP (*42*).

*Capping of mapping qualities*

We noticed that the genotypes called from many of the Sanger PCR-free libraries displayed higher rates of discordance with array genotypes from the same individuals than the other sample sets, including the Sanger PCR libraries. To test if this could be due to cross-sample contamination caused by index hopping in the multiplexed sequencing runs performed for the PCR-free libraries, we ran the VerifyBamID tool (*43*) to estimate a per-library contamination rate (the "FREEMIX" estimate). These estimates were higher for many of the Sanger PCR libraries (in the highest cases 1-2%), and correlated strongly with the array discordance rate, strongly suggesting this was the cause of the reduced genotype accuracy (Figure S1A).

To reduce the impact of the index hopping on genotype accuracy, we applied a per-sample cap on the mapping qualities (MAPQ) of reads as a function of the contamination estimate (https://github.com/mcshane/capmq):

$$\max(MAPQ) = \max\left(20, 10 \cdot \log_{10}\frac{1}{FREEMIX}\right)$$

The rationale behind this is that if e.g. 0.1% of the reads are contaminant and do not actually derive from the given individual, for any given read we cannot, regardless of how well it has aligned to the reference genome, have more than 99.9% confidence that it reflects the genome sequence of the individual. We thus express that uncertainty by lowering the mapping quality of any read above this confidence level, in this example corresponding to a mapping quality of 30. However, we do not set the cap for any sample at lower than 20.

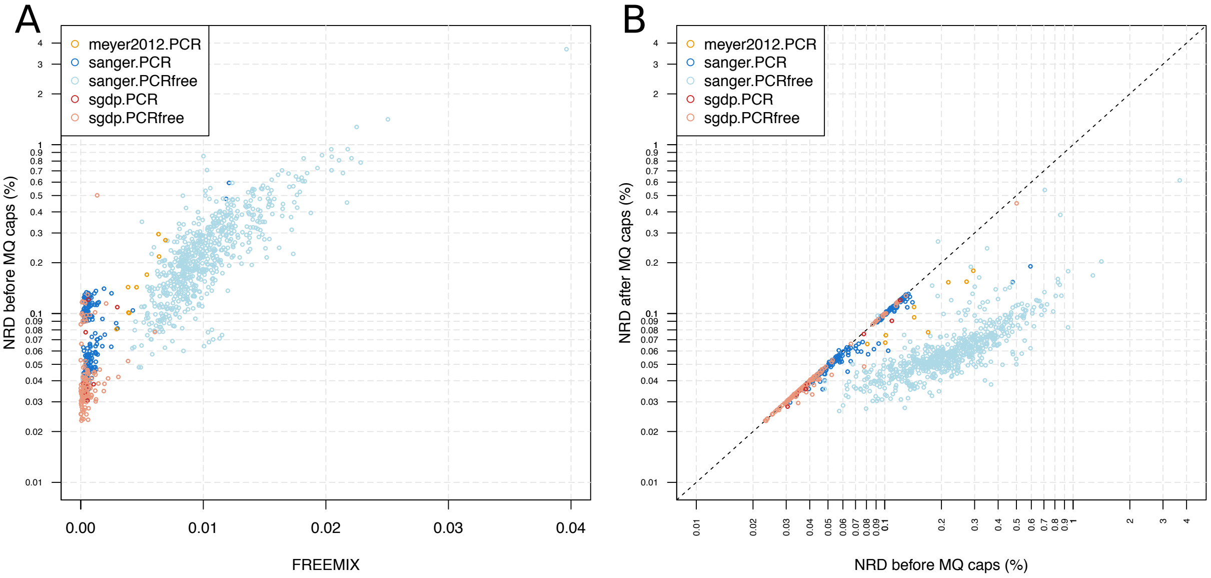


***Figure S1. Sample specific mapping quality caps offsets the effects of index hopping on genotype accuracy**. **A)** The FREEMIX contamination estimate strongly correlates with array genotype discordance (non-reference discordance, NRD) in the multiplexed Sanger PCR-free libraries. **B)** The application of sample specific mapping quality caps as a function of the FREEMIX estimate decreases the genotype discordance.*

Applying these sample-specific mapping quality caps substantially improved the array discordance rate for the Sanger PCR-free libraries, bringing them into the same range as non-multiplexed libraries (Figure S1B). It also slightly improved the rate for some other libraries. One library displayed a very high contamination rate (~4%) and a high array discordance even after capping mapping qualities, and was therefore marked as failing quality control.

*Cell line copy number alterations*

To identify any large-scale chromosomal copy number changes having occurred during culturing of the HGDP-CEPH lymphoblastoid cell lines from which DNA was obtained, we studied the mapped read coverage along the chromosomes of each sample. We calculated the coverage at approximately 300,000 single positions across the genome, and then plotted rolling means of these normalized by the genome-wide median. We visually inspected the plots for each sample and identified local deviations from the expected normalized copy number. As the cell line is a population of cells, the magnitude of the coverage deviation will be proportional to the fraction of the cells carrying the copy number alteration and can thus fall anywhere within the continuous range between the expected (1) and the most extreme possible values (1.5 in the case of a gain and 0.5 in the case of loss).

We marked 66 libraries as displaying at least one instance of mild deviation in coverage along a substantial stretch of a chromosome or major deviation in a megabase-sized stretch. Most events affected entire chromosomes or chromosome arms, but some smaller events were also observed. An especially large number of whole-chromosome gains were observed on chromosomes 9 and 12. We considered 9 of these 66 libraries to have deviations that were too extreme, informed by a computational experiment described below, and marked these as failing quality control. This included one sample (HGDP01097, Tujia) which did not display any copy number alterations, but which we discovered was completely homozygous along the entire length of chromosome 1 (we confirmed this was also the case in previously published genotype array data), most likely reflecting a cell line uniparental disomy event (though we cannot rule out that this reflects an actual germline uniparental disomy event in the sample donor). While several of the identified cases of unexpected copy number involved the sex chromosomes, particularly loss of chromosome Y in males and loss of chromosome X in females (Figure S2B), we did not exclude any sample on the basis of sex chromosome coverage alone. One self-reported male individual displayed what is likely an actual XXY karyotype rather than a cell line alteration.

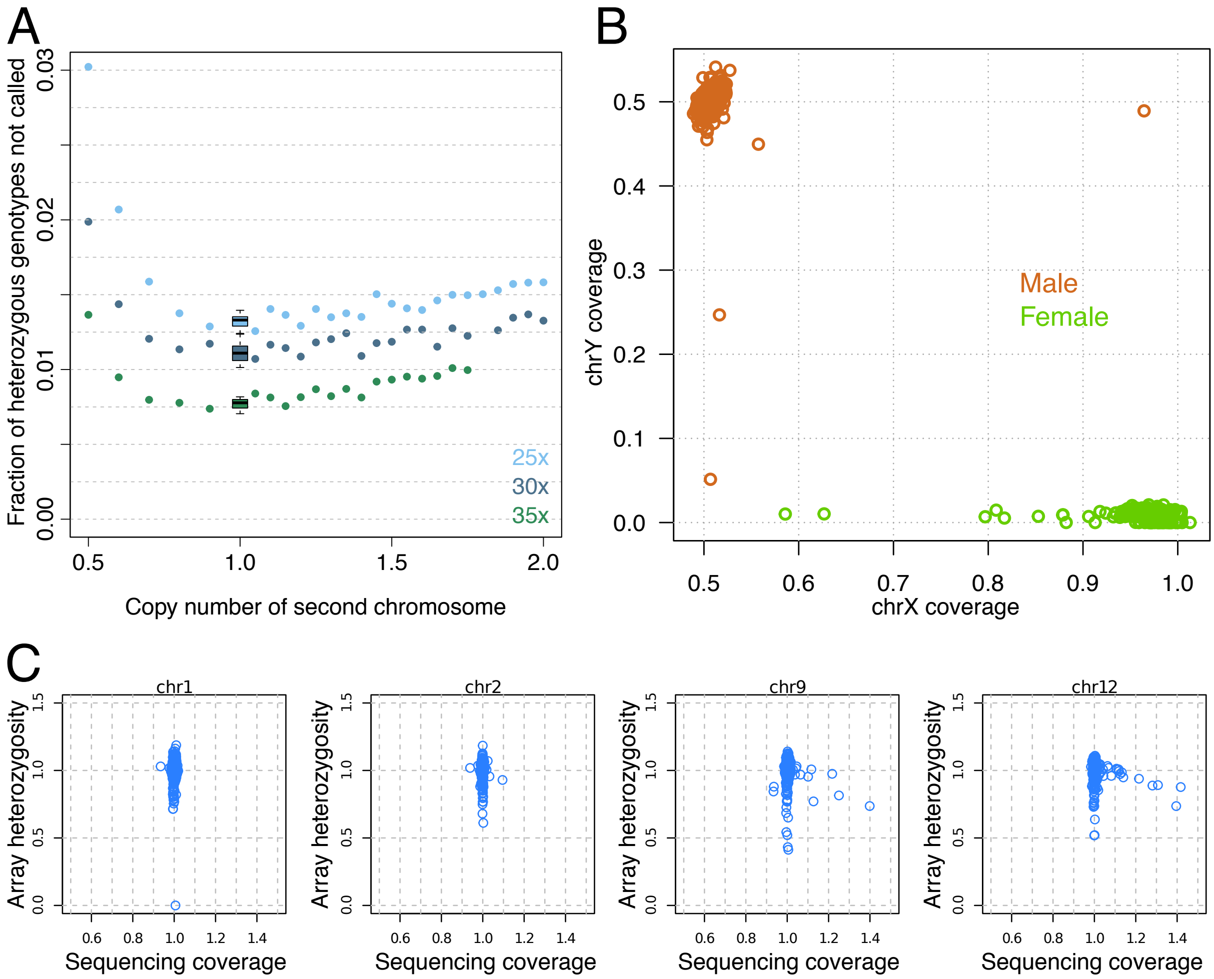


***Figure S2: The effects of cell-line chromosomal copy number alterations**. **A)** The fraction of heterozygous genotypes not correctly called in pseudo-diploid datasets constructed from two male X chromosomes down-sampled to varying degrees corresponding to copy number alterations of varying levels of severity, relative to a case with perfectly balanced copy number. Ten replicate experiments were performed for the balanced copy number case, the results of which are represented by box plots. The experiment was performed at three different levels of overall coverage; 25x, 30x and 35x. **B)** Sequencing coverage of chromosomes X and Y for all sequenced samples, relative to the genome-wide coverage, coloured by the self-identified gender of the sample donor. **C)** Sequencing coverage in our generated sequencing data of a chromosome in a sample against the fraction of heterozygous genotypes on that chromosome in that sample (normalized by chromosome and population) in a previously published array genotype dataset (11), displayed for two typical chromosomes as well as chromosomes 9 and 12 which display the largest number of whole-chromosome copy number alterations.*

We performed a computational experiment to assess the effects of coverage deviations on the accuracy of genotype calling at small nucleotide variants and determine whether it was appropriate to retain samples with mild deviations in our final dataset. We took the reads from the haploid chromosome X of two male individuals (HGDP00262 and HGDP00251, both Pathan) and subsampled these in varying proportions to generate synthetic diploid datasets corresponding to varying levels of coverage imbalance between the two chromosomes. We then called genotypes and determined how often the genotype called from a given imbalanced dataset matched that called from the perfectly balanced dataset at heterozygote SNPs (copy number changes will not impact the calling of homozygous genotypes). We found that copy number gains of a chromosome, even extreme ones, have only very minor effects on genotyping accuracy on high-coverage data (Figure S2A), consistent with the variant calling algorithm being able to tolerate some fluctuation from the expected balanced proportions of allele observations. However, we found that copy number losses result in larger effects on genotype accuracy. We therefore considered copy number losses as particularly strong reasons for sample exclusion.

DNA from these cell lines has previously been used to generate a large amount of data using a variety of technologies, and such data might potentially have been affected by the chromosomal alterations we identify here. To test to which extent a previously published and widely used array genotype dataset from the panel (*11*) was affected, we asked if the total sequencing coverage of a chromosome in our whole-genome sequencing data correlated with heterozygosity of the array genotypes. We found that increased coverage of a chromosome is associated with reduced heterozygosity in the array genotypes (Figure S2C), consistent with imbalanced allele counts leading to undercalling of heterozygotes. However, the effect is not very dramatic and thus is unlikely to have had much impact on population genetic analyses.

*Genotype calling and filtering*

We identified and genotyped SNPs and small indels using GATK HaplotypeCaller version 3.5.0, applying genotype priors without bias towards the reference allele through the "--input_prior 0.001 --input_prior 0.4995" arguments (*3*), the "--pcr_indel_model NONE" argument for the PCR-free libraries and the "--includeNonVariantSites" argument to include monomorphic sites in the output VCF files.

We wished to apply filters that are equally stringent for variant sites as for non-variant sites. Any filter that applies to variant sites but not to non-variant sites, e.g. GATK's Variant Quality Score Recalibration, comes with a risk of introducing a bias against variants and thereby introduce skews into various population genetic analyses that rely on the balance between these two classes of sites. We applied filters to the genotype calls using the GQ ("Genotype Quality") and RGQ ("Reference Genotype Quality") annotations that are produced by GATK for variant sites and non-variant sites, respectively. The GATK software outputs these as separate annotations and does not guarantee that they are comparable, but we performed a computational experiment to test how these annotations behave in practice. We performed single-sample calling for a given sample, and from these calls extracted the genotype annotations at the ~60 million sites which during the joint calling of the whole panel had been called as polymorphic SNPs but with a homozygous reference ("0/0") genotype for the given sample (meaning some other sample in the panel carried an alternative allele at this site). This thus gives us a set of sites where the genotypes for the given sample has been evaluated by GATK once as non-variant sites and assigned RGQ values, and once as variant sites and assigned GQ values. For the sample HGDP01377, out of the 61,028,821 surveyed sites, GQ != RGQ at 6,650,620 sites (10.9%), but |RGQ – GQ| > 1 only at 23,372 sites (0.038%), suggesting most differences are just due to numerically rounding off the values into different consecutive integer bins. These comparisons are slightly complicated by multi-allelic sites in the joint calls – restricting to bi-allelic SNPs, |RGQ – GQ| > 1 at 0 sites. We performed the same experiment for two additional samples (HGDP00995 and HGDP01377) and obtained similar results. Thus, at least on this dataset called with non-reference biased priors, GQ and RGQ behave extremely similarly in practice and we thus take the same threshold applied to these annotations as providing equally stringent filtering for variant and non-variant sites.

We proceeded to filter the genotypes as follows. For each sample, we set any genotype to missing if it had a GQ or RGQ value equal to or lower than 20, or a coverage ("DP" annotation) equal to or greater than 1.65 times the genome-wide average coverage for the sample.

We also computed two site level annotations: GATK's Variant Quality Score Recalibration (VQSR) and excess heterozygosity. VQSR was run on the unfiltered genotypes, for SNPs with the hapmap_3.3.hg38.vcf.gz, 1000G_omni2.5.hg38.vcf.gz and 1000G_phase1.snps.high_confidence.hg38.vcf.gz variant sets from the GATK GRCh38 bundle for training and the former two for truth sets, and using the QD, MQRankSum, ReadPosRankSum, FS and MQ annotations as features. VQSR was run for indels with the Mills_and_1000G_gold_standard.indels.hg38.vcf.gz variant set for training and truth sets, and using the FS, ReadPosRankSum, InbreedingCoeff, MQRankSum and QD annotations as features. After annotating the VCF with the obtained VQSLOD values, another annotation "VQSRMODE" was added to each site to indicate whether it was evaluated in the SNP or indel mode, as at multi-allelic site this information might otherwise not always be retained when filtering out one of the alleles or subsetting to a sample set in which the evaluated allele is not present. The excess heterozygosity "ExcHet" annotation was calculated on a per-allele basis using the bcftools (*44*) fill-tags plugin. We then marked any site with a SNP VQSR score below -8.3929 or indel VQSR score below -1.0158 with the "LOW_VQSLOD" filter tag, and any site harbouring an allele with an ExcHet value equal to or larger than 60 (corresponding to an excess heterozygosity p-value of $10^{-6}$) with the "ExcHet" filter tag. For downstream analyses, we did not make use of the VQSR scores for filtering, unless noted, but we excluded sites tagged with excess heterozygosity from all analyses.

We wished to restrict certain analyses to bi-allelic SNPs. Rather than excluding the entire site in case a third allele and/or an indel allele is present, which would mean that an e.g. an additional allele observed in a single individual would lead to the exclusion of a site harbouring two otherwise informative and perfectly usable common alleles, for most analyses we instead masked out alleles. If an allele is lower in frequency than two other alleles at the same site, or if the allele is an indel allele, we excluded the allele and set the genotype of any individual carrying a copy of the allele to missing, allowing us to retain the site and the two most common SNP alleles.

We annotated SNP variants with two ancestral allele tags: one using the allele predicted by the Ensembl 8 primates EPO alignments (cc21_ensembl_compara_86), and one using the Chimpanzee allele in the GRCh38-PanTro4 alignment from the UCSC genome browser.

We noticed that in population genetic analyses (e.g. principal component analyses) of the unfiltered genotype calls, there were noticeable batch effects between the libraries of different sources, primarily between Sanger and SGDP libraries but also to a smaller extent between PCR and PCR-free libraries. The genotype filtering described above reduces these effects, as does increasingly stringent VSQLOD thresholds, and when applying the accessibility mask described above the effects are not discernible. However, we still urge any users of the data to be aware of the possibility that some sensitive analyses could still be affected by these effects, particularly if not applying the accessibility mask.

*Overview of small variant callset*

Our filtered variant call set across 929 samples, excluding sites labelled with excess heterozygosity, contained 75,310,370 variant sites. This included 67,325,692 SNPs and 8,797,538 indels. 3,085,457 of all variants were multi-allelic, and 855,977 of SNP sites (1.3%) were multi-allelic. The transition/transversion ratio was 1.88. 29,279,819 SNPs (43.5%) and 3,431,934 indels (39.0%) were singletons (the alternative allele observed only in one copy across the individuals).

We compared the identified variants with those identified by the 1000 Genomes Project (*2*) (the 20170504 GRCh38 lift-over version). While both datasets contain millions of variant alleles that are not found in the other dataset, the vast majority of these are very low frequency, including many singletons, and a simple count of overlapping versus unique variants does not reveal to which extent either of the datasets contains variants that might be of higher frequency in particular populations. We therefore calculated, for each variant present in one dataset but not the other, its maximum allele frequency in any population. To avoid cases where a variant is actually present in the sequenced individuals in both datasets but absent from one of the VCFs because of technical issues in variant calling and/or lift-over, we excluded 1000 Genomes variants that did not have the "GRCH37_38_REF_STRING_MATCH" tag (indicating that the reference allele string matches between GRCh37 and GRCh38), and we excluded from both datasets any variant that had a global allele frequency across the dataset of 30% or higher (the reasoning being that it would be very unlikely that such a globally common variant would not be sampled by both of these datasets).

The HGDP dataset contains a larger number of populations and these populations have smaller sample sizes than in the 1000 Genomes dataset, leading to higher variance in the per-population allele frequency estimates and thus potentially resulting in an upwards bias when finding the maximum allele frequency across populations. To enable a fair comparison, we down sampled the 1000 Genomes dataset to resemble the HGDP dataset: for each of the 26 1000 Genomes populations, two random subsets were constructed with sizes determined by sampling without replacement from the set of HGDP population sizes. The distribution of maximum allele frequencies across these size-matched 1000 Genomes populations for variants not present in HGDP shifted upwards, but variant numbers still remained much lower than the number of variants private to HGDP. We performed this down-sampling three times, and the results were very similar across the three replicates.

*gVCF construction*

Having genotype calls at monomorphic sites is very valuable, as they are needed in certain population genetic analyses and also as they make appropriate merging of different datasets possible. We produced VCFs containing every called site, but these files are very large and therefore challenging to distribute and store. We therefore constructed per-sample gVCFs ("genomic" VCFs) in which consecutive sites with homozygous reference genotypes and similar confidence level are grouped into single, block VCF records. We grouped such sites into four classes of blocks, in which the GQ and RGQ annotations are collapsed into a single GQ annotation:

| **Block definition** | **GT field** | **FILTER tag** |
|---|---|---|
| DP ≥ mean(DP) × 1.65 | set to missing | EXCESS_DP |
| GQ>60 & DP < mean(DP) × 1.65 | unchanged | . |
| GQ>20 & GQ<=60 & DP < mean(DP) × 1.65 | unchanged | . |
| GQ<=20 | set to missing | GQ20 |

For each block record, only the DP and GQ genotype annotations are retained and used to represent the minimum value across all sites contained within the block. If a site with a non-reference genotype fails any filter, we set the genotype to missing but still retain the alternative allele and all the genotype annotation fields, such that it's always possible to restore the genotype called by GATK if desired. For sites with non-reference genotypes we also carried over a few site-level annotations from the joint variants VCF (ExcHet, VQSLOD, VQSRMODE).

These gVCF files constitute compact representations of the genome sequences of single individuals, while retaining most of the information of relevance to assess the uncertainty of a given genotype call and allow for custom filtering. The mean size of these files across the 929 samples is 1.32 GB (standard deviation = 0.48, min = 0.25, max = 3.25), with the variation largely explained by differences in sample coverage ($r_{coverage,file\ size} = -0.79$).

*Accessibility mask*

Rather than variant level filtering, for most analyses we instead relied on an accessibility mask. This mask was constructed on the basis of the 1000 Genomes Project's strict mask for GRCh38 (20160622 version), which is based on coverage and mapping quality patterns in the 1000 Genomes Project dataset. From this mask, we also subtracted any regions of the primary GRCh38 assembly which have alternative loci or patch scaffolds, as defined by the NCBI assembly resource for the GCA_000001405.15_GRCh38 entry. As we performed variant calling on GRCh38 read alignments that had not been post-processed to adjust the mapping qualities in an alt-aware manner, genotypes in these regions are likely not reliable. We also subtracted all sites tagged with excess heterozygosity, considering them unsuitable for genotyping from Illumina reads. The resulting masks leaves approximately 73% of the primary assembly for analyses, and unless otherwise noted our analyses are restricted to this mask. While we find that restricting to this mask without further site-level filtering appears suitable for the population genetics analyses we perform here, other types of analyses using this dataset might benefit from alternative filtering strategies, e.g. using the VQSR scores or other annotations. In particular, analyses of variants of potential functional, medical or selection relevance might benefit from interrogating variants also in the approximately 27% of the genome that falls outside of this mask.

*10x Genomics sequencing and haplotype phasing*

We selected 26 samples from 13 populations to process with the 10x Genomics Chromium technology (*8*), producing linked reads with long-range physical information that enable haplotype phasing. We selected 13 of the 54 populations representing key ancestries. Within each of these populations, we tried to fulfil several criteria when selecting which two samples to process: an absence of any large-scale chromosomal copy number alterations; a typical ancestry profile with respect to their population as assessed through principal component and model-based clustering analyses; no evidence of high relatedness between the two individuals; and male individuals if possible, to obtain Y chromosome data.

DNA quality and molecule length distributions were assessed using the Agilent TapeStation, and Chromium libraries were then constructed for the 26 selected samples and sequenced on single lanes of Illumina HiSeqX machines (2×151 bp reads, average coverage 30.2X) (Table S1). Four of these libraries were described and used for population genetic analyses in a previous publication (*45*). The resulting barcoded reads were processed using the 10x Genomics Long Ranger software version 2.1.2 with genotype calling through GATK HaplotypeCaller 3.5.0, to obtain phased VCFs for each individual. In order to maintain only a single set of genotype calls for these 26 individuals for which we had calls both from the standard Illumina data and from the Chromium data, we lifted over the haplotype phase information at heterozygous sites from the latter to the former. Any genotype where the two calls disagreed were set to unphased, and variants within phase blocks that contained only one variant were also set to unphased. The resulting VCF files thus contained the unaltered genotypes that we called from the standard Illumina data, with only haplotype phase information obtained from the Chromium experiments.

*Table S1: Details on the results of the 10x Genomics Chromium sequencing experiments. Metrics were calculated by the 10x Genomics Loupe software.*

| Sample | Population | Average Molecule Length | Linked Reads per Molecule | Sequencin g coverage | DNA fragments >20kb (%) | DNA fragments > 100kb (%) | Phase block N50 | SNPs phased (%) |
|---|---|---|---|---|---|---|---|---|
| HGDP00460 | Biaka | 16,299 | 12 | 33.0 | 37.8 | 2.07 | 1,524,737 | 98.9 |
| HGDP00472 | Biaka | 8,258 | 6 | 30.2 | 17.5 | 2.79 | 191,984 | 97.5 |
| HGDP00562 | Druze | 8,294 | 6 | 24.0 | 12.8 | 2.12 | 119,503 | 97.0 |
| HGDP00580 | Druze | 15,756 | 20 | 31.1 | 33.3 | 3.47 | 417,052 | 98.7 |
| HGDP00774 | Han | 12,518 | 7 | 31.2 | 25.8 | 2.02 | 222,937 | 97.7 |
| HGDP00819 | Han | 22,113 | 15 | 35.2 | 55.8 | 2.00 | 601,565 | 98.7 |
| HGDP01013 | Karitiana | 30,225 | 22 | 32.9 | 69.6 | 3.14 | 596,564 | 98.3 |
| HGDP01019 | Karitiana | 7,713 | 4 | 23.2 | 11.3 | 1.91 | 81,409 | 95.7 |
| HGDP00450 | Mbuti | 14,468 | 10 | 35.0 | 33.5 | 2.03 | 990,820 | 98.7 |
| HGDP01081 | Mbuti | 12,242 | 9 | 32.7 | 24.4 | 2.70 | 560,811 | 98.6 |
| HGDP00549 | PapuanHighlands | 8,724 | 5 | 25.8 | 14.3 | 1.78 | 126,844 | 96.8 |
| HGDP00551 | PapuanHighlands | 14,200 | 15 | 31.2 | 26.0 | 3.21 | 206,857 | 97.7 |
| HGDP00542 | PapuanSepik | 11,604 | 6 | 29.1 | 24.4 | 1.81 | 176,726 | 97.4 |
| HGDP00547 | PapuanSepik | 11,157 | 11 | 29.0 | 16.9 | 3.48 | 139,899 | 97.0 |
| HGDP00224 | Pathan | 7,214 | 5 | 27.6 | 10.7 | 2.41 | 81,721 | 96.9 |
| HGDP00228 | Pathan | 7,589 | 5 | 24.6 | 10.9 | 1.89 | 101,084 | 97.0 |
| HGDP01043 | Pima | 27,111 | 28 | 33.2 | 64.7 | 2.72 | 591,200 | 98.6 |
| HGDP01056 | Pima | 31,234 | 13 | 26.7 | 70.8 | 3.28 | 636,630 | 98.9 |
| HGDP01029 | San | 11,063 | 11 | 29.7 | 17.7 | 2.52 | 406,739 | 98.5 |
| HGDP01032 | San | 22,654 | 20 | 33.7 | 56.2 | 2.78 | 2,941,155 | 98.9 |
| HGDP00670 | Sardinian | 12,126 | 14 | 32.9 | 24.4 | 2.64 | 292,920 | 98.4 |
| HGDP01067 | Sardinian | 33,980 | 37 | 35.9 | 74.2 | 4.73 | 1,463,166 | 98.8 |
| HGDP00946 | Yakut | 7,376 | 6 | 30.4 | 10.2 | 2.34 | 80,637 | 96.3 |
| HGDP00954 | Yakut | 5,820 | 4 | 23.6 | 8.94 | 2.21 | 40,003 | 94.9 |
| HGDP00930 | Yoruba | 7,913 | 6 | 30.9 | 12.0 | 2.54 | 129,564 | 97.7 |
| HGDP00931 | Yoruba | 9,279 | 7 | 32.5 | 15.3 | 2.86 | 256,356 | 98.3 |

#### *Structural variation calling*

We called copy number variants using GenomeSTRiP v2.00 (*46*) using default parameters. We initially ran the algorithm jointly on all 965 available libraries, including libraries not passing quality control for short variant calling, and including the Meyer libraries. However, we found the quality of resulting calls for the Meyer libraries to be low and re-ran the algorithm excluding them.

As we have a number of duplicate samples prepared using PCR and PCR-free libraries for quality control purposes, we ran the algorithm twice for these, separately for each library preparation set, in each case together with the rest of the dataset. We found that more variants were called for PCR-based libraries compared to PCR-free libraries. We also found that samples prepared with PCR libraries had a larger number of shared heterozygous calls that are missing from the PCR-free libraries, suggesting these are artefactual calls. We subsequently excluded variants with excessive heterozygosity as computed by bcftools v1.9 (ExcHet < 0.0001) separately for each library preparation and sequencing location set (i.e. SGDP PCR, SGDP PCR-free, Sanger PCR and Sanger PCR-free). For the SGDP PCR samples we used ExcHet < 0.05 as this set had only 9 samples.

We investigated potential cell-line artefacts in further details by analysing coverage across the genome for each sample, as CNV calling might be more sensitive to such artefacts than short variant calling. From the 929 samples included in the SNP analysis, we excluded the Meyer samples as well as 10 samples that displayed evidence of alterations across multiple chromosomes. We also masked regions in 74 samples that showed more limited putative alterations, meaning we retain the samples but we did not consider variants called for them within these masked regions. This resulted in a VCF file with 911 samples in which GenomeSTRiP called 50,474 CNVs.

We examined the calls and found cases where the algorithm splits variants into multiple shorter entries which are not always overlapping. This a known behaviour of the GenomeSTRiP CNV pipeline, and it seems to occur when there are variants with different copy numbers across different individuals within a sub-segment of a larger variant. This issue can also occur if a low-quality variant is found within a larger CNV. To address these issues and be able to more accurately estimate the total number of identified CNVs in our dataset, we merged high quality (CNQ > 12) calls that have same diploid copy number and are within 50 kb of each other, for each sample individually. For the X-chromosome we performed this separately for male and female samples. At this step we excluded three further samples that showed elevated numbers of calls compared to the rest of the samples (thus leaving 908 samples for downstream analyses). All variants were then merged using bedmap v2.4.35 (*47*) based on 100% overlap. This resulted in 38,903 autosomal variants and 1,094 variants on the X-chromosome.

*Statistical haplotype phasing*

As we only had 10x Genomics experimental phasing data for a subset of samples, we also performed statistical phasing of the whole panel. We restricted statistical phasing to biallelic SNPs (by masking out third or higher alleles and indel alleles, rather than excluding entire sites) lacking filter tags. To take advantage of a large external reference panel without having to exclude variants not present in the reference panel, we applied a scaffold-based method implemented as a genotype calling mode in SHAPEIT2 (*48*). We first obtained the scaffold by phasing the genomes using Eagle (v2.3.2) (*49*) after three PBWT iterations with 4,956 genomes from the African Genome Resources (https://www.apcdr.org/) in the reference panel (this includes the full 1000 Genomes Project panel). Subsequently we divided the unphased chromosomes into windows each spanning 2,400 SNPs with 200 overlapping ones between adjacent windows. Beagle 4.1 (*50*) was run with the -gtgl option to produce genotype probabilities in each window. Based on the genotype probabilities, the variants were then mapped onto the phased scaffold with the --call and --input-scaffold options in SHAPEIT2. To avoid under- and overflow errors (as described in (*3*)), we ran SHAPEIT2 in the same windows as in the previous step instead of assigning windows by physical lengths, and further enlarged the windows when such errors still occurred on rare occasions. Missing genotypes in the unphased input dataset were reset to missing in the phased dataset for consistency.

We evaluated the accuracy of the statistical phasing by comparing the inferred haplotypes to those obtained for the samples sequenced with 10x Genomics linked reads. If adjacent phase-resolved heterozygous sites in the same phasing block in the 10x genome formed a different haplotype structure than that in the statistically phased genome, we considered it a switch error. Table S2 lists the switch error rates on chromosome 1, measured as the total number of switch errors divided by the total number of possible switches (namely the number of heterozygous sites minus one). When singleton alleles are excluded, the highest switch error rates of around 1.2% are found in the San and Papuan populations, which roughly corresponds to on average one switch error per 160 kB. We therefore do not expect phasing errors to have a detectable impact on downstream analysis targeting haplotypes substantially shorter than this.

*__Table S2: Evaluation of statistical phasing accuracy.__ Switch error rates were measured for 26 individuals against experimentally phased haplotypes obtained using 10x Genomics linked reads for the same individuals.*

| Individual | Population | Region | Switch error rate with singletons | Switch error rate without singletons |
|---|---|---|---|---|
| HGDP00460 | Biaka | Africa | 0.0140 | 0.0044 |
| HGDP00472 | Biaka | Africa | 0.0157 | 0.0064 |
| HGDP00450 | Mbuti | Africa | 0.0144 | 0.0051 |
| HGDP01081 | Mbuti | Africa | 0.0155 | 0.0052 |
| HGDP01029 | San | Africa | 0.0378 | 0.0131 |
| HGDP01032 | San | Africa | 0.0374 | 0.0122 |
| HGDP00930 | Yoruba | Africa | 0.0092 | 0.0033 |
| HGDP00931 | Yoruba | Africa | 0.0097 | 0.0027 |
| HGDP01013 | Karitiana | America | 0.0086 | 0.0047 |
| HGDP01019 | Karitiana | America | 0.0066 | 0.0052 |
| HGDP01043 | Pima | America | 0.0076 | 0.0039 |
| HGDP01056 | Pima | America | 0.0081 | 0.0052 |
| HGDP00224 | Pathan | Central & South Asia | 0.0130 | 0.0043 |
| HGDP00228 | Pathan | Central & South Asia | 0.0124 | 0.0040 |
| HGDP00774 | Han | East Asia | 0.0156 | 0.0049 |
| HGDP00819 | Han | East Asia | 0.0167 | 0.0054 |
| HGDP00946 | Yakut | East Asia | 0.0061 | 0.0033 |
| HGDP00954 | Yakut | East Asia | 0.0088 | 0.0046 |
| HGDP00670 | Sardinian | Europe | 0.0107 | 0.0033 |
| HGDP01067 | Sardinian | Europe | 0.0115 | 0.0038 |
| HGDP00562 | Druze | Middle East | 0.0067 | 0.0031 |
| HGDP00580 | Druze | Middle East | 0.0125 | 0.0038 |
| HGDP00549 | PapuanHighlands | Oceania | 0.0237 | 0.0117 |
| HGDP00551 | PapuanHighlands | Oceania | 0.0281 | 0.0120 |
| HGDP00542 | PapuanSepik | Oceania | 0.0281 | 0.0119 |
| HGDP00547 | PapuanSepik | Oceania | 0.0278 | 0.0115 |

*Metadata curation*

There is some inconsistency in the population labels used in the prior literature on the HGDP-CEPH samples. We reviewed the population labels, aiming to adhere as much as possible to the labels provided in the official CEPH sample documentation but also to define the most scientifically useful groupings with labels that are ethnically, linguistically and geographically appropriate. After this review, we arrived at 54 population labels. While most differences to previously used labels involve only minor spelling variations, a few cases involve more notable changes. We comment on some of the population labels and the motivation behind our choices, as well as any notes on the coordinates used to indicate geographical origins, here:

- **BantuSouthAfrica** and **BantuKenya**: These have sometimes been collapsed into a single "Bantu" label, however we use two labels as they are substantially separated both geographically and genetically ($F_{ST} \approx 0.008$). The South African samples have sometimes been further subdivided into "South-Western" and "South-Eastern" sets or into individual language groups, however to retain a decent sample size for the population we do not make use of these further subdivisions.

- **Colombian**: These have sometimes been subdivided into two individual language groups (Piapoco and Curripaco), however to retain a decent sample size we do not subdivide these samples (the CEPH documentation also does not describe which of the samples are from which of the two language groups).

- **Han** and **NorthernHan**: While representing individuals from the same ethno-linguistic group, given the large number of Han individuals in the panel we made use of the sometimes utilized second label for the set sampled in northern China, for which we assign new geographical coordinates based on documentation from the original sample collection.

- **Mongolian**: "Mongola" has been used, however we believe this reflects a spelling error. The geographical coordinates reported for this population in (*51*) differs slightly from those in the CEPH documentation – we believe the latter are the correct coordinates.

- **Bougainville**: "Melanesian" or "NAN Melanesian" (NAN=Non-Austronesian language) has been used for this group from Bougainville Island, but is overly generic. The name of a specific language group has sometimes been used, but we do not make use of this as it is not part of the CEPH documentation.

- **PapuanSepik** and **PapuanHighlands**: The prior literature has used the single label "Papuan" for these samples from Papua New Guinea, however it has been shown that they consist of two genetically highly distinct ($F_{ST} \approx 0.03$) subsets, one with affinities to populations in the eastern highlands and one with affinities to populations in the Sepik river region of the northern lowlands of New Guinea (*45*), and we thus separate them into two separate labels. For the PapuanSepik population, we use the geographical coordinates provided in the CEPH documentation, which are consistent with a Sepik region location. For the PapuanHighlands population, we assign new coordinates on the basis of the results reported in (*45*).

- **San**: "Ju|'hoan North" has been used, but this label is not used in the CEPH documentation and so we use the "San" label, even if it is more generic.

- **BergamoItalian**: "North Italian" has been used, but we include the Bergamo origin of these samples in the label to clarify its distinction from the Tuscan population that is also part of the panel.

### Population genetic analyses

#### *f-statistics analyses*

We calculated $f_4$ and D-statistics using the ADMIXTOOLS package version 5.0 (*13*). As the ADMIXTOOLS programs used excessive memory when attempting to use all variants to calculate large numbers of statistics, e.g. the 948,753 $f_4$-statistics corresponding to all possible relationships among the 54 populations, we ran ADMIXTOOLS separately on 5 Mb blocks across the genome and then performed our own block jackknifing across the per-block estimates. We verified on a small subset of statistics that the obtained $f_4$ values and Z-scores were highly correlated to those calculated one by one directly by ADMIXTOOLS.

#### *Effects of variant ascertainment on population genetic analyses*

The ideal class of variants for analyses that rely on genetic drift are those that were polymorphic in the shared ancestral population of the given populations under study. One approach to approximate this ancestral polymorphism is to ascertain variants in an outgroup. We ascertained SNPs that are polymorphic among three high-coverage archaic human genomes: the Altai Neanderthal, the Vindija Neanderthal and the Denisovan genome. Within the accessibility mask, this resulted in 2,809,464 variants. Out of these, 1,350,097 were also polymorphic within the set of 929 modern humans. Out of these, 931,790 (69.0%) were polymorphic also among Africans only (101 individuals, excluding three BantuKenya individuals displaying some non-African ancestry components in ADMIXTURE runs), demonstrating that the presence of these variants in present-day modern human populations is to a large extent explained by them being present in the shared ancestral population of modern and archaic humans, and only to a smaller extent a consequence of archaic admixture into non-Africans.

We compared results of various population genetic analyses obtained on all the discovered variants and the above outgroup ascertained set to sets of variants present on commonly used genotyping arrays: the Illumina 650K (or "Li 2008") array (*11*), the Humans Origins array (*13*) and the Illumina Multi-Ethnic Global Array ("MEGA"). We lifted over the site lists of these arrays to GRCh38 using the NCBI Remap tool. While most of the samples in the HGDP-CEPH panel have been typed on the former two arrays, rather than using those datasets directly we extracted the genotype calls made from our whole-genome sequencing data on the sites present on the arrays, such that genotypes at any given site are held constant and only the sites used differ. We found:

- $f_4$-statistics calculated using array sites are highly correlated to those calculated using all variants or outgroup ascertained variants, overall. However, some statistics, especially those involving African populations, deviate in the array sites as a result of ascertainment bias (Figure 1C,Figure S3A). The overall magnitude of the $f_4$ statistics differs systematically between array sites and all sites, but this just reflects the overall larger number of variants included in the latter, many of which will not be polymorphic among the set of four populations used in a given statistic and therefore just contributes to smaller allele frequency differences on average.

- Estimates of individual ancestry components using ADMIXTURE (*52*) are cleaner when using outgroup ascertained sites, with less "leaking" of a given regional ancestry component into individuals outside that region (Figure S3B). For example, running ADMIXTURE on k=5 using the Li 2008 sites, individuals from Central and South Asian populations are assigned 5-10% of the Oceanian component, but this is substantially reduced with outgroup ascertained sites. Similarly, it reduces the fraction of Native American component in Europeans.

- $F_{ST}$ is generally slightly overestimated when using array sites, especially so when comparing African to non-African populations (Figure S3C).

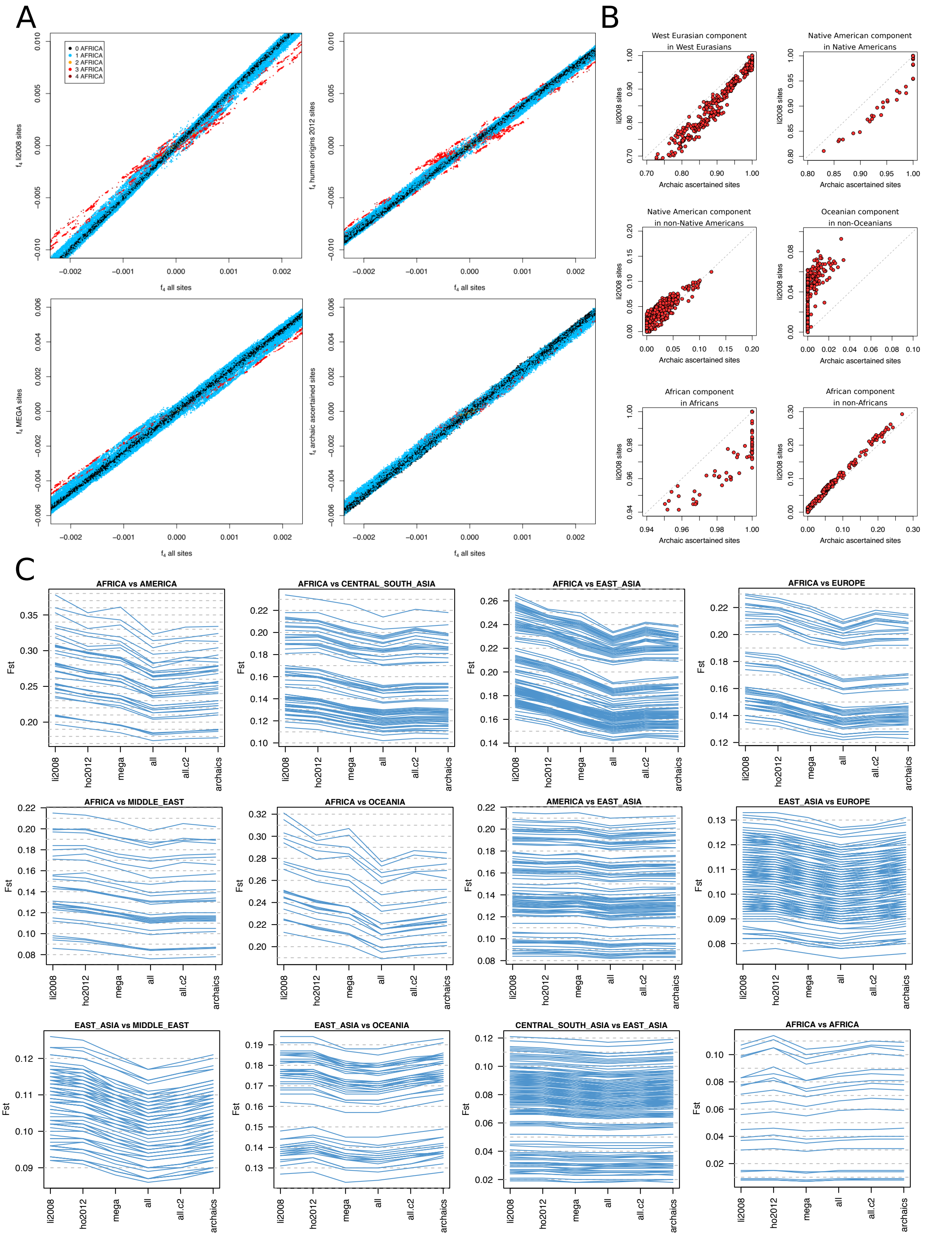


***Figure S3. The effects of variant ascertainment on population genetic analyses****. A) Comparisons of all possible f4 statistics involving the 54 populations calculated using different sets of array sites and sites ascertained in archaic genomes, against the values calculated using all discovered variants. Points are coloured according to the number of African populations included in the statistic. B) Estimates of individual ancestry components using ADMIXTURE at k=5, at which the five components correspond well to five major regional ancestries. Compared to using the Li 2008 array sites, the sites ascertained in archaic genomes leads to cleaner ancestry estimates with less "leaking" of a given regional ancestry component into individuals outside that region. C) Estimates of $F_{ST}$ for a selection of population pairs, grouped by region, using the Li 2008 array sites ("li2008"), the Human Origin 2012 sites ("ho2012"), the MEGA array sites ("mega"), all variants discovered in the sequencing data ("all"), all variants excluding singletons ("all.c2") and variants ascertained in archaic genomes ("archaics"). Each line connects the $F_{ST}$ values for one pair of populations across the different set of sites.*

*Region specific variants*

We studied the number and frequency distributions of variant alleles that are private to a particular region of the world, meaning alleles that have counts of zero across all individuals outside the given region. For these analyses, we did not restrict only to variants falling within the accessibility mask, with the rationale that technical errors are unlikely to result in genotype distributions that correlate perfectly with geographical labels. To reduce the effects of recent admixture between regions, we excluded individuals displaying evidence of such admixture in model-based clustering analyses. We ran ADMIXTURE (*52*), on 1,350,097 bi-allelic SNPs ascertained as polymorphic among three high-coverage archaic human genomes, with five ancestry components. The components obtained corresponded well to five continental-level ancestries: sub-Saharan African, West Eurasian, East Eurasian, Native American and Oceanian. We used the estimated per-individual ancestry proportions together with the population and region level metadata to define which individuals should be included when counting variants private to a given region (the "ingroup") and which individuals should be used to ascertain the allele count of zero (the "outgroup") (Table S3). Our rationale when defining these criteria was that, if we are looking for variants that are found in region A but not region B, it is more important to exclude individuals from B with recent admixture from A than vice versa – the inclusion of the former might cause otherwise private A variants to be observed in B individuals and therefore not be identified as private, while the inclusion of the latter will only lead to underestimation of the allele frequencies of private variants in A.

***Table S3. Defining regions for analyses of region-specific variants****. These were used to count variants that were present in the "Ingroup" and absent from the "Outgroup". Labels in all capital letters denote continental-level region labels, while those in lower case letters denote populations. The notation "anc(X)" denotes estimates of individual fractions for an ancestry component X, as estimated using ADMIXTURE at k=5.*

| **Region label** | **Ingroup** | **Outgroup** |
|---|---|---|
| **AFRICA** | AFRICA | !AFRICA & anc(AFRICA) <= 0.01 |
| **CENTRAL AFRICA** | Biaka or Mbuti | (Biaka or Mbuti) |
| **San** | San | !(San or BantuSouthAfrica) |
| **OCEANIA** | OCEANIA | !OCEANIA |
| **AMERICA** | AMERICA & anc(AMERICA) => 0.95 | !AMERICA |
| **CENTRAL AMERICA** | (Pima or Maya) & anc(AMERICA) => 0.95 | !(Pima or Maya) |
| **SOUTH AMERICA** | (Colombian or Surui or Karitiana) & anc(AMERICA) => 0.95 | !(Colombian or Surui or Karitiana) |
| **EUROPE** | EUROPE | !EUROPE & !(AMERICA & anc(WestEurasian) >= 0.01) |
| **EAST ASIA** | EAST_ASIA | !EAST_ASIA & !Hazara & !Uygur & !(OCEANIA & anc(EastEurasian) >= 0.01) |
| **MIDDLE EAST** | MIDDLE_EAST | !MIDDLE_EAST |
| **CENTRAL & SOUTH ASIA** | CENTRAL_SOUTH_ASIA & anc(AFRICA) <= 0.05 | !CENTRAL_SOUTH_ASIA |
| **NON_AFRICA_X** | region X as defined elsewhere | AFRICA & anc(AFRICA) >= 0.91 |

When analysing CNVs, we set any individual CNV genotypes with GQ < 20 to missing and excluded variants with >15% missingness across individuals (calculated in the VCF after filtering but before merging of adjacent variants). To further conservatively avoid overcounting single CNVs that have been called as multiple adjacent entries, we subsequently merged variants with similar allele frequencies and the same copy number lying within 25 kb of each other and only report the variant with the lowest genotype missingness.

If merged variants have the same missingness but slightly different allele frequencies, we calculated and report the average frequency.

To assess whether the high number of high-frequency private Oceanian CNVs could be expected by sampling noise alone or representing an enrichment indicative of positive selection, we randomly sampled 123 private Oceanian SNPs (matching the number of private Oceanian CNVs, with minimum allele count > 2) 1000 times and compared the frequency distributions in these random samples to the observed CNV distribution. We observed 12 sample sets with a variant having an equal or higher frequency than the single most frequent CNV (at a frequency of 82.14%). However, among these 12 sets, 10 set had only one variant >50% frequency and two sets had two variants >50%, whereas in the observed CNV distribution we find four variants at >50% frequency. The observed CNV distribution is thus highly unlikely to be the result of sampling noise.

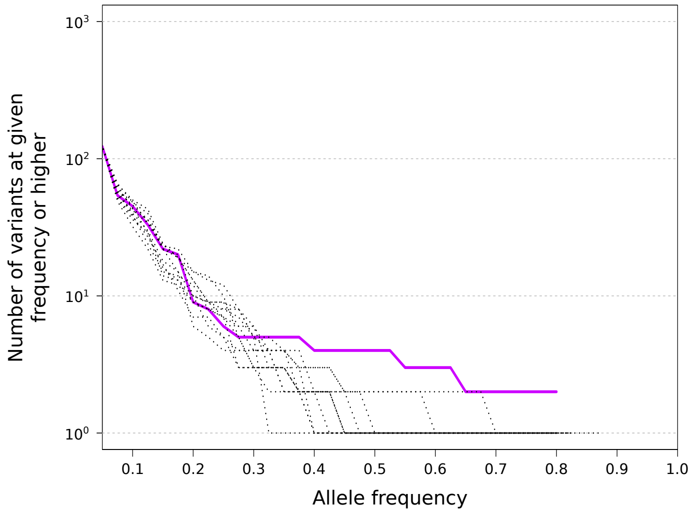


***Figure S4: Assessing the statistical significance of the high number of high-frequency private Oceanian CNVs.*** *The solid line displays the number of CNVs private to Oceanian populations that have an allele frequency in those populations equal to or higher than the corresponding value at the horizontal axis. The dashed lines are the results of 1000 random samples of equal numbers of private Oceanian SNPs, displaying only the most extreme 12 samples out of these that contained a variant reaching as high a frequency as the highest-frequency observed CNV.*

### Time depth of population structure

#### *MSMC2 split time analyses*

Pairwise population separation histories were studied using the MSMC2 software (*21, 27*). Input files were prepared from genotypes on all called sites, including non-variant sites, and with haplotype phase obtained from the 10x Genomics Chromium experiments we had performed on two individuals each from 13 populations. MSMC2 v2.0.2 was run on eight haplotypes (four from each of the two populations) with the "--skipAmbiguous" argument to skip unphased segments of the genome. Results were scaled to real time by applying a mutation rate of $1.25\times10^{-8}$ per site per generation and a generation time of 29 years.

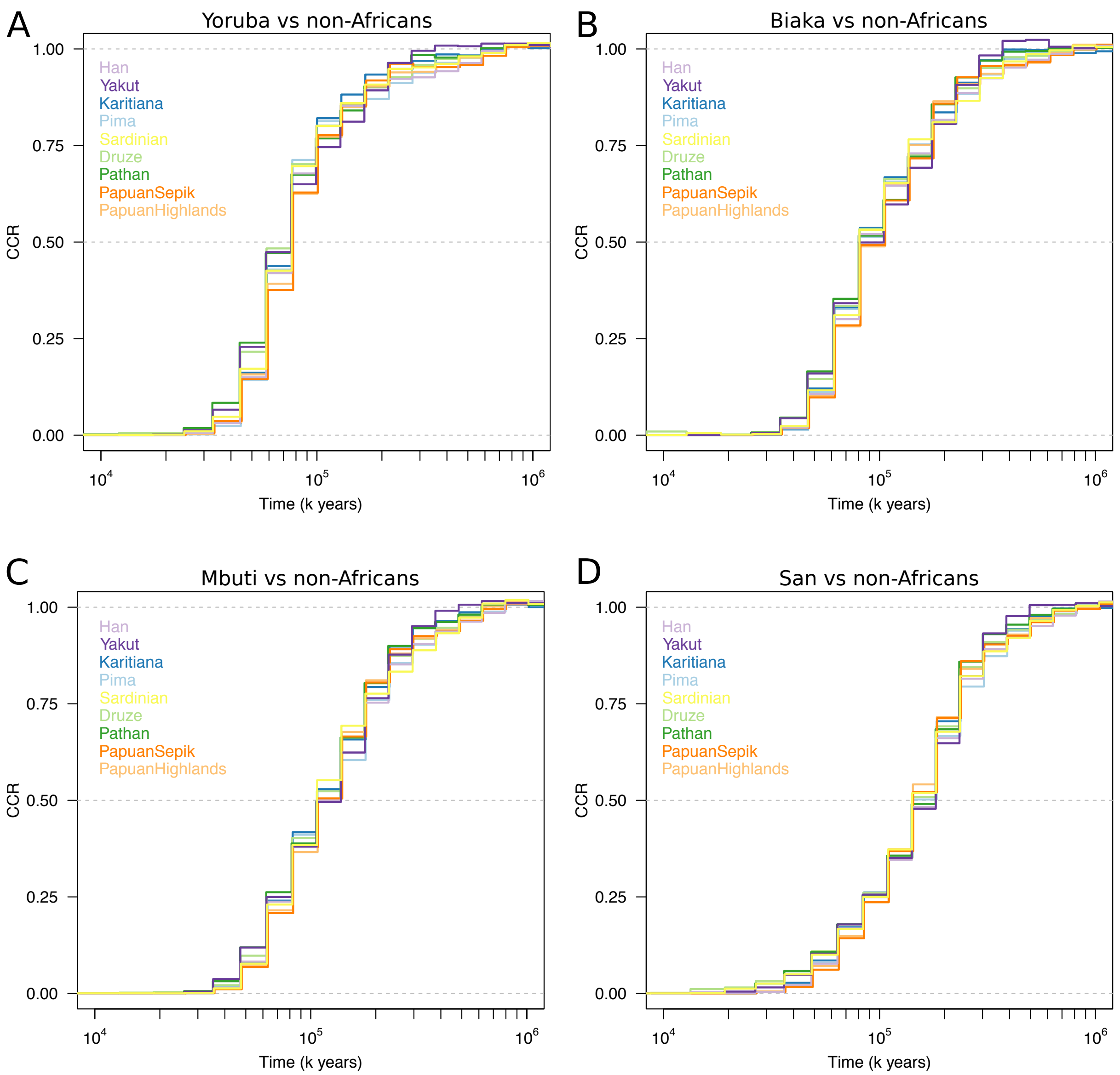


***Figure S5: MSMC2 analyses of divergences between African and non-African populations.*** *Curves were computed using 4 physically phased haplotypes per population. A) Yoruba versus non-African populations. B) Biaka versus non-African populations. C) Mbuti versus versus non-African populations. D) San versus non-African populations*

Clean split scenarios were simulated using scrm (*53*) version 1.7.2, using a mutation rate of $1.25\times10^{-8}$ per site per generation, a generation time of 29 years, a recombination rate of $1.12\times10^{-8}$ per site per generation, an initial $N_e$ of 20,000, a doubling of $N_e$ between 300 and 1000 kya and simulating 14 chromosomes each of size 150 Mbp (for a total genome size of 2.1 Gbp, similar to the size of the empirical genomes restricted to the accessibility mask). The average heterozygosity of the resulting sequences was 0.986 per kbp, similar to human genomes of African ancestry. The sequences were analysed with MSMC2 as above, except the --fixedRecombination parameter was applied (running without this resulted in non-monotonic curves for many simulated histories).

We performed additional MSMC2 runs to test if the deep structure observed in our empirical results, i.e. that the cross-coalescence curves remain below 1 even several hundreds of thousands of years ago, could be caused by batch effects associated with sequencing and processing the two haploid genomes from a diploid human sample together. Any process that could cause the genotypes of these two genomes to appear artificially similar, from somatic or cell-line loss of heterozygosity events to under-calling of heterozygous genotypes during data processing, could cause the time to coalescence between these two genomes to appear lower than that to genomes from another individual. We ran MSMC2 on four haplotypes, but in each of the two populations selecting one haplotype from two different individuals, using the “-I” command line syntax of MSMC2 v2.1.1 on input files prepared with eight haplotypes, e.g. “-I 0,2”, “-I 4,6” and “0-4,0-6,2-4,2-6” for the within first population, within second population and between population runs, respectively. We thus use four haplotypes sequenced in four different individuals, and should avoid any batch effects of the type described above. The curves obtained from these runs in many cases tend towards slightly higher relative cross-coalescence values in deep time periods then when using haplotypes sequenced in the same individual, and more so when involving two non-African populations, but they still display largely the same behaviour (Figure S6). While it’s possible that the magnitude of the deep structure exhibited in our results could thus be slightly exaggerated by technical artefacts of this nature, they are unlikely to be responsible for the whole effect.

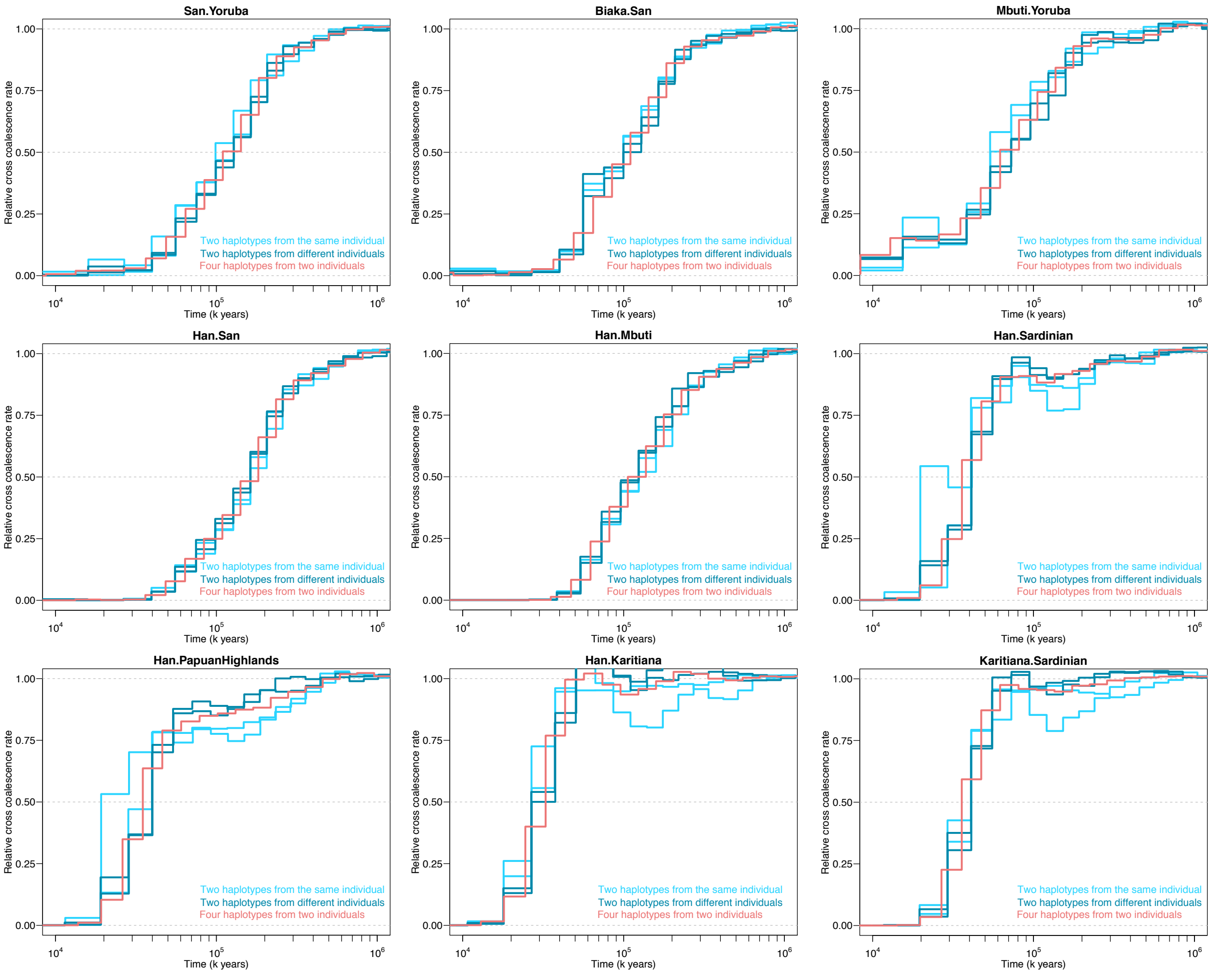


*__Figure S6: Testing diploid batch effects in MSMC2.__ Curves are displayed that use different numbers and configurations of haplotypes from each of the two populations. The “four haplotypes from two individuals” curves are from the standard runs that are also displayed in Figure 5 (a total of eight haplotypes in the run across the two populations). The “Two haplotypes from the same individual” curves use two haplotypes from one individual from each population (a total of four haplotypes in the run). The “Two haplotypes from different individuals” curves also use two haplotypes from each population (a total of four haplotypes in the run), but from two different individuals. For these latter two runs with four haplotypes, there are two independent replicate curves using different pairs of individuals.*

#### *Application of MSMC2 to archaic genomes*

We also ran MSMC2 on pairs of modern human and archaic populations. We used the high-coverage Altai Neanderthal, Vindija Neanderthal and Denisovan genomes, left all heterozygous genotypes in these as unphased, and ran MSMC2 as above except using only two haplotypes per population (we also performed runs without the "–skipAmbiguous" argument but found this made little difference to the results). While the method relies on phased genotypes for the between population coalescence rate estimation, the low heterozygosity of the archaic genomes means that many segments of these genomes will be homozygous and thus by necessity phased, which might be the reason we obtain seemingly sensible results.

We plotted the results naively as above, but also made plots where we attempted to correct for the fact that the archaic genomes are from ancient remains that stopped accumulating mutations at their time of death several tens of thousands of years ago. MSMC2 reports the within and the between population coalescence rates separately, and we could thus adjust these as a function of the sample age before calculating the relative cross-coalescence rates. We shifted back the within population coalescence rates by adding a new time segment between 0 and the sample age with a rate of 0 and then adding the sample age to all existing time segments. We similarly shifted back the between populations coalescence rates by half the age of the sample age. We used sample ages of 122,000 years for the Altai Neanderthal, 52,000 years for the Vindija Neanderthal and 72,000 years for the Denisovan genome (*19*). Overall, we found that these sample age adjustments did not have substantial effects on the results, especially not the timing of the separation between modern and archaic humans on the order of 500 kya. They did however shift backwards in time the archaic admixture signal in non-African genomes such that it peaks around 80-120 kya, rather than 40-80 kya, when using the Vindija Neanderthal genome – this might actually more accurately reflect the timing of the event, as the introgressing Neanderthal population has some level of divergence from the Vindija individual (*19*) such that the coalescence events with the introgressed haplotypes will be older. The curves obtained when using the Altai Neanderthal are shifted backwards in time relative to when using the Vindija Neanderthal, reflecting how the former is further diverged from the introgressing source.

We observe the MSMC2 Neanderthal gene flow signal in all non-African genomes and to a much-reduced degree in Yoruba. The most likely explanation for these results is non-African gene flow carrying Neanderthal ancestry into West Africa (*37*). The other African populations do not display the same behaviour, with the tiny deviations from zero relative cross-coalescence rate in recent time periods, especially so when shifting back rates by sample age (including when using the Denisovan genome), likely not being distinguishable from technical noise. We do not observe any clear recent gene flow signal when running the Denisovan genome against African, Eurasian or Native American genomes (a tiny increase in the Yakut is difficult to distinguish from technical noise). However, when running Denisovan against Oceanian genomes (PapuanHighlands and PapuanSepik) a subtle upwards shift in the curve is visible roughly in the time span between 140 and 400 kya (moving back by ~40 kya when shifting rates by the Denisovan sample age). This very likely reflects the Denisovan gene flow in these populations, with the large backwards shift in time of the signal reflecting how the sequenced Denisovan from the Altai mountains is highly diverged from the population that contributed to Oceanians (*15*), such that the coalescence events with the introgressed haplotypes in Oceanians are quite old.

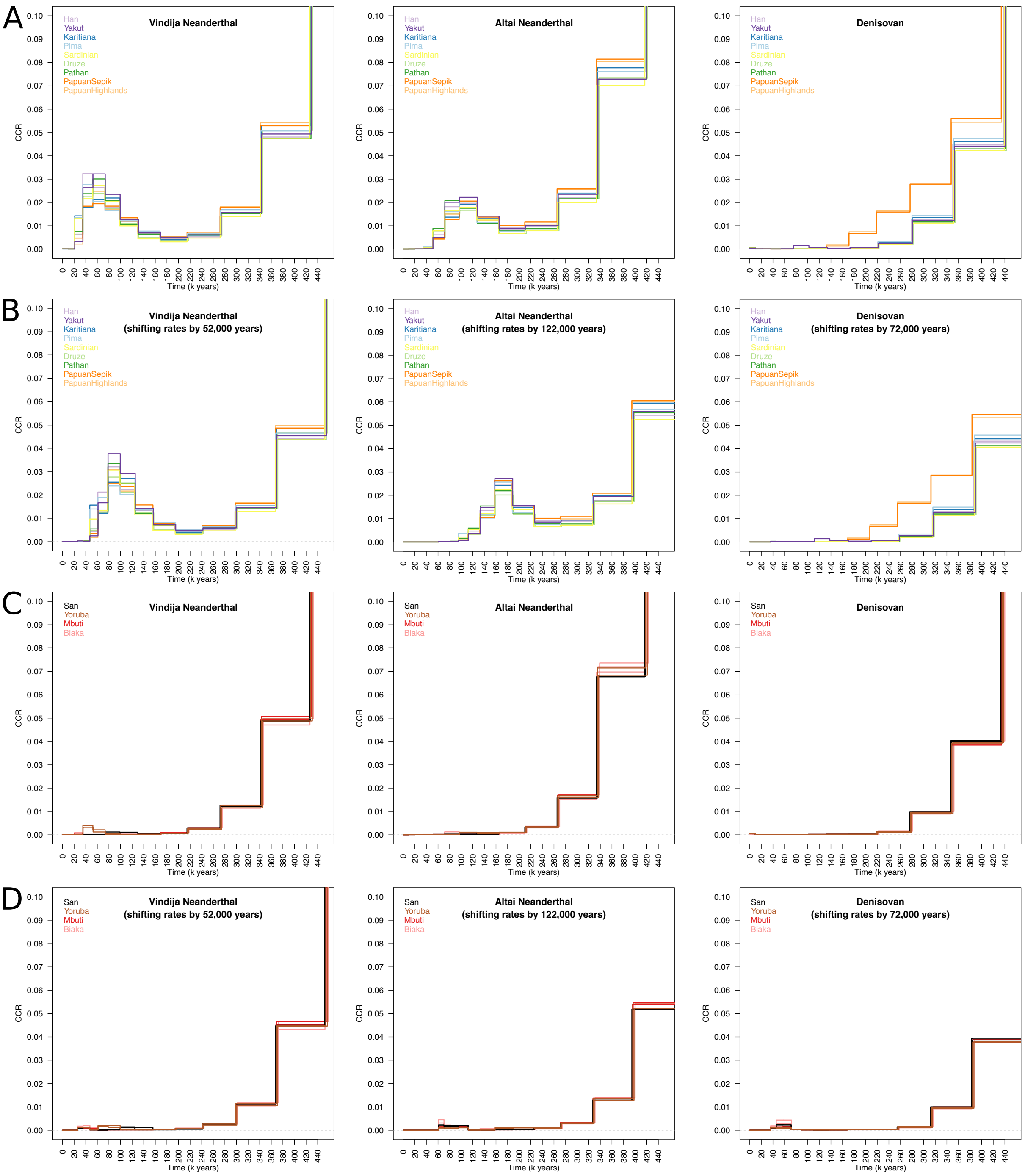


*Figure S7: Signals of archaic gene flow in MSMC2 curves. Results of cross-population MSMC2 runs, zooming in on the signal of Neanderthal genome flow in modern human genomes (note the highly reduced range of the vertical axis). A) Archaic genomes against non-African genomes. B) Archaic genomes against non-African genomes, attempting to adjust for the age of the archaic specimens. C) Archaic genomes against African genomes. For each African population, curves for two different individuals are displayed in the same colour. D) Archaic genomes against African genomes, attempting to adjust for the age of the archaic specimens.*

The application of MSMC2 to archaic genomes thus provides an additional line of evidence for the known archaic admixture in non-Africans, as well as for small amounts of Neanderthal ancestry in West Africans. We note that a conceptually similar approach, identifying haplotypes in non-Africans with low absolute divergence to archaic genomes, was used previously (*15*). Importantly, while methods relying on allele frequency correlations, e.g. D or $f_4$-statistics, produce results that are always relative between pairs of modern human populations, these MSMC2 results involve just one modern human genome at the time without the need for any baseline assumptions. These results thus allow us to say not only that most sub-Saharan African groups have less Neanderthal ancestry than non-Africans, but also that they most likely in an absolute sense have very little Neanderthal ancestry, e.g. in the case of Mbuti a level that, within the limits of resolution of the method, likely is compatible with no Neanderthal ancestry at all.

*Site frequency spectrum models with momi2*

We used the momi2 software (*29*), which fits models to the site-frequency spectrum, to estimate pairwise split times between all 1431 combinations of the 54 populations, assuming a simple clean split without subsequent gene flow and a mutation rate of $1.25\times10^{-8}$ per site per generation, using ancestral allele information from the Ensembl EPO alignments, and with confidence intervals obtained through bootstrapping across 500 genomic blocks. While the clean split assumption will be unrealistic in many cases, as evidenced by our MSMC2 results, the overall correlation between these momi2 split time estimates and the MSMC2 midpoint estimates is quite high ($r = 0.93$), suggesting the former will provide a decent approximation of the latter without requiring phased haplotypes. However, we also observed that the momi2 estimates are affected by the sample size of a population, and that they clearly greatly underestimated split times involving Native American populations (for example, some Native American against East Asian split estimates at just a few thousand years), perhaps as an artefact of the low recent effective size of these populations. We therefore do not place much emphasis on any particular single estimates. More elaborate models incorporating multiple populations, post-split gene flow and archaic admixture in the history of non-African populations have been shown to provide more accurate split time estimates (*29*).

### Effective population size histories

We used smc++ v1.12.1 (*23*) to estimate effective population size histories for each of the 54 populations separately. Input files were prepared from genotypes on all called sites, masking out indel alleles and any third or further minor alleles at multi-allelic sites by setting the genotype of any individual carrying such alleles to missing. In addition to sites falling outside the accessibility mask, sites not present in the input VCFs were masked out from the analyses. smc++ requires one individual in the run to be specified as the "distinguished" individual, forming the basis of the coalescence time element of the inference, with the remaining individuals only contributing allele frequency information. For each population, we established a ranking for which individuals to use the distinguished individual, firstly prioritizing individuals not listed as ancestry outliers relative to their population by (*13*), secondly prioritizing Sanger PCR-free, then Sanger PCR, then SGDP PCR-free and then SGDP PCR-free libraries, and thirdly prioritizing higher sequencing coverage.

We ran smc++ assuming a mutation rate of $1.25\times10^{-8}$ per site per generation, inferring effective population size up until 34 generations (approximately 1000 years assuming a generation time of 29 years) ago and otherwise default settings. For each run we prepared six alternative input files with different individuals as the distinguished individual and then gave all of these as input, resulting in composite likelihood results.

Some populations display substantial decreases in inferred effective population size in the last ~5000 years, but we suspect that in many cases this reflects very recent endogamy or bottlenecks, the effects of which are being spread out over a larger time interval by the inference. We also noticed that under some parameter settings, in particular when decreasing the strength of the regularization (allowing more flexible curves to be fit, but also with a greater risk of overfitting) or when stopping the inference at 172 instead of 34 generations, Native American groups are inferred to have experienced dramatic growth approximately in the period between 20 and 10 kya. This is not observed when running on the parameter settings above, but we speculate that this could be a real signal which is counteracted in the inference by the strong bottlenecks experienced in the very recent time by the analysed Native American groups. Rapid population growth at in this time period, coinciding with the initial peopling of the American continents, would be consistent with observations from the mitochondrial and Y-chromosomal phylogenies, which both display dramatic star-like behaviour starting around 15 kya indicative of rapid population expansion at this time. While the large degree of variability in the inferred curves between SMC++ parameter settings means we cannot be highly confident that the inferred growth reflects real growth rather than an artefact, we do not observe the same behaviour in other populations under the same settings, demonstrating that at least the phenomenon is a function of Native American genomes specifically rather than simply inherent to the inference method itself.

For the 13 populations for which we had physically phased genomes, we also inferred effective population sizes histories using MSMC2. We ran MSMC2 on the two diploid genomes (four haplotypes) from each of these populations with the "--skipAmbiguous" argument to skip unphased segments of the genome. The MSMC2 results obtained on these physically phased genomes were largely concordant with the SMC++ results (Figure S8), confirming broad observations including recent declines in the African hunter-gatherer groups Mbuti, Biaka and San. We also ran MSMC (*21*), and those curves were also very similar (not shown).

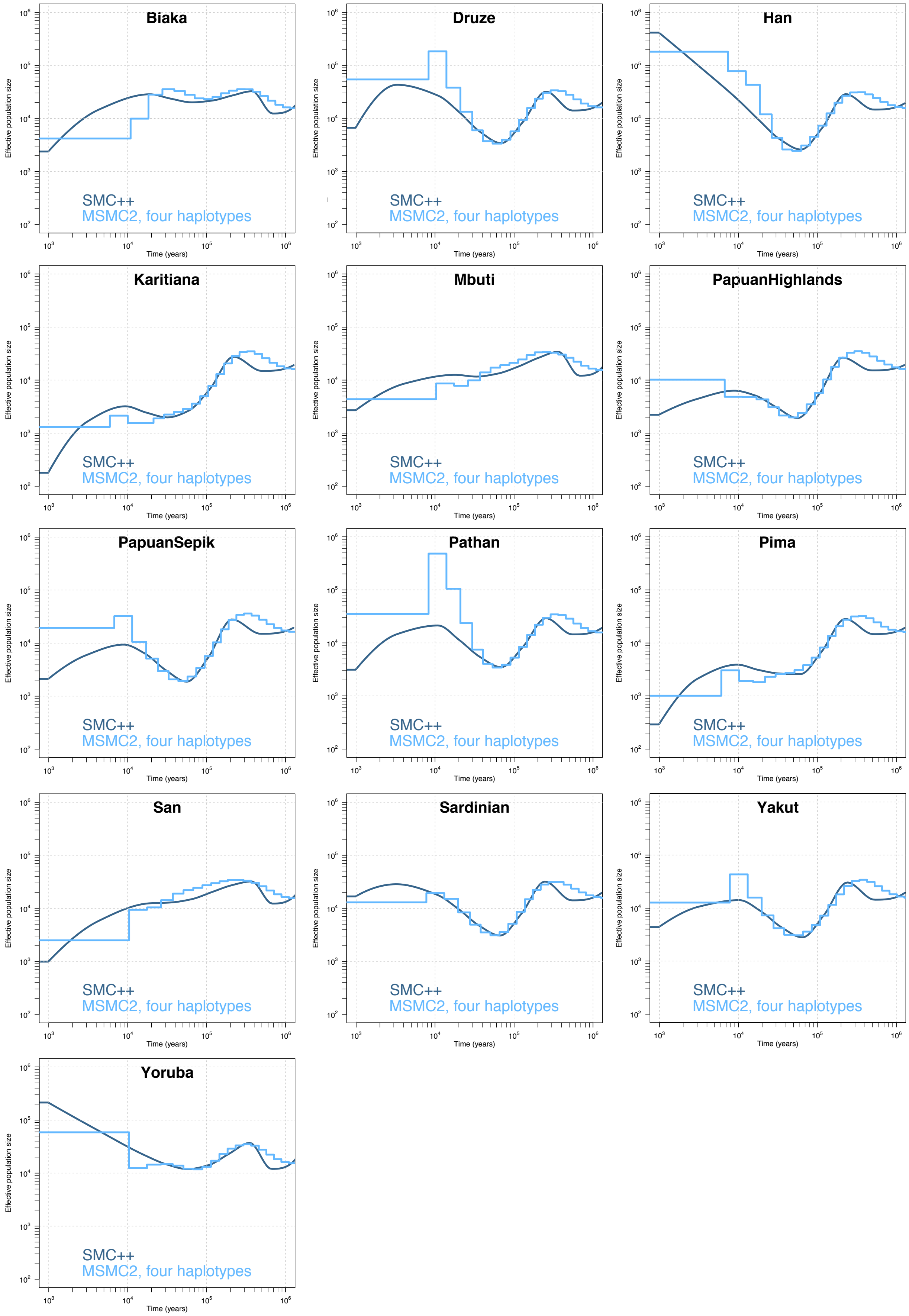


***Figure S8: Comparison of effective population size histories inferred using SMC++ and MSMC2****. The SMC++ runs used all individuals from the given population, across six alternative choices of the distinguished individual so as to produce composite likelihoods. The MSMC2 runs used two physically phased genomes (four haplotypes) per population.*

Y chromosome analyses

The haploid genotype calls for 603 males (here excluding the Meyer libraries which were of lower quality) across 10.3 Mb of accessible regions on the Y chromosome (*54*), lifted over to GRCh38 using the UCSC liftOver tool, were extracted using bedtools v2.22.0 (*55*). Within these regions, sites were filtered out if they contained indels, had missing genotype calls in more than 5% of the samples, had genotype qualities below 30 in more than 5% of the samples or had coverage above twice or below a third of the sample mean for more than 5% of the samples. This left a final set of 52,032 variant and 10,016,322 invariant sites.

The haplogroup of each sample was predicted using the yHaplo software (https://github.com/23andMe/yhaplo) after substituting the marker coordinates in the relevant input files to correspond to the GRCh38 assembly. An initial maximum likelihood phylogenetic tree was constructed using RAxML (v8.2.10) (*56*) with the GTRCAT substitution model with the set of 52,032 variant sites and then used as a starting tree for dating with BEAST v1.7.2 (*57, 58*). Markov chain Monte Carlo samples were based on 11,000,000 generations, logging every 1,000 generations. The first 10% of generations were discarded as burn-in. Eight independent runs were combined using LogCombiner. A constant-sized coalescent tree prior, the HKY substitution model, accounting for site heterogeneity (gamma) and a strict clock with a substitution rate of $0.76\times10^{-9}$ (95% confidence interval: $0.67\times10^{-9}$ to $0.86\times10^{-9}$) single nucleotide mutations per bp per year (*59*) was used. A prior with a normal distribution based on the 95% confidence interval of the substitution rate was applied. Only the variant sites were used, but the number of invariant sites was defined in the BEAST xml file. A summary tree was produced using TreeAnnotator v1.8.1. The final tree (Figure S9) was visualised using the FigTree software (http://tree.bio.ed.ac.uk/software/figtree/).

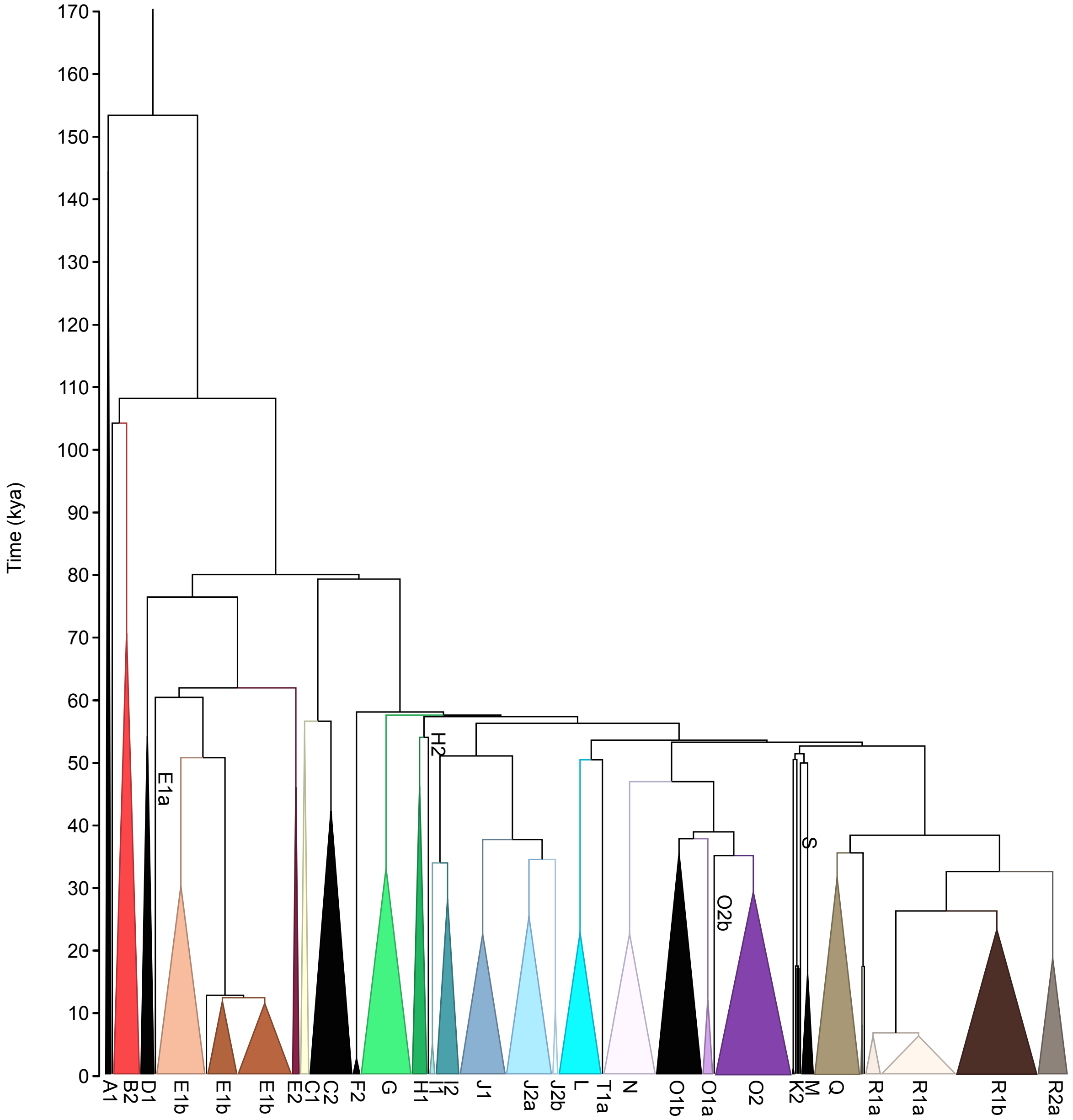


***Figure S9: Y-chromosomal phylogeny**. Branch lengths are proportional to the estimated times between splits. Coloured triangles represent collapsed major clades, with the width of the triangles proportional to the number of samples in each clade.*

### Archaic introgression

Genotypes for the high-coverage Vindija Neanderthal (*19*), Altai Neanderthal (*15*) and Denisovan (*6*) individuals on the GRCh37 reference assembly, along with corresponding filter files, were obtained from ftp.eva.mpg.de/neandertal/. The genomic coordinates were lifted over to GRCh38 using CrossMap v0.2.5 (*60*).

#### *Global estimates of archaic ancestry proportions*

To estimate global proportions of archaic ancestry, we constructed a VCF which contained genotypes for the high-coverage Vindija Neanderthal, Altai Neanderthal and Denisovan genomes not just at sites that are polymorphic among modern human genomes, but also including sites that are monomorphic among modern humans but polymorphic among the entire set of modern and these three archaic genomes. We used these files, restricted to the accessibility mask, to estimate:

- **The proportion of Neanderthal ancestry in Eurasians**: We used an $f_4$-ratio, assuming a baseline of no Neanderthal ancestry in Mbuti, calculated using popstats (*61*), with the "--informative" option to include only variants polymorphic on both sides of the $f_4$ statistic:

$$\frac{f_4(Chimpanzee, Altai\ Neanderthal; X, Mbuti)}{f_4(Chimpanzee, Altai\ Neanderthal; Vindija\ Neanderthal, Mbuti)}$$

  The statistic gives mean estimates of 1.18% for Middle Eastern, 2.09% for Central & South Asian, 2.14% for European, 2.26% for American and 2.24% for East Asian populations. This statistic will not provide accurate estimates for Oceanian populations because of their high Denisovan ancestry. The highest population estimate is for the Chinese Xibo population at 2.41% (95% CI: 2.12 – 2.70%). Within Africa, the estimates are 0.048% (95% CI: -0.061 – 0.16%) for Biaka, 0.13% (95% CI: -0.025 – 0.28%) for San, 0.16% (95% CI: 0.031 – 0.29%) for Mandenka, 0.16% (95% CI: 0.035 – 0.29%) for BantuKenya, 0.16% (95% CI: 0.035 – 0.29%) for BantuSouthAfrica and 0.18% (95% CI: 0.05 - 0.30%) for Yoruba.

- **The proportion of Denisovan ancestry in Oceanians**, in two different ways:
  1) Using an $f_4$-ratio, assuming a similar level of Neanderthal ancestry in PapuanHighlands as in Han (*32*), calculated using popstats with the "--informative" option to include only variants polymorphic on both sides of the statistic:

$$\frac{f_4(Mbuti, Vindija\ Neanderthal; Han, PapuanHighlands)}{f_4(Mbuti, Vindija\ Neanderthal; Han, Denisovan)}$$

     This gave an estimate of 2.86% (95% CI: 2.08 - 3.64%).

  2) Using the qpAdm method (*62*) in ADMIXTOOLS version 5.0 (*13*). A model with PapuanHighlands as target, Karitiana and Denisovan as sources, and Yoruba, Altai Neanderthal, Chimpanzee, Vindija Neanderthal, Mbuti and Biaka as outgroups gives an estimate of 2.70% (95% CI: 2.11% - 3.29%), with a model fit p-value of 0.1892. The rationale behind this approach is to simply model PapuanHighlands as a two-way mixture between Denisovan and non-African ancestry, and we here use Karitiana as a representative of the latter. It's likely that Karitiana has a small but non-zero amount of Denisovan ancestry, which would bias the estimate downwards, but given the very low amount of this ancestry the bias should be very small. Using a West Eurasian population with likely zero levels of Denisovan ancestry in place of Karitiana in the model would not be ideal as the lower levels of Neanderthal ancestry in such populations compared to Oceanians would lead to bias.

     If we take the midpoint of these two estimates and conservatively take the largest confidence intervals, we obtain an ancestry fraction of 2.78% (CI: 2.08 - 3.64%)

     In summary, these estimates of the proportion of Denisovan ancestry in Oceanians are largely consistent with previous f-statistics based estimates, but lower than some of the highest estimates: 4.8% (*31*), 3.0% (*6*), 3.5% (*63*), 3.4% (*32*), 3.2% (*33*). The first of these estimates was made using a low-coverage Denisovan genome sequence and with lower quality modern data, and is thus likely not very reliable. A proportion of around or, as suggested by our new estimates here, even slightly below 3.0%, thus seems probable.

- **The proportion of Neanderthal ancestry in Oceanians:** Due to the high levels of both Neanderthal and Denisovan ancestry in Oceanians, and the partially shared drift between these two archaic groups, it is difficult to obtain an unbiased estimate of Neanderthal ancestry in Oceanians using a simple $f_4$-ratio. To get around this, we tried to estimate both Denisovan and Neanderthal ancestry jointly using qpAdm. We tried a model with PapuanHighlands as target, Denisovan, Vindija Neanderthal and Yoruba as sources and Chimp, Altai Neanderthal, Mbuti and San as outgroups, but this did not fit the data ($p = 4\times10^{-36}$), likely due to the model not accurately representing the relationships among the African groups and the position of Yoruba as a proxy for non-African ancestry. An East African population would likely be a better proxy, however the HGDP dataset does not contain a suitable population. We therefore tried this analysis on the Simons Genome Diversity Project (*3*) dataset which contains the Dinka population. A model with Papuan as target, Dinka, Vindija Neanderthal and Denisovan as sources and Chimpanzee, Altai Neanderthal, Yoruba, Mende, Biaka and Mbuti as outgroups fits the data ($p = 0.061$) and gives estimates of 2.0% (95% CI: 1.41 - 2.59%) Neanderthal ancestry and 3.4% (95% CI: 2.42 - 4.38%) Denisovan ancestry in Papuans. While the exact estimates might still be sensitive to slight model violations, these results suggest that the level of Neanderthal ancestry in Papuans is similar to the levels in other non-Africans.

*Implementation of a Hidden Markov Model for archaic haplotype detection*

A hidden Markov model (HMM) was used to detect introgressed segments from the Neanderthal and Denisovan populations in modern human genomes. The HMM decodes segments of haploid genomes (thus requiring phased haplotypes) into two hidden states: unadmixed (0) and archaic (1). We summarize the observed data by the pattern of allele sharing between a panel of sub-Saharan African genomes, the haploid genome under examination, and a panel of one or more archaic genomes. Only informative sites where a derived allele is shared between two groups and absent in the third are considered by the HMM. For convenience, the three informative combinations are encoded as emission types 1, 2 and 3 (Table S4). If a genetic segment entered the modern human population from an archaic hominin population relatively recently, it should share more derived variants with the archaic genomes. Since sub-Saharan Africa is assumed to have no or very small amounts of Neanderthal and Denisovan ancestry, an allele shared between African and non-African genomes would most likely have arisen on a lineage within the modern human population. Incomplete lineage sorting in modern segments might cause some African lineages to coalesce first with the archaic genomes, thus sharing a derived allele unseen in a non-African genome. Such observations are less likely to occur in the archaic segments due to the small effective size of the archaic populations.

***Table S4: Encoding of HMM emission types***. *A value of "0" in the genotype column means ancestral allele in all genomes in the panel, while "1" means derived allele in at least one genome*

| **Genotype** | | | **Emission type** |
|---|---|---|---|
| Sub-Saharan African panel | Sample of interest | Archaic panel | |
| 1 | 0 | 1 | 1 |
| 1 | 1 | 0 | 2 |
| 0 | 1 | 1 | 3 |

Transitions are only allowed between informative sites. Because the distance between informative sites is not constant, the transition probabilities are updated on-the-fly using genetic distance:

$$T = \begin{bmatrix} 1-(1-e^{-dt})\cdot\alpha & (1-e^{-dt})\cdot\alpha \\ (1-e^{-dt})\cdot(1-\alpha) & 1-(1-e^{-dt})\cdot(1-\alpha) \end{bmatrix}$$

Where $T_{ij}(i,j \in \{0,1\})$ is the probability to transit from state $i$ to state $j$, $t$ the time since admixture, $d$ the genetic distance either extrapolated from a genetic map or, in the absence of such a map, using the physical distance and a constant per-site recombination rate. The admixture proportion $\alpha$ also defines the initial state probabilities $\pi = (1-\alpha, \alpha)$. The full model is specified by $t, \alpha$ and the emission probabilities matrix $E$.

*Model training*

Since the HMM is designed to work with a single archaic source at the time, we simulated 20 haploid genomes under the demographic model in (Figure S10) using msprime (*64*), and obtained the maximum likelihood estimation (MLE) of model parameters based on the true underlying state of the genetic segments (Table S5). We fixed $t$ at the true value of 2,000 but noted that varying it between 1,000 and 4,000 while keeping the other parameters unchanged has little influence on the decoding results. Applying this model to simulated data recovered 90.74% of the true archaic segments, with a false discovery rate of 3.68%.

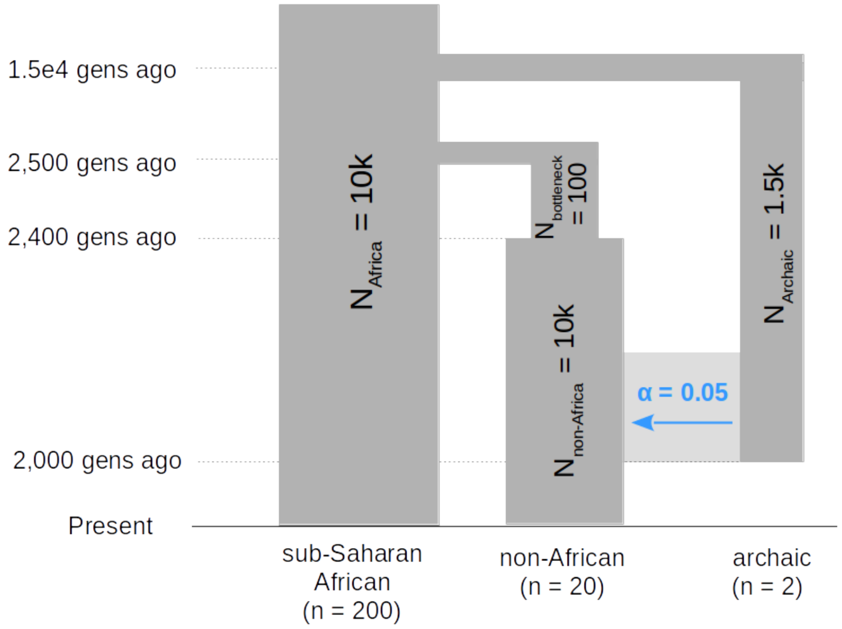


***Figure S10: Demographic model underlying simulations with only one source of archaic gene flow***

We also explored the Baum-Welch algorithm and numerical likelihood optimization to train the HMM, however both training methods sometimes produced a minor state that absorbs type 2 emissions. It appears difficult to establish an archaic and a modern state from unsupervised training.

*Table S5: HMM parameters estimated from simulations.*

| | |
|---|---|
| Initial state distribution ($\pi$) | $[0.9655 \quad 0.0345]$ |
| Emission matrix ($E$) | $\begin{bmatrix} 0.1777 & 0.8208 & 1.336\times10^{-3} \\ 1.691\times10^{-3} & 2.015\times10^{-4} & 0.9981 \end{bmatrix}$ |

*Comparison with published methods*

In a scenario where only one archaic source of introgression is concerned, we compared the performance of the HMM in detecting Neanderthal segments with the S* method (*33, 65*), which searches for long haplotypes in linkage disequilibrium unseen in the African panel, and a method using a conditional random field (CRF) (*66*), which examines allele sharing, haplotype divergence and local recombination rate. Since the implementation of neither method is publicly available, we compared the result of running CRF, S* and the informative-site-only HMM on the same set of genomes instead. The Neanderthal segments detected by CRF and S* in individuals from 1000 Genomes Project were downloaded from the authors' websites. The HMM was then run on chromosome 1 of the 544 individuals that were included in both studies and the Viterbi sequences were obtained. To be consistent with the other two methods, only the high-coverage Altai Neanderthal genome (*15*) was used in the archaic panel. The highest agreement is between HMM and S* (Figure S11A), where the shared regions constitute 72.84% of the total material recovered by the HMM and 82.43% of that recovered by S*. It is worth noting that although the S* score itself does not rely on the archaic genome, the reported segments in (*33*) have undergone subsequent filtering based on their match score to the archaic genomes. The other pairwise comparisons between methods only show around 40% reciprocal overlap. The overlap patterns are consistent across the seven Eurasian populations analyzed.

Under a different metric, a segment detected by one method is treated as a match if at least half of it is also reported by another method. Although the HMM does not preferentially detect or miss segments of particular lengths compared to the two other methods, segments detected by the HMM but not by the other methods tend to be shorter (Figure S11B). Segments shorter than 50 kB constitute 91.22% of those not detected by S* and 82.37% of those not detected by CRF. Most segments longer than 50 kB are reported by all three methods.

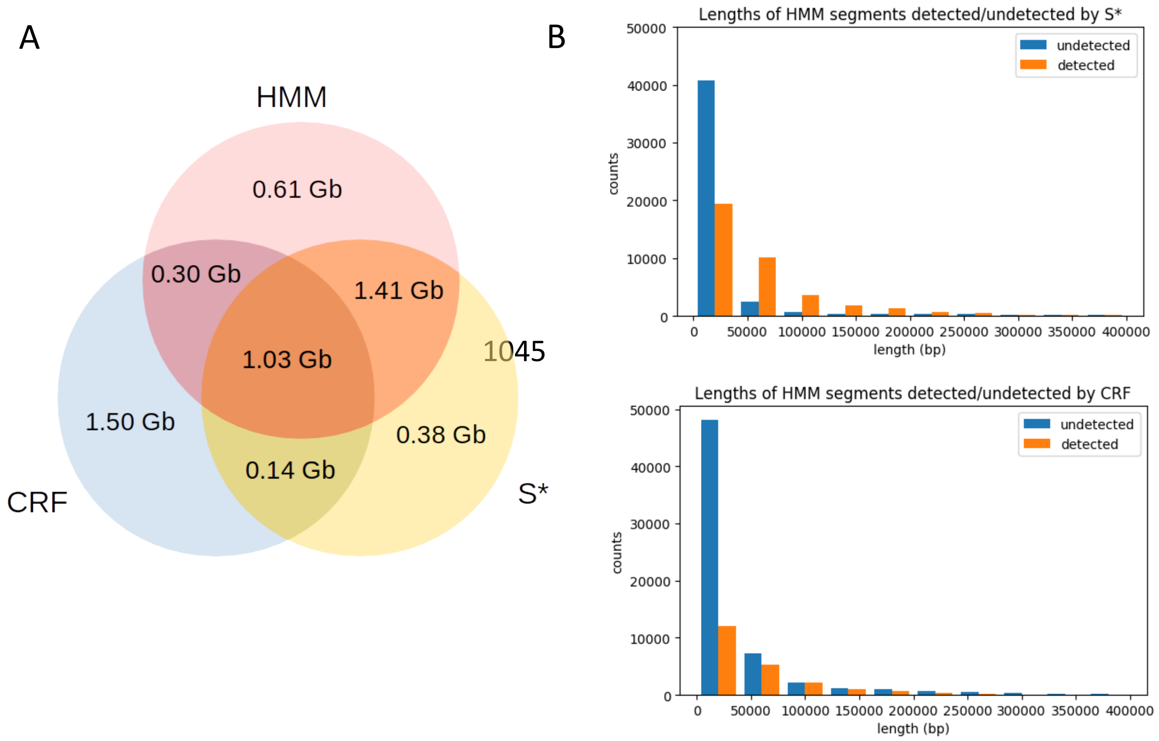


***Figure S11: Comparison of HMM for archaic haplotype detection to other methods. A)*** *Amount and overlaps of Neanderthal segments identified on chromosome 1 of 544 individuals from the 1000 Genomes Project B) Lengths of Neanderthal segments detected by the HMM and detected/undetected by other methods.*

*Distinguishing sources of archaic segments*

Since Neanderthal and Denisovan segments coexist in many non-African genomes, we also tested various methods to distinguish between them. We launched another set of simulations following the demographic model in Figure S12, where a non-African population received 3% gene flow from the Neanderthal population, followed by 1% from the Denisovan population. The two-state HMM can be extended to include a Neanderthal state, a Denisovan state and a modern state, but this model showed very high false discovery rate (i.e. mislabeling unadmixed segments as archaic) on simulated data. Instead, we ran the two-state HMM twice with the same model parameters, first with two Neanderthal genomes in the archaic panel, then with the Denisovan genome. The posterior probabilities of the archaic state at each informative site from both runs, $p_N$ and $p_D$, were used to assign them into the following categories:

- If $p_N \leq 0.5$ and $p_D \leq 0.5$, tagged as "modern";
- If $p_N > p_D$ and $p_D < 0.8$, tagged as "Neanderthal";
- If $p_D > p_N$ and $p_N < 0.8$, tagged as "Denisovan";
- If $p_N \geq 0.8$ and $p_D \geq 0.8$, tagged as "ambiguous archaic".

If a site only showed up as informative regarding the Neanderthal run or the Denisova run, the missing $p_D$ or $p_N$ was extrapolated linearly by physical distance from adjacent sites. The Neanderthal, Denisovan and ambiguous archaic segments were identified by linking neighbouring sites in the same category. We found that this approach reduces the false discovery rate in simulations to around 0.05, at the cost of pooling a large proportion of archaic segments into the ambiguous category. The probability of labeling true Neanderthal segments as Denisovan is also below 0.05, and even lower the other way around. The criteria for assigning the categories based on $p_N$ and $p_D$ can also be adjusted according to the needs of downstream analyses, reflecting a trade-off between type 1 and type 2 errors. In analyses involving the haplotypes of Neanderthal and Denisovan segments, we used a more stringent set of criteria to obtain a "strict" set of archaic segments:

- If $p_N \geq 0.8$ and $p_D < 0.5$, tagged as "Neanderthal";
- If $p_D \geq 0.8$ and $p_N < 0.5$, tagged as "Denisovan".

In simulated data, this reduced the proportion of true Neanderthal segments to 0.016 among predicted Denisovan segments, and the proportion of true Denisovan segments to 0.0013 among predicted Neanderthal segments.

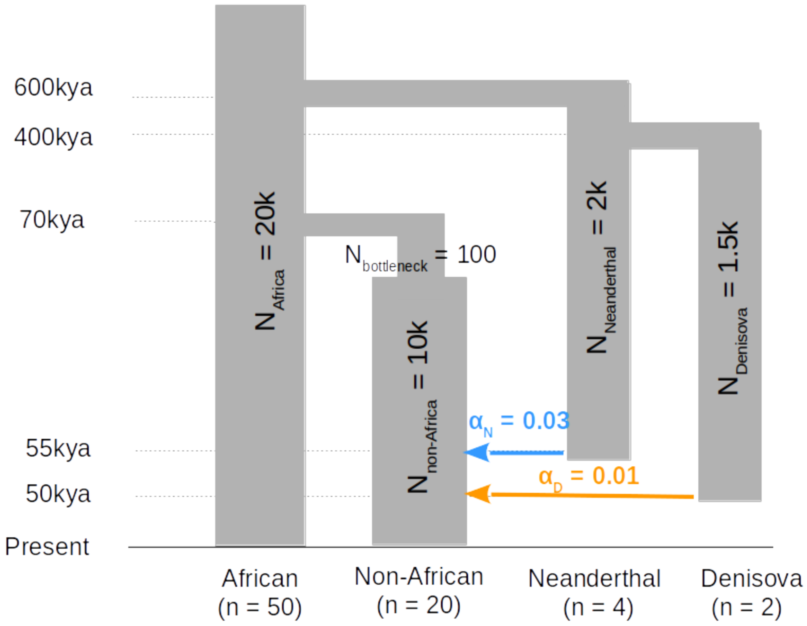


*Figure S12: Demographic model underlying simulations with both Neanderthal and Denisovan gene flow.*

*Detecting Neanderthal and Denisovan segments in modern populations*

The two-state HMM was run twice on the 929 phased HGDP genomes with a genetic map to obtain the posterior probabilities of being in the Neanderthal and Denisova state at all informative sites. All 104 genomes from sub-Saharan Africa were included in the African panel, but when extracting observations, we allowed the archaic allele to reach a maximum frequency of 0.01 in this panel to allow for small amounts of archaic ancestry within Africa. Two high-coverage Neanderthal genomes, one from Denisova cave (*15*) and the other from Vindija cave (*19*), were used in the archaic panel in the Neanderthal run whilst the Altai Denisovan genome (*6*) was used in the Denisova run. In addition to the HGDP accessibility mask, we also included a low complexity regions mask (*67*). All sites that do not pass the masks were ignored as non-informative in the HMM runs. We used the ancestral sequences from Ensembl EPO alignment to determine the ancestral state; and in case of unknown sites in this panel, assumed the genotype in chimpanzee (Pan_tro 3.0) to be ancestral. Only sites that are polymorphic in the HGDP dataset were retained after merging. In effect, this leaves out derived sites shared by all modern human genomes but not the archaics, which are type 2 emissions (Table S4) that supports assigning the modern state. However, such sites should be very rare in the genome, as derived alleles shared by all modern humans will be older than 200k years.

We obtained two sets of archaic segments following the first (hereafter the "basic" set) and the second (hereafter the "strict" set) criteria described in the previous section. In practice, the "basic" set assigns less material to the ambiguous category than in simulation studies, such that the actual distinguishing power is expected to be greater than in the results in the simulations.

*Geographical distribution of archaic ancestry*

Figure S13A compares the average amount of Neanderthal, Denisovan and ambiguous segments identified in the HGDP genomes (from the "basic" set) by geographical region, and Figure S13B shows the mean and standard deviation of the amount of Neanderthal and Denisovan segments identified in each population. The length of masked regions was excluded in all plots.

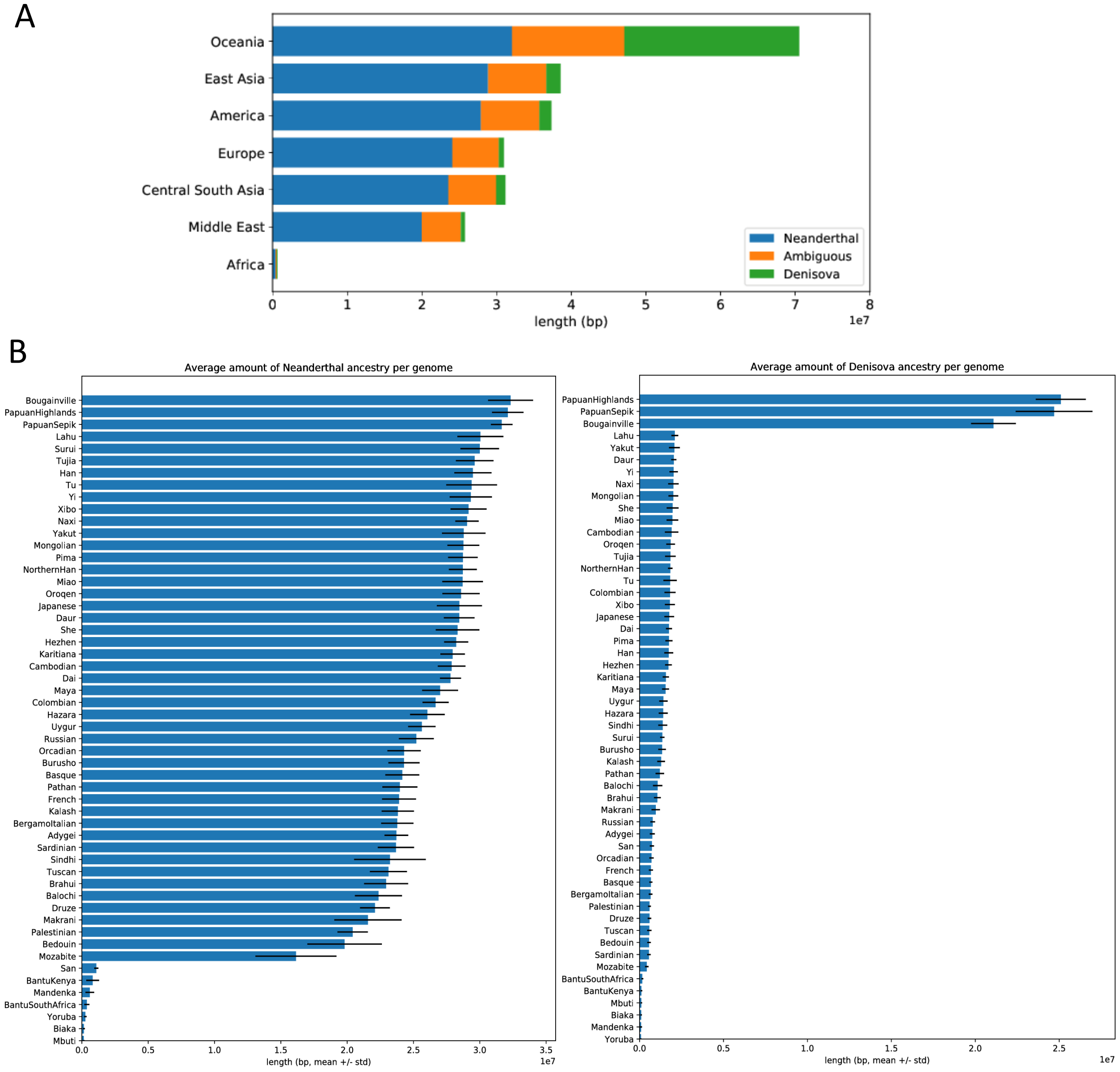


***Figure S13: Amounts of archaic segments identified.*** *A) Average amount of archaic segments identified per genome by geographical region. B) Average amount of Neanderthal and Denisovan segments identified per genome, by population.*

Very few archaic segments are detected in sub-Saharan African populations, but this is expected on technical grounds alone as our method conditions on allele frequencies in these populations. In accordance with previous studies, the amount of Neanderthal ancestry is higher in East Asia and the Americas than in Europe and the Middle East. The highest amount of Neanderthal ancestry is found in Oceania, but this is likely an artefact caused by some misclassified Denisovan segments. No prominent differences are observed between populations within the same geographic regions (Figure S13B). The intra-population variance is higher in Middle Eastern populations (especially Mozabite and Bedouin), likely reflecting recent admixture between sources with different levels of Neanderthal ancestry (e.g. African and West Eurasian). Denisovan segments are most abundant in Oceania (Figure S13A). They are also detectable at much lower levels in East Asia, the Americas and Central and South Asia, but negligible in Europe and the Middle East. Within Oceania, the Bougainville population has less Denisovan ancestry than the two populations from New Guinea (Figure S13B), consistent with dilution due to Southeast Asian admixture in the former.

All archaic segments in the "strict" set were pooled by geographical region to obtain maps of archaic ancestry frequencies along the genome. Figure S14 depicts the distribution of Neanderthal and Denisovan segments along chromosome 1 as an example.

(a) Distribution of Neanderthal segments along chromosome 1

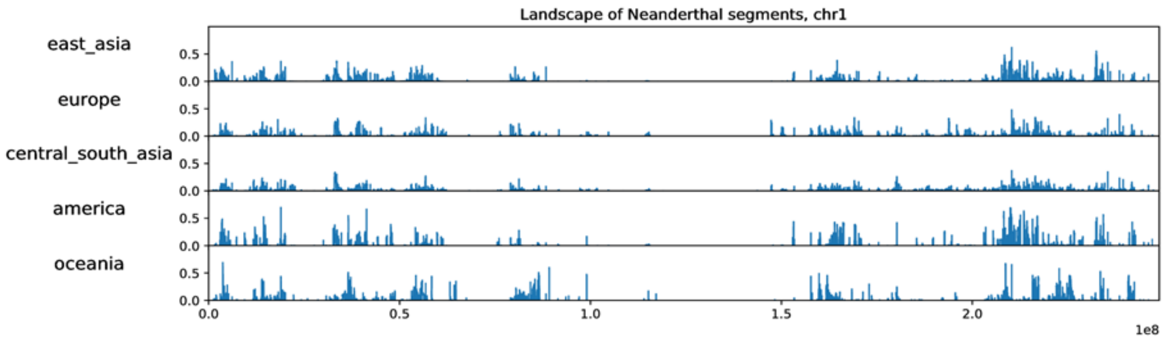


(b) Distribution of Denisova segments along chromosome 1

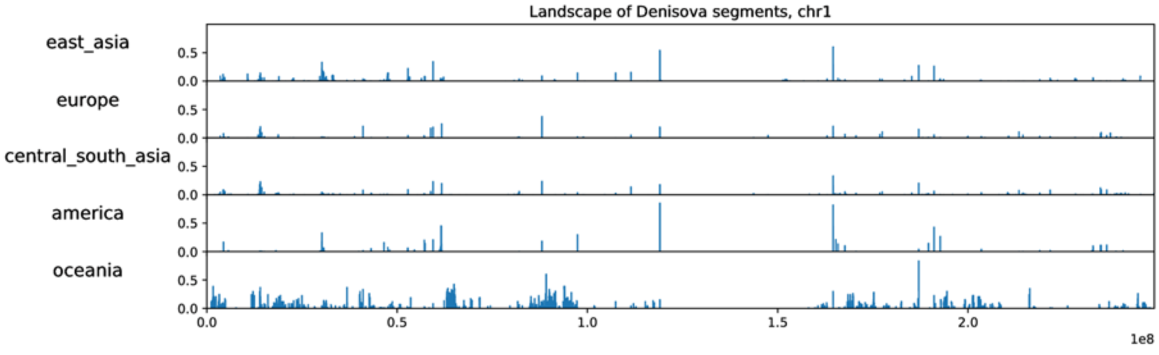


*Figure S14: Distribution of archaic segments ("strict" criteria) along chromosome 1 by geographical region.*

The difference between Oceania and other non-African populations appears more pronounced in the distribution of Denisovan than Neanderthal segments (Figure S14). To address this more formally, we quantified the length of overlapping genomic regions throughout the genome covered by at least two archaic segments between pairs of geographical regions, regardless of the genotypes in the segments (Table S6). $P(A|B)$ here denotes the probability that a genomic region being observed in geographical region $A$, conditioned on it being observed in geographical region $B$. The geographical structure is stronger in the genomic distribution of Denisovan segments with less overlapping between America, East Asia, Central/South Asia and Oceania. Denisovan segments might have been lost through genetic drift more often or sampled less frequently than Neanderthal segments because of their low frequency in most populations; however, despite similar amount of Neanderthal and Denisovan ancestry in Oceania, *P*(Oceania|non-Oceania) is also lower for Denisovan segments than for Neanderthal regions. This thus indicates that the local landscapes of Denisovan ancestry across the genome is less similar between Oceanian and Eurasian populations than what the Neanderthal landscapes are.

***Table S6: Intersection of genomic regions covered by at least two archaic segments between non-African populations****.* *Each value represents the probability of finding a genomic region in the column label conditioned on finding it in the row label.*

| *Neanderthal* | | | | | | | |
|---|---|---|---|---|---|---|---|
| **Geographic region** | **Total length (bp)** | **Conditional probability to also be found in** | | | | | |
| | | America | CS Asia | E Asia | Europe | Middle East | Oceania |
| America | 204886844 | - | 0.9117 | 0.9105 | 0.7235 | 0.6155 | 0.3579 |
| C&S Asia | 671828352 | 0.2780 | - | 0.6099 | 0.6458 | 0.6056 | 0.2409 |
| E Asia | 525811986 | 0.3548 | 0.7793 | - | 0.5513 | 0.4900 | 0.3000 |
| Europe | 482248609 | 0.3074 | 0.8997 | 0.6011 | - | 0.7981 | 0.2399 |
| Middle East | 453085016 | 0.2783 | 0.8979 | 0.5687 | 0.8495 | - | 0.2261 |
| Oceania | 218296694 | 0.3359 | 0.7413 | 0.7226 | 0.5300 | 0.4693 | - |

| *Denisovan* | | | | | | | |
|---|---|---|---|---|---|---|---|
| **Geographic region** | **Total length (bp)** | **Conditional probability to also be found in** | | | | | |
| | | America | CS Asia | E Asia | Europe | Middle East | Oceania |
| America | 13333796 | - | 0.5761 | 0.8382 | 0.3202 | 0.2470 | 0.2309 |
| C&S Asia | 55284712 | 0.1389 | - | 0.3665 | 0.2106 | 0.1773 | 0.1980 |
| E Asia | 56280344 | 0.1986 | 0.3600 | - | 0.1330 | 0.0852 | 0.1878 |
| Europe | 14067884 | 0.3035 | 0.8276 | 0.5321 | - | 0.6230 | 0.2146 |
| Middle East | 13893642 | 0.2370 | 0.7054 | 0.3451 | 0.6308 | - | 0.2091 |
| Oceania | 190235182 | 0.0162 | 0.0575 | 0.0556 | 0.0159 | 0.0153 | - |

*Divergence of archaic segments to archaic genomes*

All Neanderthal and Denisovan segments from the "strict" set in each genome were compared with the Altai Neanderthal, Vindija Neanderthal and Denisovan genomes. To recover archaic-private variants that were not present in files produced by merging variants-only modern VCFs with all-sites archaic VCFs, we assumed that all modern sites passing the strict mask but not present in the VCF files carried the reference allele; if these sites also pass the respective archaic mask and appear in the archaic all-sites files with alternative alleles, they also contribute to the counts of differences.

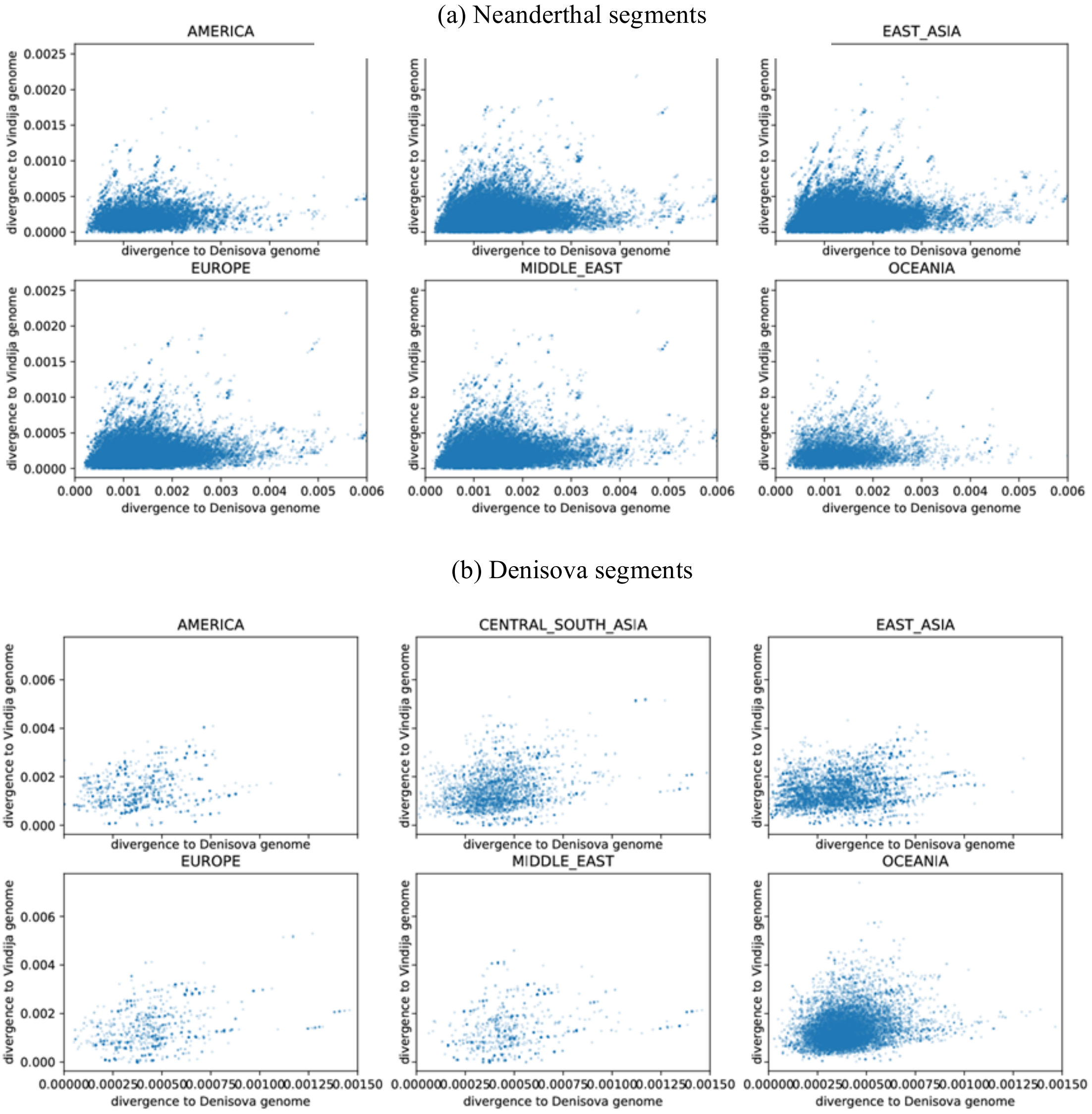


*Figure S15: Divergence of each identified archaic segment to the Vindija Neanderthal and Altai Denisovan genomes across geographical regions.*

The overall patterns of divergence to the Neanderthal genomes are almost identical across all six geographical regions (Figure S15A), while the patterns of divergence to the Denisovan genomes follow visibly different shapes in East Asia and Oceania: the points in Oceania form a well-defined, single cluster; in East Asia, the pattern appears more noisy, with additional segments displaying low divergence to the Altai Denisovan genome (divergence less than ~0.0001) that are absent from the Oceanian populations. This component is also potentially visible in America and Central/South Asia.

These results corroborate the finding from (*35*) of an additional pulse of Denisovan gene flow into East Asia. However, it is also worth noting that similarity in relation to known archaic genomes does not guarantee that the source is the same, since different source populations might show identical relationships to a given archaic individual. Based on evidence from nucleotide diversity and haplotype networks described below, we also find it plausible that some of the Denisovan ancestry in East Asian populations results from an additional admixture event separate from that in Oceanian populations. The structure in the East Asian patterns of Denisovan divergence (Figure S15B) might be due to partially overlapping distributions of divergence from the two components of admixture, or might possibly reflect an even more complicated history of admixture from several source populations at various locations and times.

#### *Nucleotide diversity within archaic segments*

Within one population, the expected number of nucleotide differences per site between two randomly drawn haplotypes is commonly known as nucleotide diversity ($\pi$). When comparing two populations, the same expected value between sequences randomly drawn from two populations (excluding all comparisons within the same population) is commonly known as absolute divergence ($D_{XY}$, also referred to as $\pi_{XY}$ , $\pi_B$ or $d_{XY}$ in the literature) (*68*). Based on the "strict" set of result, three sets of $D_{XY}$ between all pairs of populations were obtained: values calculated from only the Neanderthal segments in the genomes, from only the Denisovan segments in the genomes, and from only the unadmixed (also referred to as "modern" hereafter) segments of the genomes.

In this context, a "haplotype" refers to the collection of all Neanderthal (or Denisovan or modern) segments found in the same haploid genome. Since introgressed segments typically span different genomic regions in different individuals, it is only meaningful to compare nucleotide differences in the overlapping regions of two haplotypes (Figure S16). To limit computational costs when calculating $D_{XY}$, if the sample size of a population exceeds 10 individuals, only 20 haplotypes are randomly drawn for pairwise comparison with a maximum of 20 haplotypes from the other population (we found repeated draws produced very similar results).

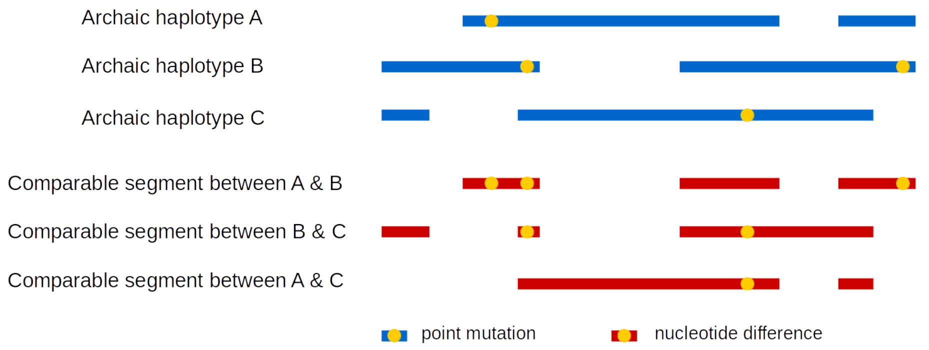


*Figure S16: Schematic showing comparable regions between three archaic haplotypes and nucleotide differences*

#### *Neanderthal vs. unadmixed regions*

The absolute divergence in Neanderthal regions ($D_{XY\text{-}N}$) and in unadmixed regions ($D_{XY\text{-}M}$) are colour-coded onto the upper-right and lower-left triangles, respectively, of a matrix in a heat map (Figure 6A). All $D_{XY\text{-}M}$ and $D_{XY\text{-}N}$ values were normalised such that variation within each is displayed using the same colour scale. A neighbour-joining tree was built with $D_{XY\text{-}M}$ values using San as outgroup (Figure S17A), and the populations in Figure 6A are ordered according to this tree. The heatmap is generally symmetrical: the pattern in Neanderthal segments largely mirrors that in unadmixed segments, forming major clusters separating Oceanian, American, East Asian and European-Central/South Asian populations. The neighbour-joining tree built from $D_{XY\text{-}N}$ (not shown) only differs from the unadmixed tree by two swappings of adjacent branches, between Xibo and Mongolian and between Tuscan and the French-Orcadian clade.

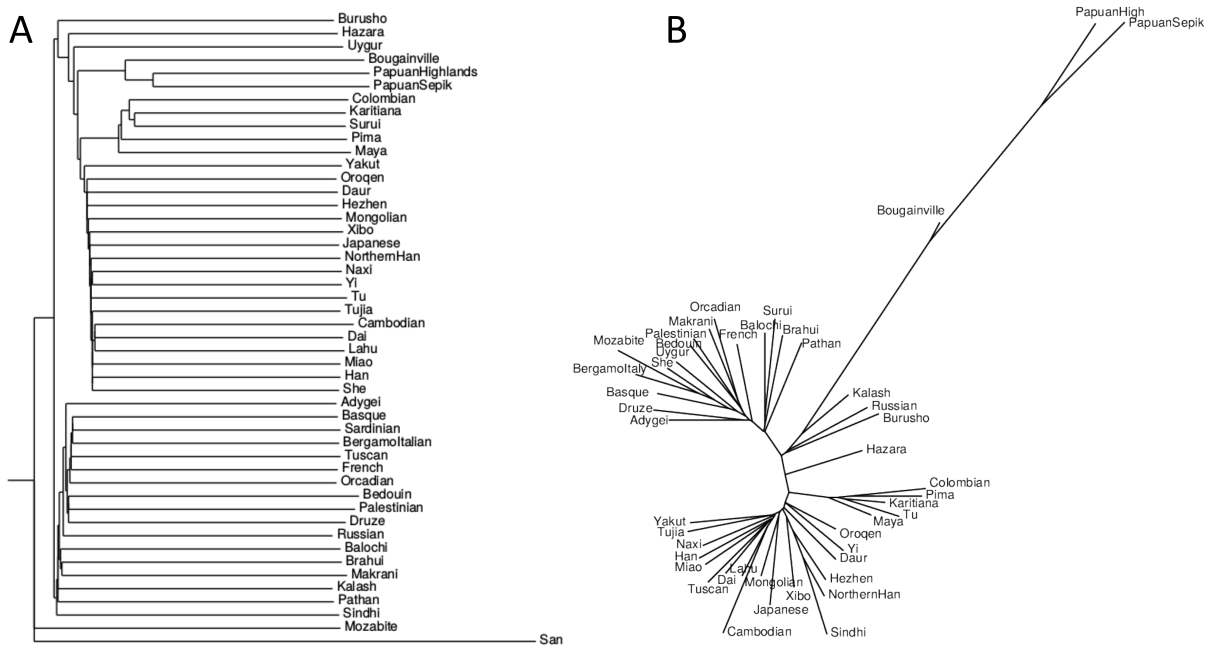


***Figure S17: Population relationships in different classes of genome segments.*** *A) Neighbour-joining tree built from* $D_{XY}$ *measured in unadmixed segments of the genome, rooted by San as outgroup. B) Unrooted neighbour-joining tree built from* $D_{XY}$ *measured in Denisovan segments of the genome.*

The symmetry is broken on finer scales. $D_{XY\text{-}N}$ between the North African Mozabite and European/Middle Eastern populations appears lower than that between some Central/South Asian populations and the latter; in fact, it is almost as low as comparisons within Europe. But in terms of $D_{XY\text{-}M}$ all Central/South Asian populations are closer to European/Middle Eastern ones than to Mozabite. Most likely this is because the sub-Saharan African ancestry in Mozabite increases $D_{XY\text{-}M}$ to Europe/Middle East, but the Neanderthal ancestry in Mozabite remains what was received from the same source as Europe/Middle East. The values in the cluster in the lower right corner, including populations from Europe, the Middle East and Central/South Asia (excluding three with high genetic affinity to East Asia), were normalized again excluding other populations (Figure S18). Now a European cluster can be distinguished, yet the relationships involving the Middle East and Central & South Asia are not well-defined. A history involving complex admixtures among the ancestors of these various groups, especially if including sources with no or very low levels of Neanderthal ancestry (including sub-Saharan Africans and the proposed basal Eurasian lineage (*34, 69*)), could have contributed to these patterns.

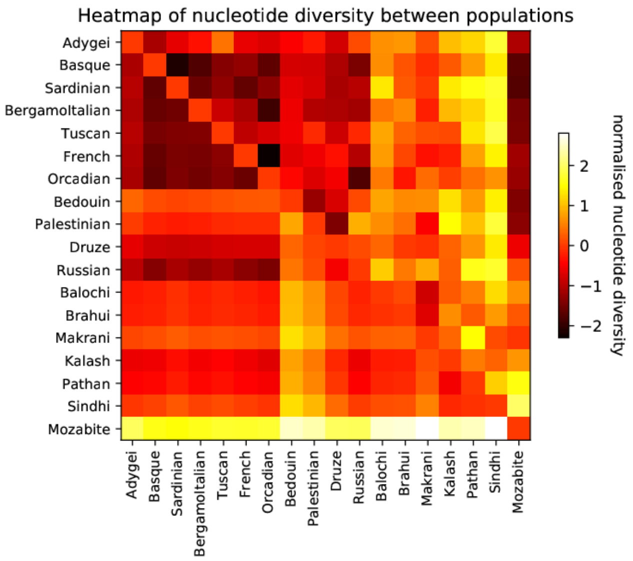


*Figure S18: Heat map comparing normalised $D_{XY}$ measured in Neanderthal vs. unadmixed regions of the genome, for West Eurasian populations only.*

*Denisovan vs. unadmixed regions*

Figure 6B shows the normalised absolute divergence in Denisovan segments ($D_{XY\text{-}D}$) and unadmixed regions ($D_{XY\text{-}M}$). The three Oceanian populations form a sister clade relative to all East Asian and American populations in unadmixed segments but exhibit very high divergence to all other populations in Denisovan segments. Their $D_{XY\text{-}D}$ to other non-African populations is largely homogenous, with only a faint affinity to East Asian populations. The Bougainville population displays lower $D_{XY\text{-}D}$ to East Asian populations than the other Oceanian populations, consistent with some Southeast Asian ancestry in Bougainville.

In Cambodians, the Denisovan segments show increased divergence to other East Asian populations, but decreased divergence to Oceanian populations in comparison to the unadmixed segments. In the tree built from $D_{XY-D}$ (Figure S17B), the Cambodian branch in also slightly longer in relation to other East Asian and American populations. Similarly to the much more prominent behavior of Oceanians, this evidence may suggest the presence of another component of Denisovan ancestry in Cambodians, with tentative connection to that in Oceania. One possibility is that this behavior might be driven by some fraction of South Asian related ancestry in Cambodians, which is likely not found in the other East Asian populations in the panel.

The $D_{XY\text{-}D}$ tree (Figure S17B) deviates considerably from the unadmixed tree (Figure S17A) even if we ignore structure within a given geographical region. The relatively low levels of Denisovan ancestry in most parts of the world likely add noise to these inter-population comparisons.

*Neanderthal vs. Denisovan regions*

Figure S19 directly compares intra- and inter-population divergence in Neanderthal and Denisova segments, again highlighting the distinct Denisovan ancestry in Oceania: in all comparisons excluding Oceanians, we again observe a strong correlation between $D_{XY\text{-}D}$ and $D_{XY\text{-}N}$, with the former typically smaller than the latter; in contrast, almost all comparisons including Oceanian populations show higher $D_{XY\text{-}D}$ than $D_{XY\text{-}N}$, and deviate from the otherwise largely linear relationship.

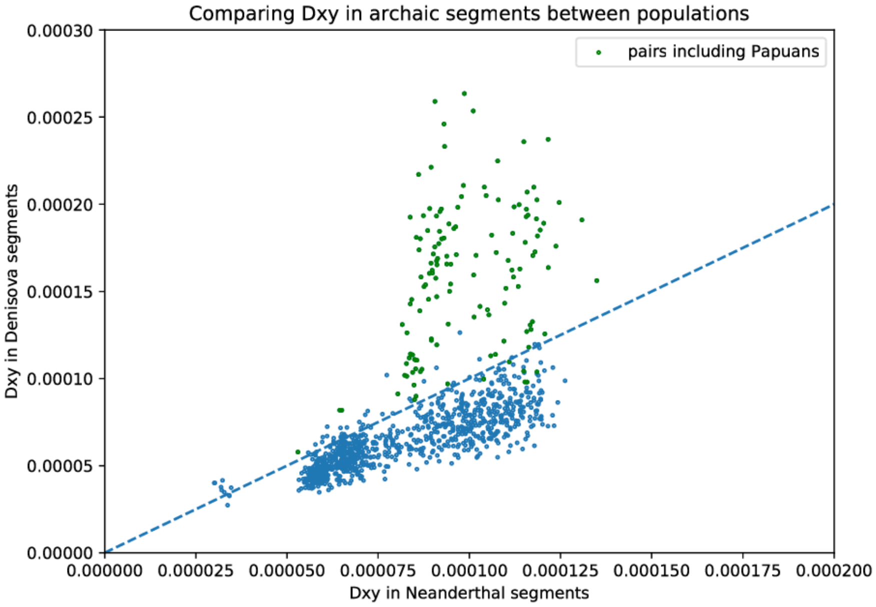


***Figure S19: Absolute divergence ($D_{XY}$) between all pairs of non-African populations measured in Denisovan and Neanderthal segments of the genome.***

#### *Archaic haplotype networks*

We attempted to reconstruct the relationships between all the archaic segments identified in different individuals in a given region of the genome. Assuming two archaic sequences descend from the same ancestral sequence at the time of admixture and a mutation rate of $1.25 \times 10^{-8}$ per site per generation, after 2,000 generations one would only expect to observe one difference per 20 kB. Long regions covered by as many archaic haplotypes as possible are therefore necessary to achieve a reasonable resolution. We searched for candidate regions in the genome by the following procedure:

- A multiple intersection of all Neanderthal/Denisovan segments (the "strict" set) on each chromosome was performed using multiIntersectBed from BEDTools (*55*) to obtain the total number of archaic haplotypes in a given genomic interval;
- The list of intervals was scanned to add new intervals by merging adjacent ones if a subset of individuals are present in both;
- A score is also assigned to each interval based on the length ($L$) and the number of samples ($n$):

$$s = n^w \cdot L$$

where $w$ can be tuned to adjust the weight of including more haplotypes over extending the genomic region; here we fixed it at 1, hence the score equals the total length of archaic sequences in the interval;

- Intervals shorter than 50 kB or with fewer than 5 occurrences were removed;
- Non-overlapping intervals with the highest scores were collected following a greedy algorithm: within a minimal candidate set of intervals that do not overlap with any other, the interval with the highest score is selected and moved to the selected set, and any other intervals that overlap with it removed from the candidate set; next to be

selected is the interval with the highest score among the remaining ones in the candidate set, and so on; the process repeats until no interval remains in the candidate set, and the algorithm moves on to the next set of intervals.

We constructed phylogenetic trees and haplotype networks using aligned archaic segments. When very few differences exist between haplotypes, haplotype networks have the advantage of allowing alternative links other than imposing a bifurcating tree with high uncertainty. The median joining network algorithm implemented in the pegas R package (*70*) was used to construct haplotype networks. Preliminary analysis also showed that the time to the most recent common ancestor (tMRCA) estimated from haplotype network analysis and maximum likelihood trees were highly correlated, although alternative links in the network tend to reduce tMRCA, especially when the sample size is large.

In each interval, polymorphic sites in all archaic haplotypes were retrieved to form a sequence alignment for haplotype network analysis. If a singleton allele among the archaic haplotypes was also present at a frequency above 1% in sub-Saharan African populations, we considered it was likely the result of phasing error and ignored it.

A total of 4,153 Neanderthal haplotype networks and 727 Denisovan ones were constructed using identified segments in all non-African genomes. A few examples are shown in Figure S20 and Figure S21. The sizes of the networks range from 3 to over 200 nodes. In some networks, haplotypes from the same geographical region cluster together (e.g. Figure S20A and Figure S21A), but in others identical haplotypes can be found across distant geographical regions (e.g. Figure S20B and Figure S21B). The Neanderthal haplotypes do not clearly tend to fall into separate clusters, as would be expected if there were multiple admixture events, whilst Denisovan haplotypes in Oceania are often separated from those in other geographical regions.

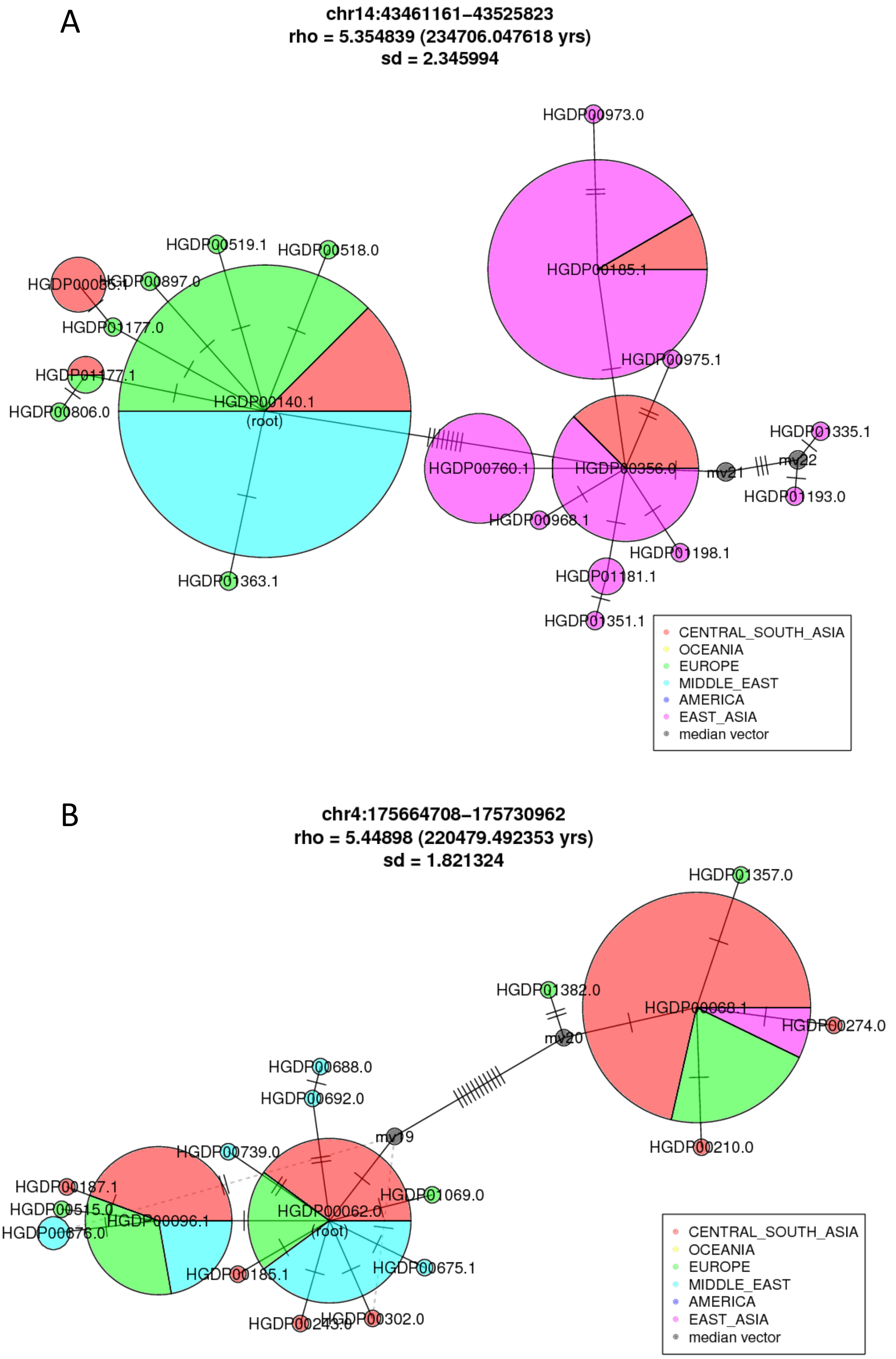


***Figure S20: Examples of Neanderthal haplotype networks.*** *Each circle represents a distinct haplotype, labelled by one sample name and coloured by the geographical origins of the samples, and the radius is proportional to the number of samples carrying that haplotype. The number of bars on the edges equals the number of mutations between haplotypes. Small grey circles labelled "mv" represents median vectors reconstructed in the median joining algorithm. Dashed lines: alternative links.*

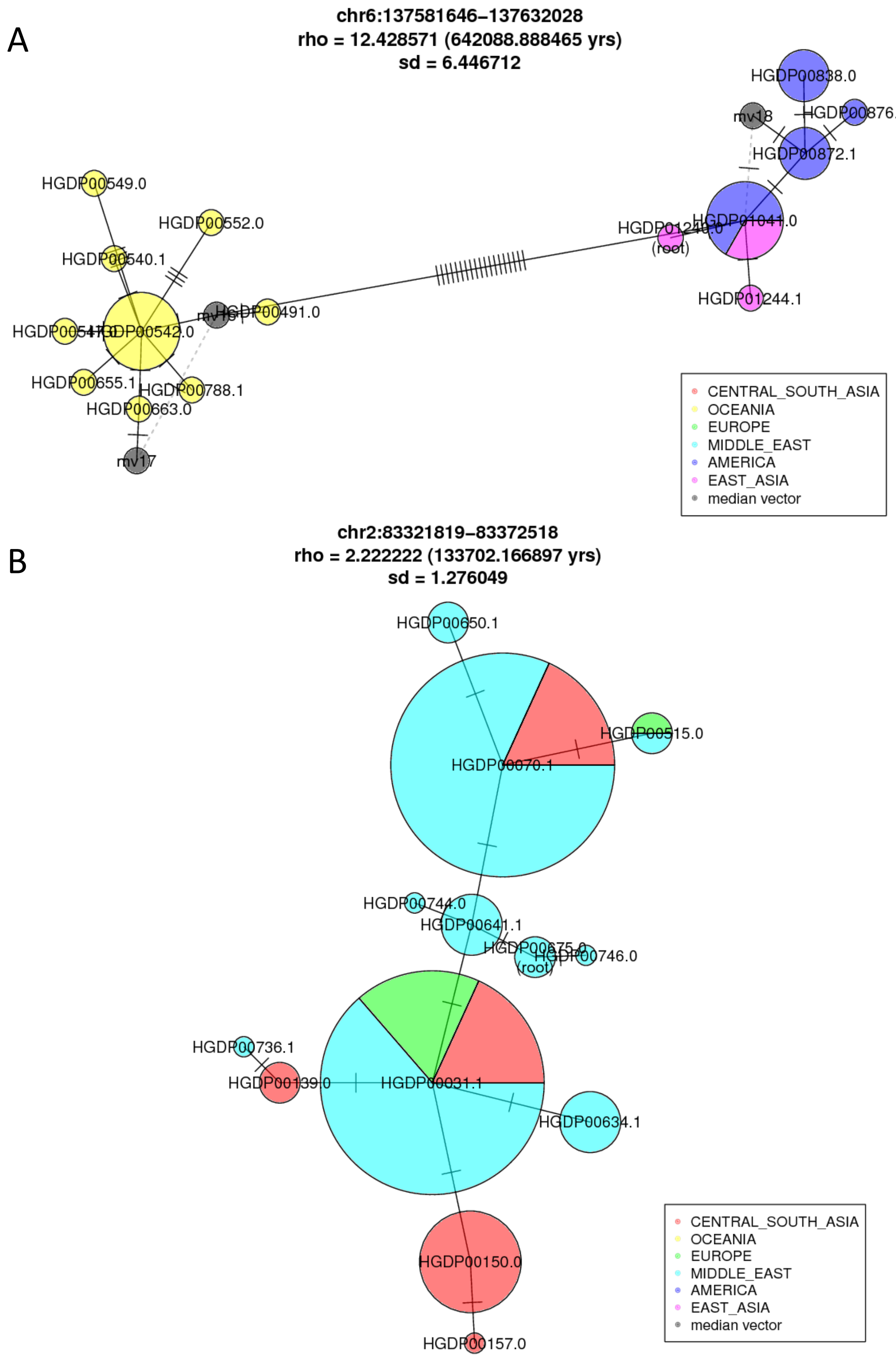


***Figure S21: Examples of Denisovan haplotype networks.*** *Each circle represents a distinct haplotype, labelled by one sample name and coloured by the geographical origins of the samples, and the radius is proportional to the number of samples carrying that haplotype. The number of bars on the edges equals the number of mutations between haplotypes. Small grey circles labelled "mv" represents median vectors reconstructed in the median joining algorithm. Dashed lines: alternative links.*

*Age of archaic haplotype networks*

To estimate the number of founding lineages contributing to extant haplotypes, we calculated the age ($\rho$) of each network (equivalent to tMRCA, or the height of a phylogenetic tree). The haplotype closest to the Vindija Neanderthal genome was assumed to be the root node. Following (*71*), $\rho$ is measured as the average shortest distance from all nodes to the root:

$$\rho = \frac{1}{n}\sum_{i=1}^{m} n_i l_i$$

and the variance:

$$\sigma^2 = \frac{1}{n^2}\sum_{i=1}^{m} n_i^2 l_i$$

where $n$ is the number of sequences, $m$ the total number of edges, and $n_i$ the number of samples whose shortest route to the root node passes through the $i$th edge. $\rho$ can then be converted into time in years with the mutation rate and the number of comparable sites in the genomic region.

Figure S22A shows the distribution of Neanderthal and Denisovan network ages in years. Filtering networks based on the length of the genomic region passing the mask, average *B* value of the genomic region, the number of missing sites in archaic genomes, the number of sites skipped (singletons also present in Africa), or the total number of polymorphic sites does not alter the shape of the distribution visibly; nor do these values exhibit distinct distributions between the groups of largest and smallest networks.

The age distribution from the two archaic sources are similar, both reaching the highest density below 50k years. The median of the Neanderthal and Denisovan haplotype networks are 55,613 and 55,070 years, respectively. However, in the tail of the distributions we also observe networks that are hundreds of thousands of years old.

We also constructed median joining networks on simulated haplotypes conditioned on the maximum number of introgressing haplotypes, using the simplified demographic history shown in Figure S22B. The sample sizes in Eurasia (including East Asia, Central/South Asia, the Middle East, and Europe), Oceania, and America populations were specified to match the geographical origin of actual Neanderthal haplotypes observed in each genomic region. The number of introgressing haplotypes was measured as the number of surviving lineages 2,000 generations ago, that is (backward in time) at the end of the bottleneck associated with admixture (which could also be an effect of negative selection). The duration and size of the bottleneck was arbitrarily selected to efficiently sample genealogies with few introgressing haplotypes. For each genomic region used to construct a haplotype network, coalescent trees were repeatedly simulated with a matching sample size, until the number of introgressing haplotypes became equal or less than the desired maximum. Genetic sequences of a length matched to the genomic region after filtering were then generated from the tree. In principle, the choices of bottleneck severity and ancestral Neanderthal size will influence the distribution of the coalescent trees retained; yet we found that in practice the effects on the age distribution was minimal.

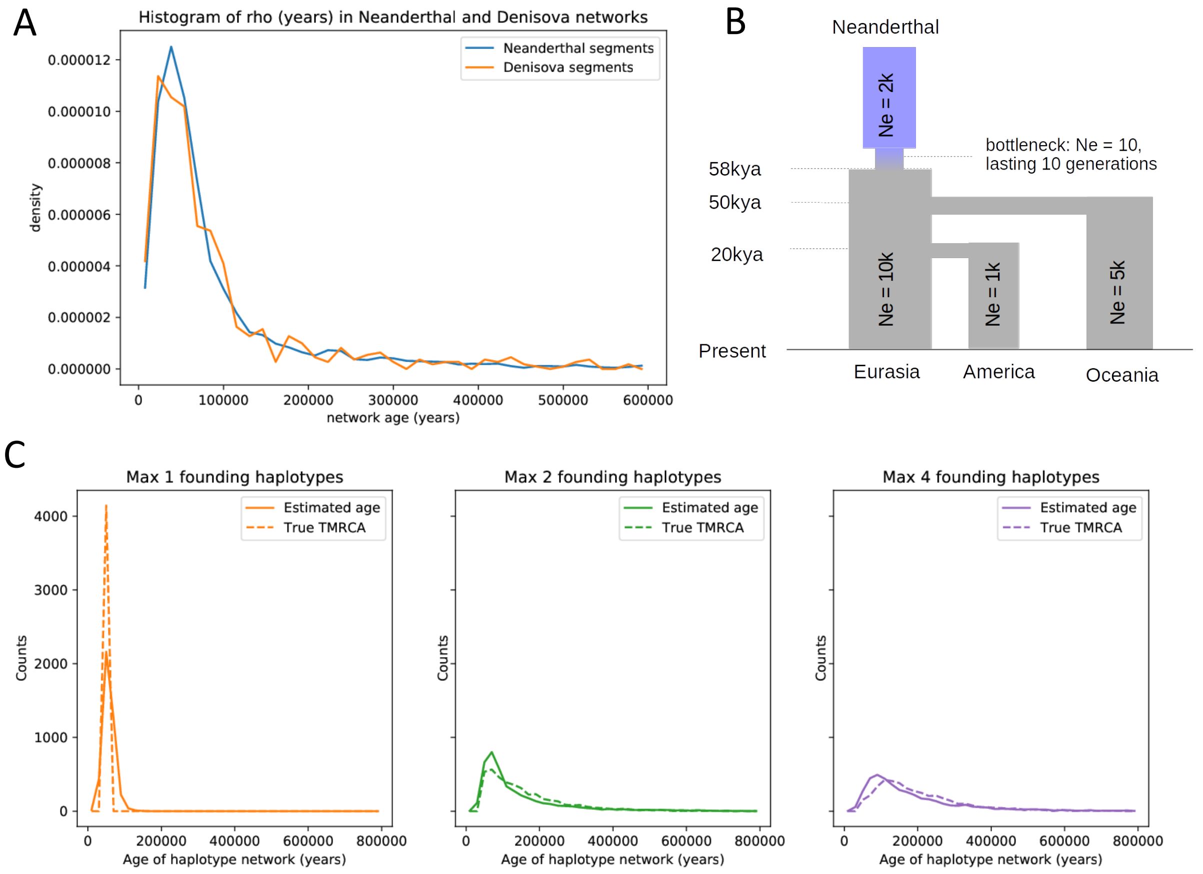


***Figure S22: Archaic haplotype network ages.*** *A) The distribution of Neanderthal and Denisovan haplotype network ages. B) Demographic model used in simulations exploring different numbers of founding Neanderthal haplotypes. C) Distribution of haplotype network ages estimated from simulations, and true tMRCA.*

Three sets of 4,153 genealogies with at most 1, 2 and 4 founding haplotypes respectively were obtained, to match the 4,153 genomic regions used in Neanderthal haplotype network analysis. The same algorithm used on the empirical data was used to build haplotype networks for each simulated alignment dataset and estimate $\rho$. The distribution of $\rho$ reasonably reflects the true tMRCA (Figure S22C). By allowing alternative links, the age estimated from haplotype networks can potentially underestimate the age of moderately old networks, but not the extremely old ones in the right tail of the distribution. The number of unique haplotypes in simulated and in empirical data align along the identity line with a strong correlation in all three sets of simulations, validating that the demographic model used in the simulations is a reasonable approximation of the true history.

Figure 6C compares the distribution of network ages estimated from empirical data and three sets of simulated data. The empirical distribution clearly differed from that produced in simulations with only one founding haplotype, yet its overall shape appears shifted towards the left in comparison to the curves produced in simulations with a maximum of two and four haplotypes. The shift could result from negative selection against archaic haplotypes or sub-population structure not implemented in the simulations. The simulations with a maximum of two and four haplotypes generated very similar distributions. As few as two founding haplotypes (i.e one individual) appear sufficient to produce the number of very old haplotype networks observed. A much larger number of Neanderthal individuals could still have been involved, contributing a reduced number of distinct haplotypes depending on the genetic diversity of the Neanderthal population. The combined effects of negative selection, genetic drift, dilution etc. could then have further reduced the diversity of the introgressed Neanderthal material to a very low level.

*Number of founding lineages*

The estimates of the ages of the haplotype networks reflect the genome-wide average number of founding archaic haplotypes. Here we estimate the number of founding archaic haplotypes in each genomic region as the number of surviving lineages in the tree at the time of admixture.

For each of the 4,135 genomic regions, a maximum likelihood tree was built from the sequence alignment and rooted by the haplotype closest to a San individual (HGDP00991). We chose to work with the tree structure rather than network here mainly because it more easily enables bootstrap analyses. The height at each node was determined by assigning height 0 to the tip farthest from the root, and positive heights to all other nodes, with the largest value at the root. Then the tree is truncated at the height corresponding to the expected number of differences per base pair in the time since admixture - assumed to be 2,000 generations - and the number lineages remaining connected to the root is counted. To gauge uncertainty, 1,000 non-parametric bootstrap replicates were performed for each genomic region.

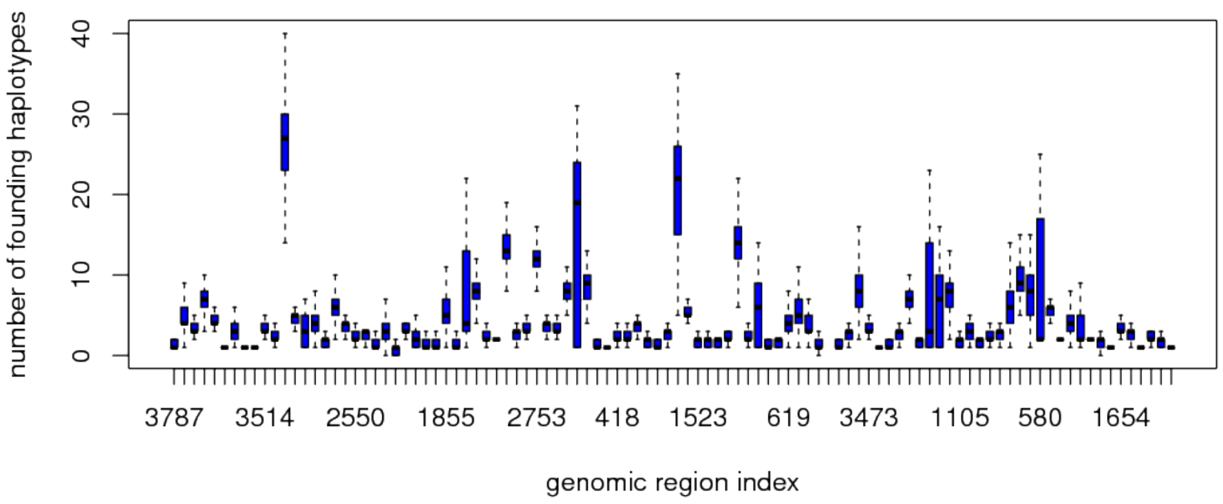


***Figure S23: The number of founding Neanderthal lineages across the genome.*** *A boxplot showing the number of founding Neanderthal lineages from 1,000 bootstraps in 100 genomic regions randomly selected from the total set of 4,135*

Figure S23 shows the number of founding haplotypes in 100 randomly sampled genomic regions, and Figure 6B shows the distribution of the mean number from all genomic regions. The estimated number of haplotypes were mostly low: in over 70% of the trees, the value of two standard deviations below the mean is lower than 2. However, there are also cases where more than 10 or even 20 lineages existed at the time of introgression. A total of 17 genomic regions were estimated to have more than 20 founding Neanderthal haplotypes (Table S7); their haplotype networks exhibit complicated structures radiating from one or two core haplotypes. It could mean that initially around a dozen (or more, depending on their relatedness) Neanderthal individuals contributed material, but except for very few regions of the genome, most Neanderthal lineages were subsequently lost through genetic drift and negative selection, so that only a handful of them remain among the diversity of Neanderthal segments in present-day modern humans.

*Table S7: Genomic regions with more than 20 founding Neanderthal haplotypes.*

| Chr | Start | End | #Haplotypes | #Unique haplotypes | Rho | sd | Rho (years) | sd (years) |
|---|---|---|---|---|---|---|---|---|
| 9 | 110937947 | 111009555 | 220 | 75 | 6.059 | 3.814 | 223447.6 | 140658.6 |
| 6 | 66848428 | 66915134 | 163 | 68 | 2.178 | 0.349 | 89369.29 | 14324.72 |
| 12 | 113841761 | 113933611 | 150 | 85 | 2.94 | 0.316 | 80828.57 | 8698.694 |
| 5 | 58495484 | 58599651 | 198 | 99 | 2.581 | 0.404 | 78918.59 | 12363.74 |
| 1 | 217246438 | 217327394 | 200 | 89 | 2.325 | 0.593 | 74454.43 | 18987.48 |
| 19 | 56087294 | 56138132 | 228 | 67 | 1.110 | 0.104 | 70633.69 | 6593.898 |
| 10 | 62941526 | 62993549 | 209 | 52 | 1.187 | 0.071 | 70207.82 | 4177.36 |
| 12 | 114080296 | 114130307 | 204 | 69 | 1.304 | 0.089 | 66588.11 | 4520.688 |
| 2 | 13827358 | 13919411 | 125 | 73 | 2.136 | 0.165 | 60264.87 | 4664.095 |
| 1 | 216557045 | 216647205 | 218 | 85 | 2.147 | 0.155 | 58608.5 | 4234.904 |
| 1 | 32911081 | 32992108 | 211 | 68 | 1.422 | 0.268 | 57411.51 | 10832.91 |
| 4 | 28482612 | 28545486 | 191 | 64 | 1.277 | 0.080 | 54515.13 | 3432.053 |
| 12 | 20849814 | 20933980 | 151 | 63 | 1.709 | 0.217 | 52310.35 | 6651.919 |
| 9 | 126565708 | 126646783 | 190 | 66 | 1.389 | 0.162 | 46613.15 | 5425.191 |
| 1 | 212466385 | 212548707 | 196 | 78 | 1.301 | 0.044 | 43196.05 | 1458.018 |
| 9 | 94515802 | 94565893 | 148 | 42 | 0.534 | 0.011 | 38530.75 | 781.0288 |
| 1 | 33403459 | 33454985 | 263 | 63 | 0.669 | 0.025 | 34713.19 | 1273.397 |

*Geographical separation*

Deep splits in a haplotype network could reflect multiple sources of archaic gene flow. To measure the divergence between geographic regions, we define two regions as separate in a network if none of the nodes containing haplotypes from one region has a closest neighbour containing haplotypes from the other region, and vice versa. Table S8 displays the total number of haplotype networks analyzed in this way (which need to contain at least two haplotypes from each region), and the number of networks showing separation between pairs of geographical regions.

The diversification among modern human populations will have caused some divergence in the archaic segments even if they descended from the same source of admixture. The proportion of completely geographically separated Neanderthal haplotype networks is largely consistent with our understanding of non-African population history. Much fewer Denisovan haplotype networks are available for comparison, yet there is a deep split between Oceania and all other non-African regions: networks are completely separated in almost all cases, most strikingly between Oceania and East Asia, as well as between Oceania and Central/South Asia. Fisher's exact test confirms that the distributions of fully separated Neanderthal and Denisovan networks are significantly different in these two comparisons (Table S9). Another pair that show a near-significant difference is Central/South Asia and East Asia, but in this case they are better connected in Denisovan networks than in Neanderthal networks. Overall, if we take the Neanderthal networks as representative of a single-source scenario, the strong geographical separation between Oceania and other regions in the Denisovan haplotype network provides further evidence for different source populations of Denisovan haplotypes in Oceanian and Eurasian populations.

*Table S8: The number of completely geographically separated haplotype network between pairs of regions out of the total number of comparable networks*

***Neanderthal haplotype networks, separated / total***

| | Central/South Asia | East Asia | Europe | Middle East | Oceania |
|---|---|---|---|---|---|
| America | 254 / 862 | 184 / 878 | 221 / 618 | 245 / 506 | 159 / 263 |
| Central/South Asia | - | 228 / 2064 | 133 / 2190 | 139 / 2032 | 261 / 690 |
| Eeast Asia | - | - | 338 / 1356 | 429 / 1162 | 187 / 714 |
| Europe | - | - | - | 81 / 1932 | 249 / 448 |
| Middle East | - | - | - | - | 230 / 393 |

***Denisova haplotype networks, separated / total***

| | Central/South Asia | East Asia | Europe | Middle East | Oceania |
|---|---|---|---|---|---|
| America | 6 / 36 | 8 / 59 | 4 / 15 | 3 / 9 | 10 / 12 |
| Central/South Asia | - | 4 / 88 | 3 / 42 | 3 / 32 | 37 / 39 |
| East Asia | - | - | 5 / 27 | 5 / 14 | 35 / 40 |
| Europe | - | - | - | 2 / 31 | 7 / 9 |
| Middle East | - | - | - | - | 6 / 8 |

It is notable that in the only two networks where Denisovan haplotypes in Oceania are not fully separated from those in Central/South Asia, Oceania and East Asia are also connected. Out of the five networks where Oceania and East Asia are not separated, three involve Denisovan haplotypes from Cambodia as a bridge: in two cases the Cambodian haplotype is the sole connection, in another one a haplotype from Lahu is also involved. One possibility is that Cambodian ancestry contains a component that is somewhat intermediate between or similarly related to Oceanian and East Asian ancestries (e.g. South Asian related). But since this connection to Oceania is not observed in unadmixed regions of the genome (Figure 6B), it could also possibly suggest another independent component of Denisovan ancestry in Cambodians, whose source population is closer to the source in Oceania.

*Table S9: p-values from Fisher's exact test on different distributions of separated/connected networks in Neanderthal vs. Denisovan haplotypes between pairs of regions*

| | Central/South Asia | East Asia | Europe | Middle East | Oceania |
|---|---|---|---|---|---|
| America | 0.1320 | 0.2419 | 0.5908 | 0.5067 | 0.1375 |
| Central&South Asia | - | 0.0534 | 0.7398 | 0.4802 | $3.075 \times 10^{-13}$ |
| East Asia | - | - | 0.6523 | 1 | $5.207 \times 10^{-15}$ |
| Europe | - | - | - | 0.3801 | 0.3100 |
| Middle East | - | - | - | - | 0.4793 |
